# Chemical genetics strategy to profile kinase target engagement reveals role of FES in neutrophil phagocytosis via SYK activation

**DOI:** 10.1101/841189

**Authors:** Tom van der Wel, Riet Hilhorst, Hans den Dulk, Tim van den Hooven, Nienke M. Prins, Joost A.P.M. Wijnakker, Bogdan I. Florea, Eelke B. Lenselink, Gerard J.P. van Westen, Rob Ruijtenbeek, Herman S. Overkleeft, Allard Kaptein, Tjeerd Barf, Mario van der Stelt

## Abstract

Chemical tools and methods that report on target engagement of endogenously expressed protein kinases by small molecules in human cells are highly desirable. Here, we describe a chemical genetics strategy that allows the study of non-receptor tyrosine kinase FES, a promising therapeutic target for cancer and immune disorders. Precise gene editing was used in combination with a rationally designed, complementary fluorescent probe to visualize endogenous FES kinase in HL-60 cells. We replaced a single oxygen atom by a sulphur in a serine residue at the DFG-1 position of the ATP-binding pocket in an endogenously expressed kinase, thereby sensitizing the engineered protein towards covalent labeling and inactivation by a fluorescent probe. The temporal control offered by this strategy allows acute inactivation of FES activity both during myeloid differentiation and in terminally differentiated neutrophils. Our results show that FES activity is dispensable for differentiation of HL-60 cells towards macrophages. Instead, FES plays a key role in neutrophil phagocytosis by activation of SYK kinase, a central regulator of immune function in neutrophils. This strategy holds promise as a target validation method for kinases.

## Introduction

Protein kinases comprise a 518-membered family of enzymes that play an essential role in intracellular signaling processes. They transfer a phosphate group from ATP to specific amino acid residues in proteins, thereby modulating protein activity, localization and protein-protein interactions.^1,2^ Protein kinases are involved in many cellular functions, including proliferation, differentiation, migration and host-pathogen interactions. Kinases are also an important class of drug targets for the treatment of cancer^3^. However, current FDA-approved kinase inhibitors are designed to target only <5% of the entire kinome^4^ and therapeutic indications outside oncology are vastly underrepresented.^5,6^ These so-far untargeted kinases thus offer great opportunities for the development of novel molecular therapies for various diseases. The non-receptor tyrosine kinase feline sarcoma oncogene (FES), subject of the here presented study, is a potential therapeutic target for cancer and immune disorders.^7–9^

FES, together with FES-related kinase (FER), constitutes a distinct subgroup within the family of tyrosine kinases, defined by their unique structural organization. FES is able to form oligomers via its F-Bin-Amphiphysin-Rvs (F-BAR) domain, which drives translocation from the cytosol to the cell membrane.^10,11^ In addition, FES possesses a Src Homology 2 (SH2) domain that binds phosphorylated tyrosine residues and thereby functions as protein interaction domain.^12,13^ The catalytic domain of FES is located on its C-terminal end. Phosphorylation of Y713 in the activation loop of FES is a prerequisite for its kinase activity and can occur either via autophosphorylation or phosphorylation by Src family kinases.^14,15^ FES has a restricted expression pattern and is found in neuronal, endothelial and epithelial cells. Its expression levels are highest in cells of hematopoietic origin, especially those in the myeloid lineage.^16^ Most of the physiological processes of FES have been studied in macrophages^17,18^ or mast cells^19^, but far less is known about its role in other terminally differentiated myeloid cells, such as neutrophils.

The successful development of new kinase-targeting drugs strongly depends on our understanding of their underlying molecular and cellular mechanism of action, *i.e.* the preclinical target validation.^20^ The physiological function of many kinases remains, however, poorly characterized and their direct protein substrates are often unknown. Genetic models (congenital deletion or expression of kinase-dead variants) may be used to study these questions. For example, FES knock-out animals revealed a role of FES in myeloid differentiation^21^, leukocyte migration^22,23^ and the release of inflammatory mediators.^17^ However, long-term, constitutive genetic disruption of kinases can result in compensatory mechanisms that counteract defects in cellular signaling. For example, mice lacking both FES and FER have more pronounced defects in hematopoiesis than counterparts lacking only one of these kinases, which may indicate that these related kinases may compensate for each other’s loss.^24^ Long-term, permanent genetic models are therefore poorly suited to study rapid and dynamic signaling processes.^25,26^ In addition, phenotypic differences between independently generated knock-out animals are not uncommon, as was the case for two independently developed *fes*^−/−^ mice.^17,27^ A complementary approach is the use of small molecule inhibitors to modulate kinase activity in an acute and temporal fashion. This approach more closely resembles therapeutic intervention, but the available pharmacological tools, especially for non-validated kinases, often suffer from a lack of selectivity.^25,28^ Currently, there are no suitable FES inhibitors available for target validation studies, because they either lack potency or selectivity, and all cross-react with FER.^7,29^

A key step in the target validation process consists of obtaining proof of target engagement, which is essential to correlate inhibitor exposure at the site of action to a pharmacological and phenotypic readout.^30^ Information about kinase engagement is also useful for determining the dose required for full target occupancy without inducing undesired off-target activity.^31^ Chemical probes that make use of a covalent, irreversible mode of action are ideally suited to study target engagement.^30^ Incorporation of reporter tags enable target visualization (*e.g.* fluorophores) or target enrichment and identification (*e.g.* biotin). In the field of kinases, reported chemical probes either target a conserved active site lysine residue in a non-selective fashion^32^ or non-catalytic cysteine residues in the ATP binding pocket.^33,34^ The first class of kinase probes lacks the selectivity required for cellular target engagement studies. On the other hand, the majority of kinases, including FES, do not possess targetable cysteine residues in the catalytic pocket.^35^

Garske *et al.* previously introduced the elegant concept of covalent complementarity: the use of an engineered kinase in which the gatekeeper amino acid residue is mutated into a cysteine, combined with electrophilic ATP analogs to study target engagement.^36^ Other positions in the kinase active site have also been investigated^37–39^, but secondary mutations were required to improve cysteine reactivity or compound selectivity and potency. Of note, all studies relied on transient or stable overexpression of the mutant kinase, rather than physiological model systems with endogenous expression levels. Since overexpression of kinases is known to disrupt physiological intracellular signaling cascades^40^, there is a need for target validation methods that visualize specific endogenous kinase activity and its engagement by small molecules without perturbing normal cellular processes.

Inspired by these established and emerging concepts, we describe herein a chemical genetics strategy to profile acute target engagement of FES kinase by a complementary, mutant-specific chemical probe. To the best of our knowledge, this is the first report in which precise gene editing is used in combination with a rationally designed, complementary fluorescent probe to visualize a specific endogenous kinase in human cells. We replaced a single oxygen atom by a sulphur in a serine residue at the DFG-1 position in the ATP-binding pocket of endogenously expressed FES in HL-60 cells, thereby sensitizing the engineered kinase towards covalent labeling by the fluorescent probe. The use of mutant and wild-type cells to account for on-target and off-target effects respectively, allowed us to demonstrate that FES activity is dispensable for myeloid differentiation of HL-60 cells along the macrophage lineage. Instead, FES was shown to play a key role in phagocytosis of bacteria by neutrophils, via activation of SYK kinase, a central regulator of immune function in this cell type. Altogether, these results demonstrate that the presented chemical genetics strategy holds promise for target validation studies in kinase drug discovery.

## Results

### Chemical genetics strategy: precise gene editing combined with complementary, covalent probes

The key feature of the chemical genetics strategy is the combination of an engineered, mutant kinase with the design and application of complementary, covalent inhibitors (Figure 1A). The ATP-binding pocket of the kinase of interest is sensitized towards pharmacological inactivation using complementary probes by replacing an amino acid for a cysteine residue. The mutant is biochemically characterized to verify that the cysteine point mutation minimally affects kinase function (Figure 1B, step 1). Subsequently, complementary electrophilic inhibitors are designed to covalently react with this cysteine (Figure 1B, step 2). Covalent, irreversible inhibitors can have several advantages over reversible compounds, such as sustained target occupancy, lower susceptibility to competition by high intracellular ATP concentrations and a pharmacodynamic profile that is dependent on the target’s *de novo* protein synthesis rate.^41^ The inhibitor will have far lower potency on the wild-type kinase, which does not possess a nucleophilic residue in its active site, thereby making the inhibitor mutant-specific. An important feature of our strategy is that the corresponding cysteine point mutation is introduced endogenously in a relevant cell line using CRISPR/Cas9-mediated gene editing (Figure 1B, step 3). This circumvents the use of cellular overexpression systems, which may disturb the intrinsic balance of the kinase signaling network, leading to compensatory effects and other artefacts in cellular function.^40^ Importantly, the covalent binding mode of the inhibitor enables target engagement profiling by acting as a chemical probe and its ligation handle can be further functionalized with reporter tags for visualization by SDS-PAGE (fluorophore) or identification of the bound targets by mass spectrometry (biotin) (Figure 1B, step 4). Since this approach allows acute inactivation of kinases with high specificity, it can be employed to study kinase function and thereby aid in its validation as therapeutic target (Figure 1B, step 5).

**Figure 1.**
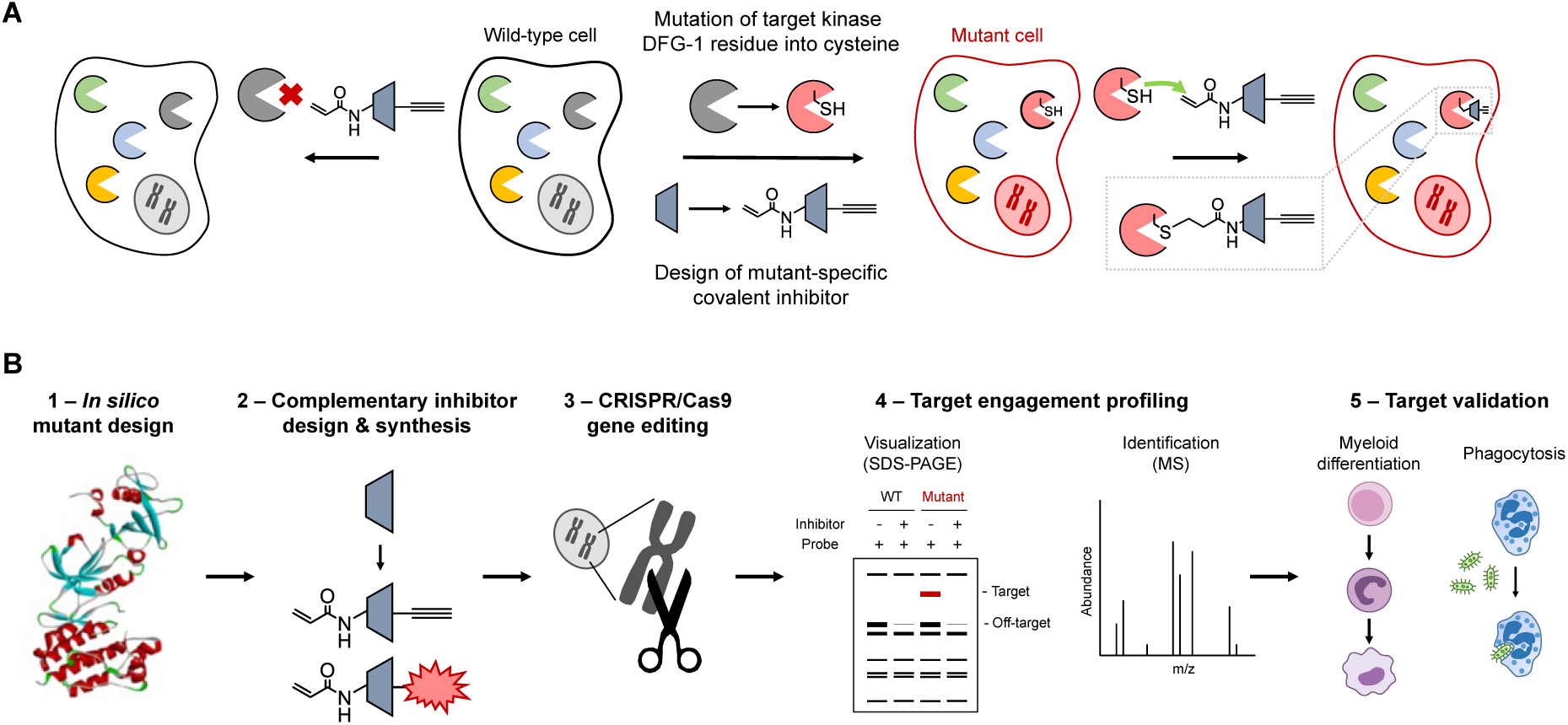
Chemical genetics strategy and workflow to visualize kinase activity and target engagement. **(A)** General strategy involving mutagenesis of a kinase DFG-1 residue into a nucleophilic cysteine accompanied by mutant-specific covalent inhibitor design. A non-selective, reversible inhibitor is modified with an acrylamide electrophile to covalently react to the introduced cysteine, along with an alkyne ligation handle for (bioorthogonal) conjugation to reporter tags using click chemistry. The mutation is introduced in a relevant cell line using CRISPR/Cas9 gene editing in order to study the kinase in an endogenous expression system. **(B)** Schematic workflow for application of mutant-specific, complementary probes in target engagement and target validation studies.

### Step 1 - Biochemical characterization of engineered FES kinases

To introduce a cysteine residue at an appropriate position in the ATP-binding pocket of FES, we inspected the reported crystal structure of FES with reversible inhibitor TAE684 (compound **1**) (PDB: 4e93).^29^ We selected nine active site residues situated in proximity of the bound ligand (Figure 2A) and generated the respective cysteine point mutants by site-directed mutagenesis on truncated human FES (SH2 and kinase domain, residues 448-822) fused to a N-terminal His-tag. The wild-type protein and the mutants were recombinantly expressed in *Escherichia coli* (*E. coli*), purified using Ni^2+^-affinity chromatography and tested for catalytic activity using a time-resolved fluorescence resonance energy transfer (TR-FRET) assay (Figure 2B). Four of the nine tested mutants did not display any catalytic activity, including G570C (located on P-loop) and G642C (hinge region). Three mutants near the kinase hydrophobic backpocket (I567C, V575C and L638C) retained partial activity, whereas only two mutants (T646C and S700C) displayed catalytic activity similar to wild-type FES. Our attention was particularly drawn to the S700C mutant, which involves the residue adjacent to the highly conserved DFG motif (DFG-1). Since several other kinases (e.g. MAPK1/3, RSK1-4and TAK1) express an endogenous cysteine at DFG-1 that can be targeted by electrophilic traps^33^, we chose to profile FES^S700C^ in more detail. The engineered kinase displayed identical reaction progress kinetics (Figure 2C) and similar affinity for ATP (K_M_ = 1.9 μM for FES^WT^ and K_M_ = 0.79 μM for FES^S700C^; Figures 2D and S1A).

**Figure 2.**
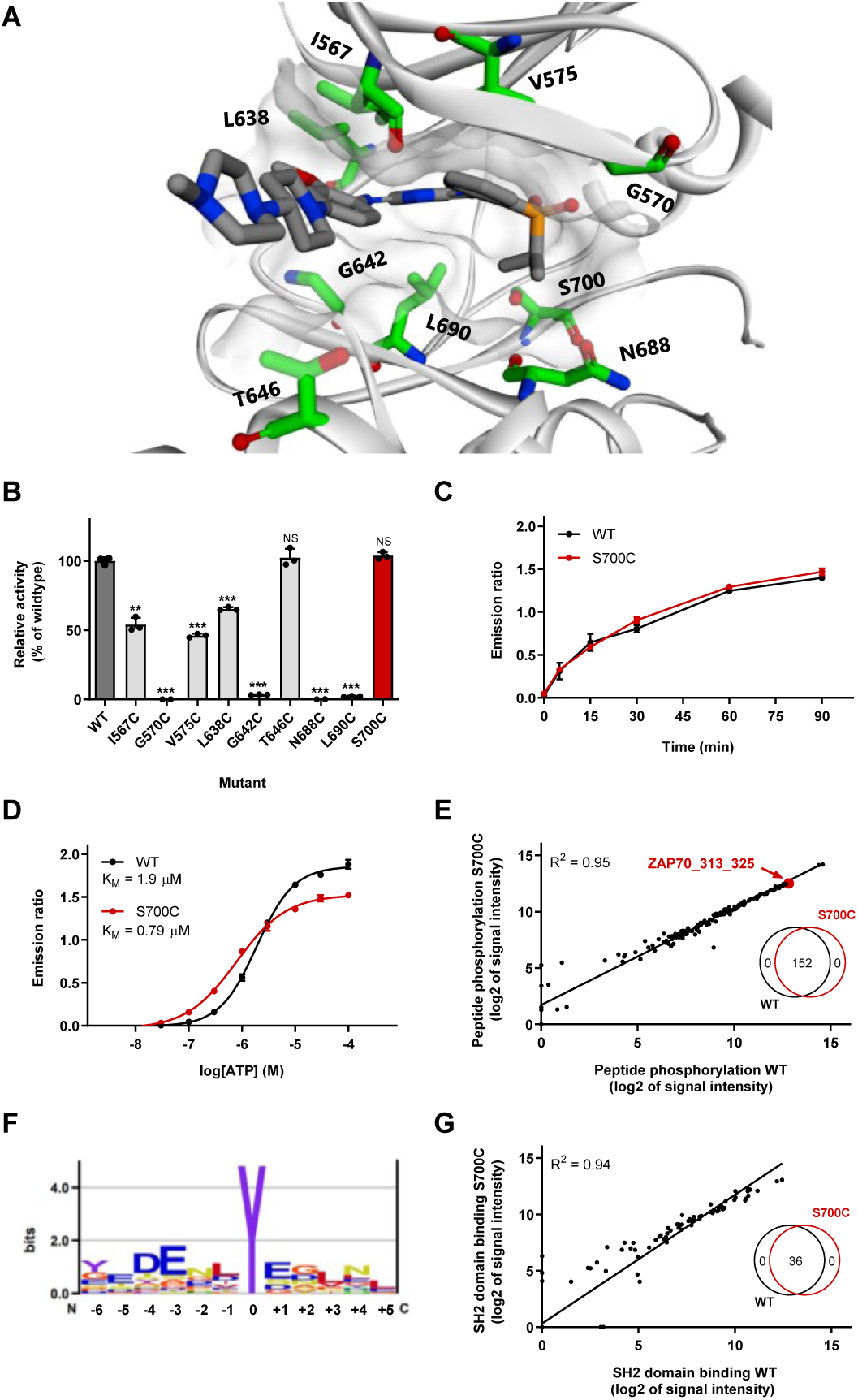
Design and characterization of FES cysteine point mutants. **(A)** Location of mutated active site residues in FES crystal structure with bound reversible inhibitor TAE684 (PDB code: 4e93). **(B)** Activity of recombinantly expressed FES mutants compared to wild-type, determined as relative amount of phosphorylated peptide substrate after 60 min incubation using TR-FRET assay. **(C)** Reaction progress kinetics for FES^WT^ and FES^S700C^. **(D)** Determination of ATP K_M_ for FES^WT^ and FES^S700C^. Enzyme reactions in TR-FRET assay were performed with U*Light*-TK peptide (50 nM) and ATP (**B**, **C**: 100 µM, **D**: variable) and quenched (**B**, **D**: after 60 min, **C**: variable). **(E)** Peptide phosphorylation substrate profile for FES^WT^ and FES^S700C^ as determined in PamChip® microarray. Peptides were filtered for those with ATP-dependent signal and log2 of signal intensity >3. The peptide substrates were identical for FES^WT^ and FES^S700C^ (Venn diagram, inset). **(F)** Preferred substrate consensus sequence based on FES^WT^ substrate profile. Illustration was rendered using Enologos (http://www.benoslab.pitt.edu). **(G)** SH2 domain binding profile for FES^WT^ and FES^S700C^ as determined in PamChip® microarray. Peptides with non-specific antibody binding were excluded. The peptide SH2 binding partners were identical for FES^WT^ and FES^S700C^ (Venn diagram, inset). Data represent means ± SEM (n = 3). Statistical analysis: ANOVA with Holm-Sidak’s multiple comparisons correction: *** *P* < 0.001; NS if *P* > 0.05.

To assess whether the introduced mutation affected substrate recognition, a comparative substrate profiling assay was performed using the PamChip^®^ microarray technology. This assay is based on the phosphorylation of immobilized peptides by purified FES and detection using a fluorescently labeled anti-phosphotyrosine antibody. Strikingly, the substrate profiles of FES^WT^ and FES^S700C^ were completely identical (Figure 2E, inset; Table S1), indicating that the S700C mutation did not affect substrate recognition. Moreover, the absolute peptide phosphorylation levels showed a strong correlation (R^2^ = 0.95). Sequence analysis of the top 30 of highest signal peptides revealed that FES prefers negatively charged substrates with hydrophobic residues at positions −1 and +3 and acidic residues at position −4, −3 and +1 relative to the tyrosine phosphorylation site (Figure 2F). These results are in line with a previous study that reported on FES substrate recognition.^42^ Lastly, a modified PamChip array to measure phosphopeptide binding to the Src homology 2 (SH2) domain of FES^WT^ and FES^S700C^ showed that the introduced mutation did not affect the SH2 binding profile (Figure 2G and Table S2). In short, the DFG-1 residue (Ser700) in the ATP-binding pocket of FES was identified as an excellent position to mutate into a nucleophilic cysteine, without affecting FES kinase activity, kinetics, substrate recognition or SH2 binding profile.

### Step 2 - Design, synthesis and characterization of complementary probes for FES^S700C^

The reversible ligand TAE684 was used as a starting point to develop a complementary probe for FES^S700C^. To assess whether the FES^S700C^ was still sensitive to inhibition by TAE684, the protein was incubated with various concentrations of TAE684 and its half maximum inhibitory concentration (expressed as pIC_50_) was determined. We found that TAE684 was a potent inhibitor both on FES^WT^ and FES^S700C^ with pIC_50_ values of 8.1 ± 0.04 and 9.0 ± 0.02, respectively (Table 1). According to the co-crystal structure of FES^WT^ with TAE684 (PDB: 4e93) the isopropyl sulfone moiety is in the close proximity of the Ser700 residue at the DFG-1 position. Therefore, several derivatives of TAE684 were synthesized, in which the R_2_−phenyl ring was substituted with an acrylamide group as electrophilic warhead (Scheme S1-4). The acrylamide is hypothesized to covalently interact with the engineered cysteine, but not with the serine of the wild-type protein. Since the strategy aims at exclusively inhibiting mutant but not wild-type FES, we also decided to remove the piperidine-piperazine group as it is known to form water-mediated hydrogen bonds with hinge region residues and contribute to ligand affinity.29 The pIC_50_ values of compound **2-6** are listed in Table 1 (dose-response curves in Figure S1B-F). Removal of the piperidine-piperazine group (compound **2**) resulted in a modest reduction in potency on FES^WT^ and FES^S700C^. Introduction of an acrylamide at the *meta*-position of the phenyl ring (compound **3**) further decreased affinity for FES^WT^, but also led to significant loss in activity on the FES^S700C^ mutant. Moving the acrylamide to the *ortho*-position resulted in compound **4**, which exhibited excellent potency on FES^S700C^ (pIC_50_ = 8.4 ± 0.03), whereas a major reduction in potency on FES^WT^ (pIC_50_ = 5.7 ± 0.21) was found, resulting in an apparent selectivity window of 238-fold. We substituted the acrylamide with a propionyl amide (compound **5**) to confirm its important role in binding to FES^S700C^. In line with the proposed mode of action, compound **5** displayed low inhibitory potency on FES^S700C^.

**Table 1.**
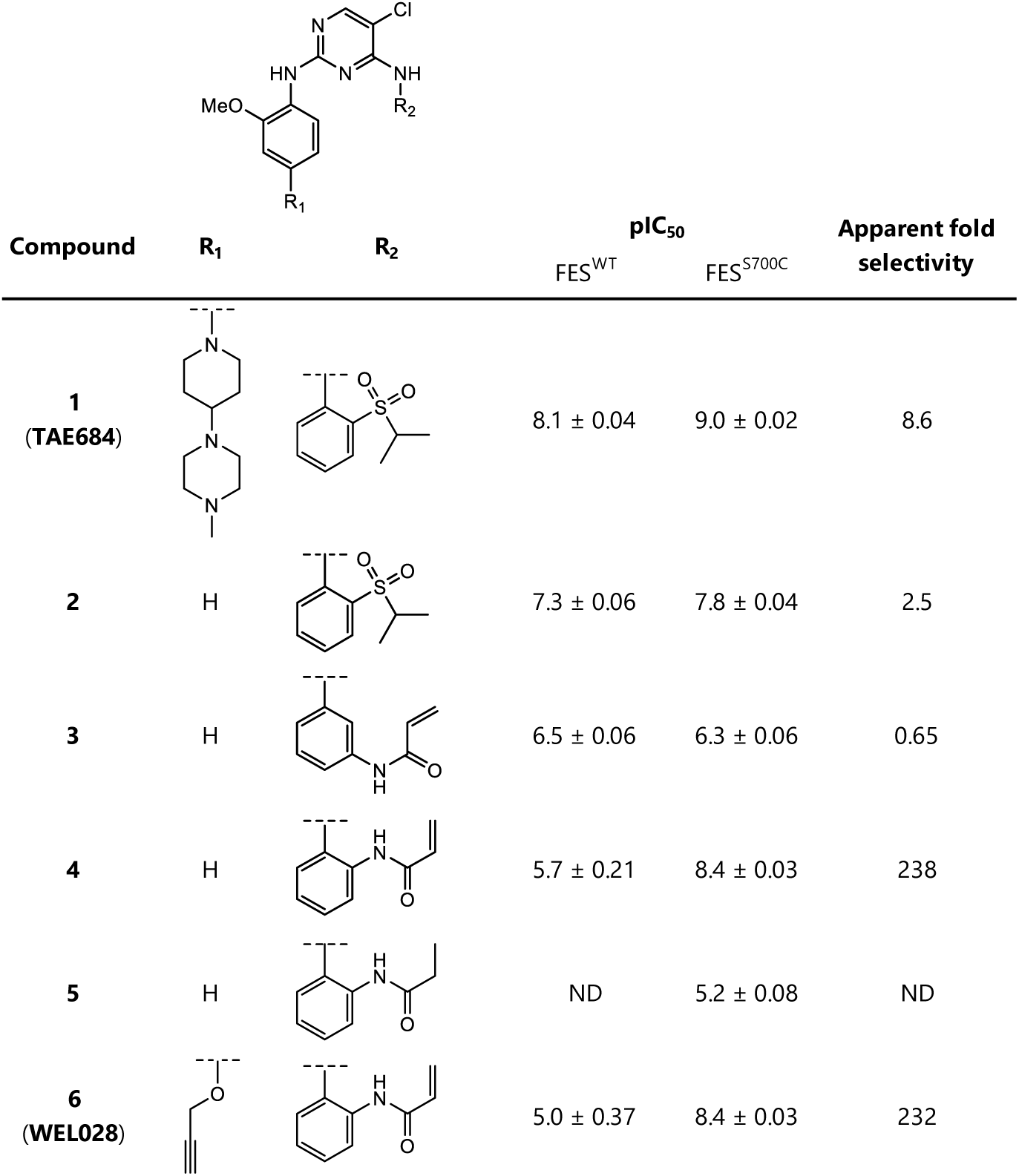
Inhibitory potency of synthesized TAE684 derivatives against FES^WT^ and FES^S700C^. Half maximal inhibitory concentrations (expressed as pIC_50_) determined on recombinantly expressed FES^WT^ and FES^S700C^ in a TR-FRET assay. Apparent fold selectivity was calculated as IC_50_ on FES^WT^ divided by IC_50_ on FES^S700C^. Data represent means ± SD; n = 3. ND: not determined. Complete dose-response curves can be found in Figure S1.

| Compound | R<sub>1</sub> | R<sub>2</sub> | pIC<sub>50</sub> |  | Apparent fold selectivity |
| --- | --- | --- | --- | --- | --- |
|  |  |  | FES<sup>WT</sup> | FES<sup>S700C</sup> |  |
| **1**<br>(TAE684) |  |  | 8.1 ± 0.04 | 9.0 ± 0.02 | 8.6 |
| **2** | H |  | 7.3 ± 0.06 | 7.8 ± 0.04 | 2.5 |
| **3** | H |  | 6.5 ± 0.06 | 6.3 ± 0.06 | 0.65 |
| **4** | H |  | 5.7 ± 0.21 | 8.4 ± 0.03 | 238 |
| **5** | H |  | ND | 5.2 ± 0.08 | ND |
| **6**<br>(WEL028) |  |  | 5.0 ± 0.37 | 8.4 ± 0.03 | 232 |

Next, we performed a docking study with compound **4** in FES^S700C^ (Figure 3A). The binding mode of **4** resembled the original binding pose of TAE684 and could explain the observed structure-activity relationships. Catalytic lysine residue 590 interacts with the amide carbonyl, ideally positioning the warhead on the *ortho*-position, but not *meta*-position, to undergo a Michael addition with the engineered cysteine. The binding pose also revealed a suitable position to install an alkyne moiety on the scaffold of compound **4** to develop a two-step chemical probe for target engagement studies. This led to the synthesis of probe **6** (hereafter referred to as **WEL028**) with an *ortho*-acrylamide and alkoxyalkyne (Supplementary Scheme 5), which displayed a similar potency profile as **4** with strong inhibition of FES^S700C^ (pIC_50_ = 8.4 ± 0.03) but not FES^WT^ (pIC_50_ = 5.0 ± 0.37) (Figure 3B). The mutant-specific inhibition profile was additionally verified using the orthogonal PamChip^®^ microarray assay (Figure 3C).

**Figure 3.**
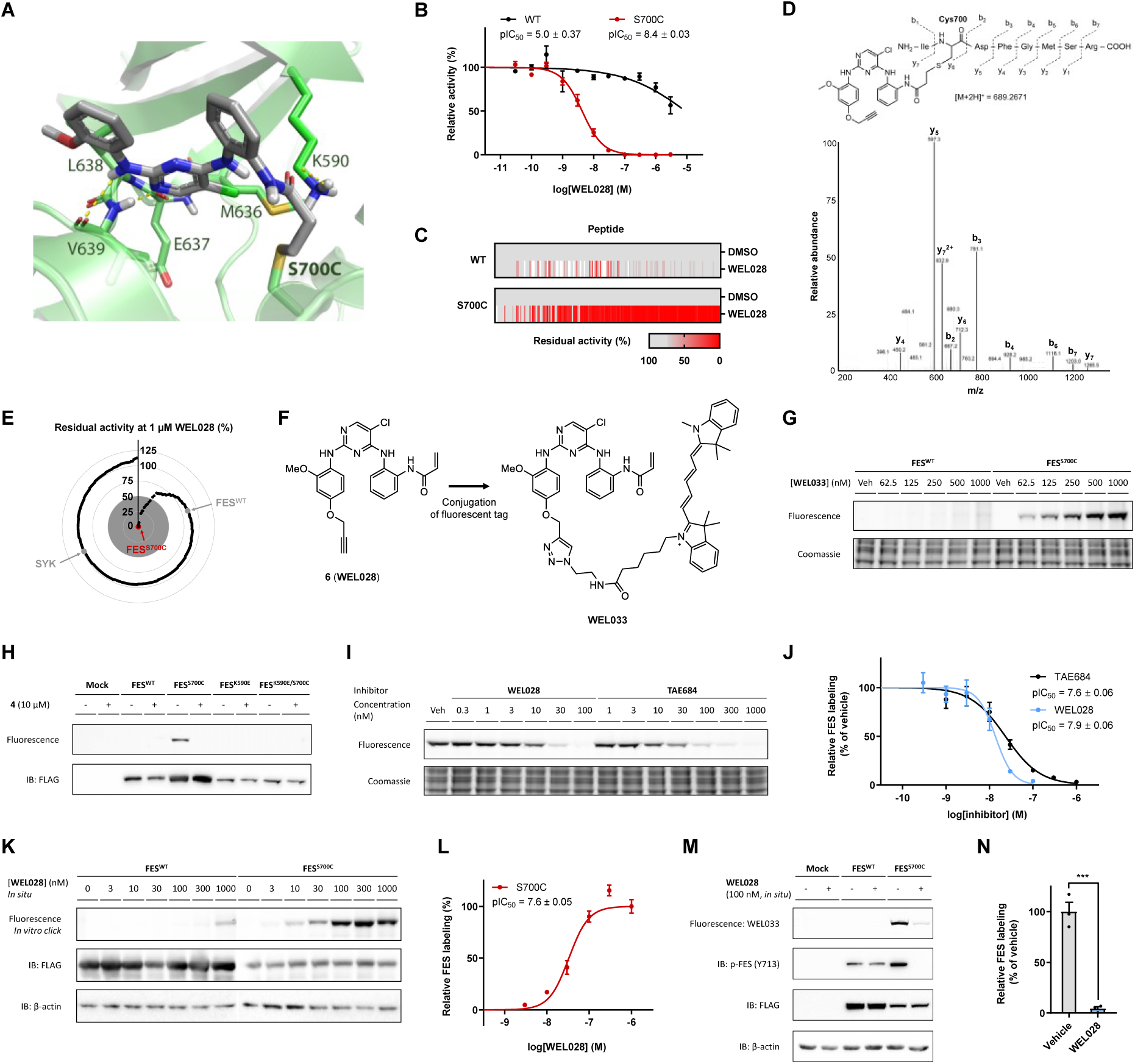
FES^S700C^ is selectively and covalently inhibited by WEL028 and can be visualized by fluorescent probe WEL033. **(A)** Proposed covalent binding mode of compound **4** to Cys700 in crystal structure of FES (PDB code: 4e93). **(B)** *In vitro* inhibition profile of FES^WT^ and FES^S700C^ by WEL028 (TR-FRET assay, n = 3). **(C)** Inhibition profile of WEL028 (100 nM) on FES^WT^ and FES^S700C^ on peptide substrates (PamChip® microarray, normalized to vehicle-treated control, n = 3). **(D)** MS/MS-based identification of WEL028 covalently bound to Cys700. Precursor ion (*m/z* [M+2H]^2+^ = 689.2671) was fragmented and signature ions are shown. Precursor ion was not observed in vehicle-treated control. **(E)** *In vitro* selectivity profile of WEL028 (1 µM, 1 h preincubation) on 279 recombinant kinases, visualized as radar plot (each data point is an individual kinase). Data represent means (n = 2). **(F)** Design of fluorescent probe WEL033 (Cy5-conjugate). **(G)** Dose-dependent labeling of full-length FES^S700C^ by WEL033 in HEK293T cell lysate. **(H)** Labeling by WEL033 is specific, exclusive for FES^S700C^ and dependent on catalytic lysine 590. Lysates were preincubated with vehicle or **4** and labeled by WEL033 (250 nM). (**I-J**) Visualization of FES^S700C^ target engagement by WEL028 and TAE684. Lysate was preincubated with WEL028 or TAE684 and labeled by WEL033 (250 nM). Band intensities were normalized to vehicle-treated control (n = 3). (**K-L**) WEL028 engages recombinantly expressed FES^S700C^ in live cells. Transfected HEK293T cells were treated with WEL028 and labeled proteins were visualized using click chemistry. Band intensities were normalized to highest concentration (n = 3). (**M-N**) WEL028 blocks FES^S700C^ but not FES^WT^ autophosphorylation (visualized by immunoblot using anti-phospho-FES Y713). Band intensities were normalized to vehicle-treated control (n = 3). Data represent means ± SEM. Statistical analysis: two-tailed *t*-test: *** *P* < 0.001.

To confirm that WEL028 undergoes Michael addition to cysteine 700, we analyzed the WEL028-FES^S700C^ complex in more detail using mass spectrometry. Recombinant FES^S700C^ was incubated with WEL028, digested to peptide fragments with trypsin and subsequently analyzed by LC-MS/MS to identify the covalent adduct (Figure 3D and S2). The mass of the expected FES peptide covalently bound to WEL028 was identified, whereas this mass was not present in a vehicle-treated control sample. The parent ion was further investigated by collision-induced dissociation and the corresponding MS/MS fragmentation pattern showed clear ladders of predicted b and y ions, which confirmed covalent addition of WEL028 to Cys700.

Cys700 is located directly adjacent to the highly conserved DFG motif. A substantial number of kinases also harbors a native cysteine at this position, which might have implications for the kinome-wide selectivity of WEL028.^41^ We therefore assessed the selectivity using the SelectScreen™ screening technology in a panel of 279 wild-type, mammalian kinases including all kinases with a native cysteine residue at any position in the active site. The assays were performed at a single dose of 1 μM with 1 h of preincubation to identify the full spectrum of potential off-target kinases (Figure S3). Only 19 of the tested 279 kinases showed >50% inhibition under these conditions, meaning that WEL028 exhibited a >100-fold selectivity window against the residual 260 kinases (93% of tested kinases, Figure 3E). Subsequently, dose-response experiments were performed for kinases showing >50% inhibition in the initial screen (Table S3 and Figure S4). The only off-targets of WEL028 with a pIC_50_ ≤ 7 were LRRK2 (pIC_50_ = 7.3 ± 0.05) and MKNK2 (pIC_50_ = 7.0 ± 0.06). Importantly, the potency of WEL028 on FES^S700C^ (pIC_50_ = 8.4 ± 0.03) was unmatched by any of the tested kinases, with a minimal 10-fold apparent selectivity window in all cases. Thus, WEL028 was identified as a complementary two-step probe for engineered FES^S700C^ that does not label wild-type FES and is highly selective over other kinases.

### Target engagement studies with complementary one-step probe WEL033

Next, a one-step fluorescent probe for FES^S700C^ was synthesized to facilitate visualization of target engagement: a Cy5-conjugated analog of WEL028 termed WEL033 (Figure 3F, Scheme 6). WEL033 dose-dependently labeled recombinantly expressed full-length FES^S700C^ but not FES^WT^ in HEK293T cell lysate (Figure 3G). Similar results were obtained using two-step labeling of WEL028-treated lysate clicked to Cy5-azide *in vitro* (Figure S5A-B). Complete labeling was achieved within 15 min and this labeling was stable up to 60 min (Figure S5C). Fluorescent labeling of FES^S700C^ was observed regardless of its autophosphorylation state, suggesting that WEL033 covalently binds to both catalytically active and inactive FES (Figure S6A). Interestingly, introduction of a secondary K590E mutation abolished labeling by WEL033 (Figure 3H). This could indicate that Lys590 is essential for covalent binding of WEL028 to the FES active site, possibly by coordination of Lys590 that positions the acrylamide warhead to undergo covalent addition to Cys700 as predicted by docking studies (Figure 3A).

To visualize target engagement of FES^S700C^, a competitive binding assay was performed with WEL033 in lysates overexpressing full-length FES^S700C^. Two inhibitors (TAE684 and WEL028) were able to prevent the labeling of FES^S700C^ in a dose-dependent manner (pIC_50_ = 7.6 ± 0.06 and pIC_50_ = 7.9 ± 0.06, respectively; Figure 3I-J). Furthermore, time-dependent displacement experiments were performed to study the mode of action of TAE684 and WEL028 (Figure S6). After inhibitor incubation at their respective IC_80_ concentrations, labeling of FES^S700C^ activity by WEL033 recovers for TAE684 but not for WEL028, indicating a reversible and irreversible mode of action for these compounds, respectively. In line with these results, inhibitor washout experiments using overnight dialysis also showed sustained inhibition by WEL028 but not TAE684 (Figure S6). Thus, gel-based probe labeling experiments using lysates of cells expressing full-length engineered FES^S700C^ is a valuable orthogonal method to standard biochemical assays using purified, truncated proteins.

Subsequently, it was investigated whether WEL028 could also engage FES^S700C^ in living cells. To this end, HEK293T cells overexpressing FES^WT^ or FES^S700C^ were incubated with various concentrations of WEL028, after which cells were harvested and lysed. WEL028-labeled proteins were then visualized using click chemistry. Dose-dependent labeling of FES^S700C^ was observed (Figure 3K-L), which indicates that WEL028 is cell-permeable and can serve as a two-step probe in living cells. Autophosphorylation of Tyr713 on the activation loop of FES is a hallmark for its kinase activity.^14,43^ Consequently, immunoblot analysis using a phospho-specific antibody for pY713 revealed that WEL028 fully abolished autophosphorylation of FES^S700C^ but not FES^WT^ (Figure 3M-N). This indicates that the target engagement as measured by gel-based probe labeling assay correlates with the functional activity of FES^S700C^ as determined in the biochemical and immunoblot assays.

### Applicability to other kinases

To investigate the broader applicability of our strategy employing engineered kinases, we explored whether the complementary probes could also target corresponding DFG-1 cysteine mutants of kinases other than FES. To this end, the DFG-1 residue (Ser701) of the FES-related kinase FER was mutated into a cysteine. FER^WT^ and FER^S701C^ were recombinantly expressed, purified and biochemically characterized and exhibited similar affinity for ATP (K_M_ = 11 μM for FERWT and K_M_ = 3.5 μM for FERS701C; Figure S7A). Profiling the panel of synthesized compounds on FER^WT^ and FER^S701C^ (Table S4 and Figure S7B-F) revealed that WEL028 potently targeted mutant but not wild-type FER (pIC_50_ = 6.3 ± 0.36 for FERWT, pIC_50_ = 8.2 ± 0.06 for FERS701C). Moreover, incubation of cell lysates from HEK293T cells overexpressing FER^S701C^ with WEL033 resulted in dose-dependent labeling that was prevented by preincubation with inhibitor **4** (Figure S8A-B). These results endorse that WEL028 is the first compound that allows acute modulation of FES activity without affecting FER.

Subsequently, we generated cysteine mutants of three other tyrosine kinases with lower sequence similarity and different amino acids at the DFG-1 position: LYN, PTK2 and PAK4 (resulting in LYN^A384C^, PTK2^G536C^ and PAK4^S457C^, respectively). Overexpression of these kinases and incubation with complementary probes exhibited dose-dependent, mutant-specific labeling (Figure S8C-H). Of note, no labeling was observed for LYN^A384C^ with one-step probe WEL033 (data not shown), which may indicate that the bulky Cy5 fluorophore prohibits active site binding in this particular case. The chemical structure of WEL028 could clearly be further optimized to improve its potency on these mutant kinases individually. Nevertheless, these results suggest that our chemical genetics strategy employing mutagenesis of the DFG-1 residue is not exclusively applicable to FES, but is possibly also suited to visualize target engagement of other kinases.

### Step 3 – CRISPR/Cas9 gene editing enables visualization of endogenous FES

To obtain a physiologically relevant model system and avoid the use of transient overexpression, CRISPR/Cas9 gene editing was employed to introduce the S700C mutation endogenously in the human HL-60 promyeloblast cell line. HL-60 cells are capable to undergo differentiation along the monocyte/macrophage as well as neutrophil lineage, depending on the differentiation agent^44^ (Figure 4A). In addition, HL-60 cells have been widely used to study neutrophil function^45^ as an experimentally tractable alternative for primary neutrophils, which have a short life-span and cannot be grown in cell culture.^46^

**Figure 4.**
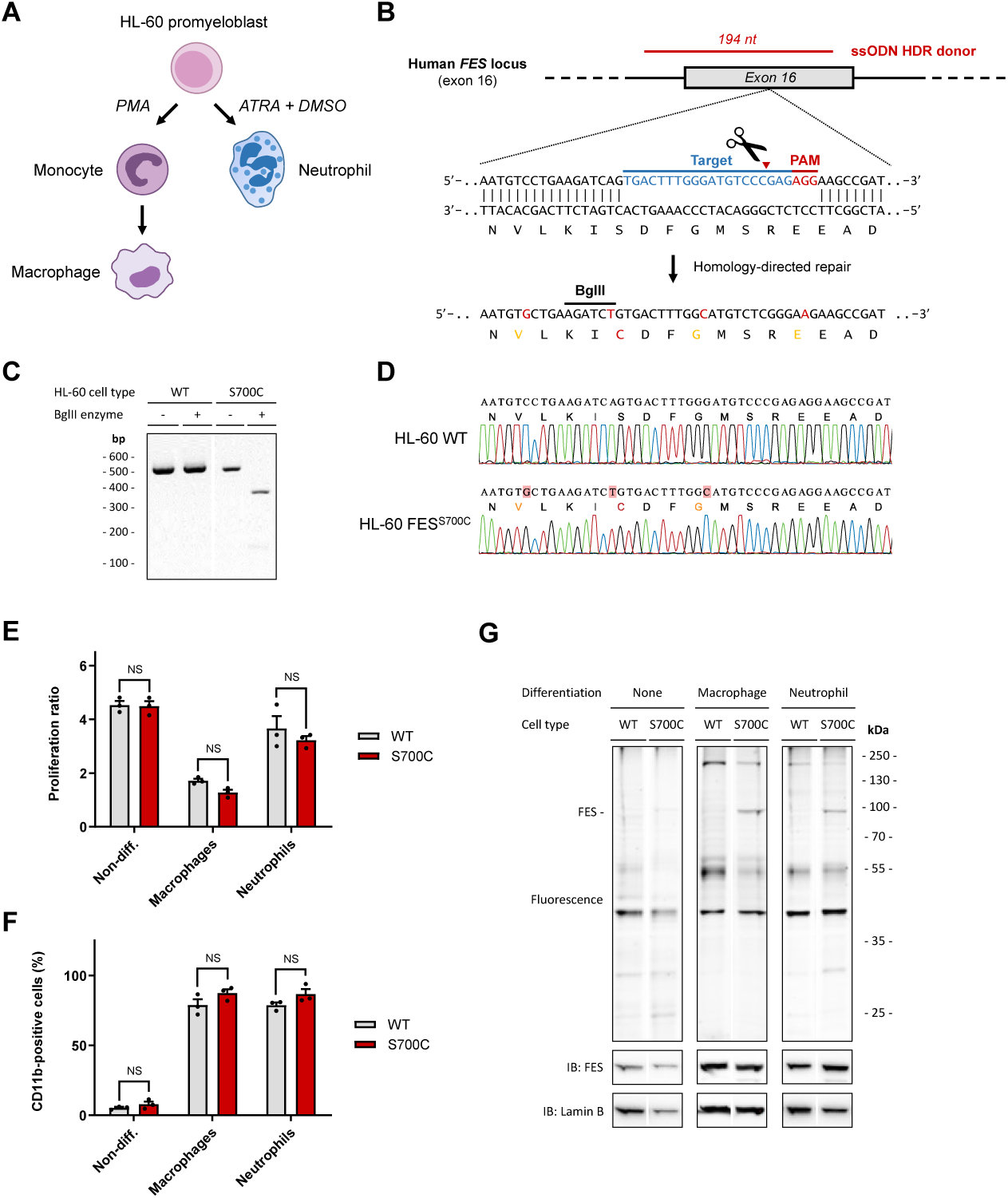
Visualization of endogenous FES^S700C^ and its target engagement in CRISPR/Cas9-edited HL-60 cells. **(A)** HL-60 cells can differentiate into macrophages upon stimulation with PMA or into neutrophils with all-*trans*-retinoic acid (ATRA) and DMSO. **(B)** CRISPR/Cas9 gene editing strategy. Selected sgRNA (bold blue) directs Cas9 to cleave at predicted site (red triangle). A ssODN homology-directed repair (HDR) donor template (red) flanks introduced mutations with 80 bp homology arms. The S700C mutation generates a BglII restriction site along with three silent mutations (orange) to remove PAM sites. Of note, two of the three mutated PAM sites correspond to sgRNAs not used in this study. PAM: protospacer-adjacent motif. **(C)** Restriction-fragment length polymorphism (RFLP) assay for identification of HL-60 FES^S700C^ clone. Genomic region was amplified by PCR and amplicons were digested with BglII. Expected fragment size after digestion: 365 + 133 bp. **(D)** Sanger sequencing traces of WT HL-60 cells and homozygous FES^S700C^ HL-60 clone. No deletions, insertions or undesired mutations were detected. **(E)** CD11b surface expression of HL-60 cells prior to and after differentiation, analyzed by flow cytometry. Threshold for CD11b-positive cells was determined using isotype control antibody. **(F)** Reduction in proliferation upon differentiation is similar for WT and FES^S700C^ cells. Proliferation ratio: cell number after differentiation divided by cell number before differentiation. **(G)** Endogenous FES is visualized by WEL033 in differentiated HL-60 FES^S700C^ cells. Wild-type or FES^S700C^ HL-60 cells promyeloblasts, macrophages or neutrophils were lysed, followed by labeling with WEL033 (1 μM). FES expression increases upon differentiation (anti-FES immunoblot).

To sensitize endogenously expressed FES in HL-60 cells to the mutant-specific probe, a single guide (sg)RNA target was selected with predicted site of cleavage in close proximity of the desired mutation in exon 16 of the *Fes* locus (Figure 4B). In conjunction, a single-stranded oligodeoxynucleotide (ssODN) homology-directed repair (HDR) donor was designed, aimed to introduce the target S700C mutation along with the implementation of a restriction enzyme recognition site to facilitate genotyping using a restriction fragment length polymorphism (RFLP) assay. Of note, the ssODN donor included also silent mutations to prevent cleavage of the ssODN itself or recleavage of the genomic locus after successful HDR (Figure 4B). HL-60 cells were nucleofected with plasmid encoding sgRNA and Cas9 nuclease along with the ssODN donor, followed by single cell dilution to obtain clonal cultures. We identified a homozygous S700C mutant clone by RFLP analysis (Figure 4C). Sanger sequencing verified that the mutations had been successfully introduced without occurrence of undesired deletions or insertions (Figure 4D). No off-target cleavage events were found in a predicted putative off-target site (Table S5 and Figure S9).

Comprehensive biochemical profiling of FES^S700C^ showed no functional differences compared to wild-type FES in any of the performed *in vitro* assays (Figure 2). In addition, we validated that HL-60 FES^S700C^ cells differentiated into neutrophils or macrophages in an identical fashion as wild-type HL-60 cells. The percentage of differentiated cells after treatment with differentiation agents was quantified by monitoring surface expression of CD11b, a receptor present on HL-60 macrophages and neutrophils but not on non-differentiated HL-60 cells (Figure 4E).^47^ No significant differences were observed between WT and FES^S700C^ HL-60 cells. In line with this observation, wild-type and mutant cells undergoing differentiation demonstrated a similar decrease in proliferation (Figure 4F). Moreover, HL-60 FES^S700C^ cells acquired typical monocyte/macrophage morphology (e.g. adherence to plastic surfaces, cell clumping and cellular elongation) comparable to wild-type cells (Figure 5G).

**Figure 5.**
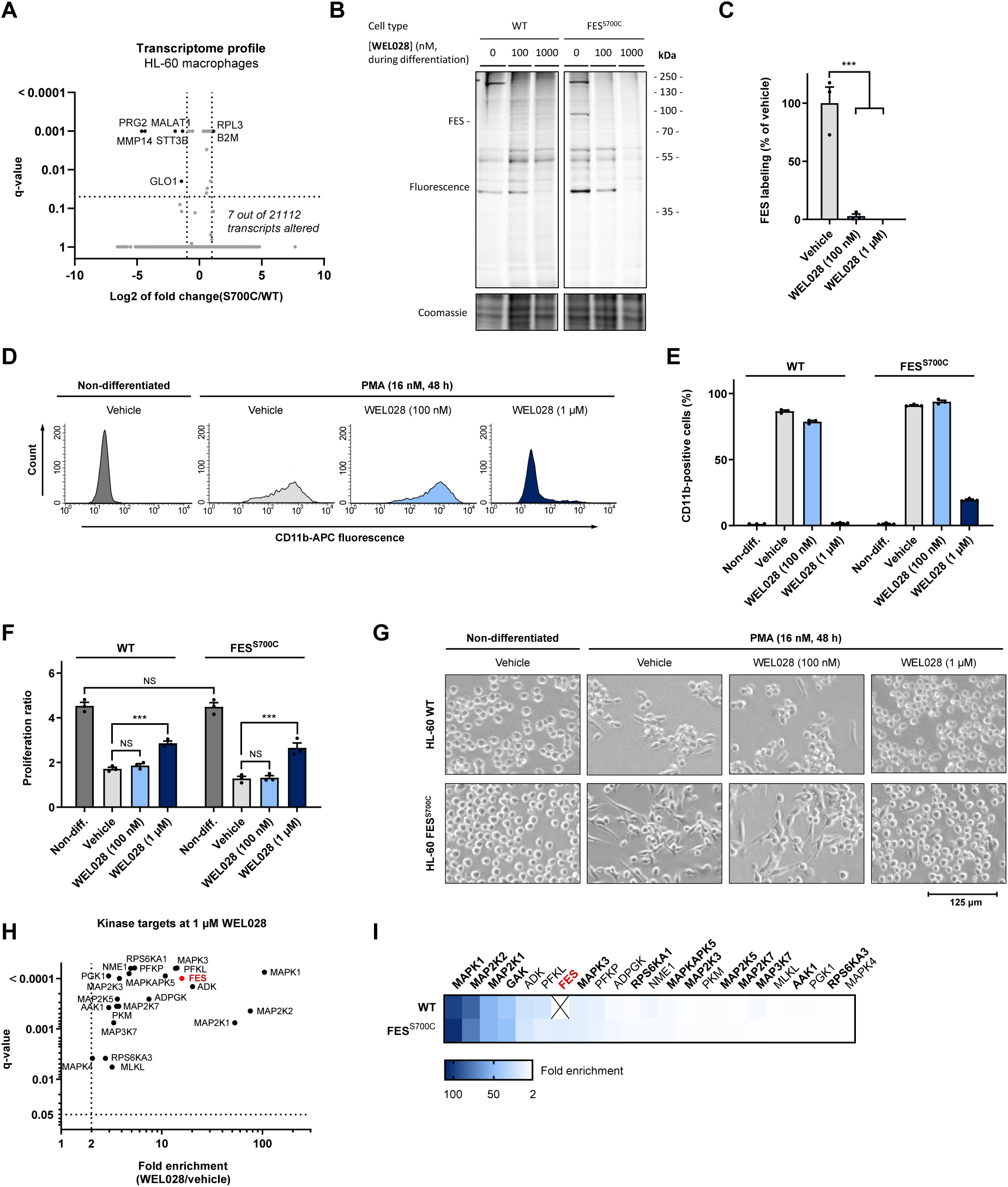
Comparative target engagement profiling in WT and FES^S700C^ HL-60 cells reveals FES activity is dispensable for macrophage differentiation. (**A**) Transcriptome profile of WT and FES^S700C^ HL-60 macrophages analyzed by TempO-Seq. Differentially expressed genes were identified using following cut-offs: log2 of fold change FES^S700C^ over WT HL-60 cells < −1 or > 1 with q-value < 0.05. (**B-C**) Gel-based target engagement profile of WEL028 on WT and FES^S700C^ HL-60 cells treated during PMA-induced differentiation. Cells were pretreated with vehicle or WEL028 (100 or 1000 nM, 1 h) prior to induction of differentiation towards monocytes/macrophages with PMA (16 nM, 48 h). Medium was refreshed with growth medium containing WEL028 and PMA after 24 h to maintain full FES inhibition. Lysates were post-labeled with WEL033 (1 µM, 30 min, rt). Band intensities were normalized to vehicle-treated control. (**D-E**) CD11b surface expression analyzed by flow cytometry. Threshold for CD11b-positive cells was determined using isotype control antibody. Histograms illustrate representative replicates of FES^S700C^ cells. **(F)** Proliferation of WT and FES^S700C^ HL-60 cells subjected to macrophage differentiation. Proliferation ratio: live cell number after differentiation divided by live cell number before differentiation. **(G)** Morphological inspection of WT and FES^S700C^ HL-60 cells. Shown images are representative for multiple acquired images at 20x magnification from replicates. **(H)** Chemical proteomics-based identification of WEL028 kinase targets at 1 μM in FES^S700C^ HL-60 cells undergoing differentiation towards macrophages. Kinases with >2-fold enrichment compared to vehicle control (q < 0.05) were designated as targets. Values represent means of fold enrichment (n = 3). **(I)** Heat map representation of WEL028 (1 µM) target engagement profile in WT and FES^S700C^ HL-60 cells undergoing macrophage differentiation, determined by chemical proteomics. Kinases with native cysteine at DFG-1 position are shown in bold. All data represent means ± SEM. Statistical analysis: two-tailed t-test with Holm-Sidak multiple comparisons correction: *** P < 0.001, NS if P > 0.05.

Next, wild-type and HL-60 FES^S700C^ cells were differentiated into macrophages or neutrophils and corresponding cell lysates were incubated with fluorescent probe WEL033 to visualize endogenous FES (Figure 4G). In-gel fluorescence scanning of the WEL033-labeled proteome of FES^S700C^ HL-60 neutrophils and macrophages revealed a band at the expected MW of FES (∼93 kDa), which was absent in wild-type HL-60 cells. Furthermore, the fluorescent band was less prominent in non-differentiated HL-60 FES^S700C^ cells, likely due to lower FES expression levels prior to differentiation (Figure 4G, anti-FES immunoblot). Of note, WEL033 labeled a number of additional proteins (MW of ∼200, ∼55 and ∼40 kDa, respectively) at the concentration used for FES detection (1 µM). In short, these results demonstrate that endogenously expressed engineered FES can be visualized using complementary chemical probes.

### Step 4 – Visualization of cellular target engagement in differentiating HL-60 cells

FES was previously reported as an essential component of the cellular signaling pathways involved in myeloid differentiation.^48,49^ However, most of these studies relied on the use of overexpression, constitutively active mutants, or antisense-based knockdown of FES. In addition, it remains unclear whether this role of FES is dependent on its kinase activity.

To first confirm that mutant HL-60 macrophages exhibited minimal transcriptional alterations compared to parental WT macrophages (*e.g.* due to clonal expansion), we performed a targeted transcriptomics analysis using the TempO-Seq technology (Figure 5A).^50^ Only seven out of 21112 of the identified transcripts (0.03%) were significantly altered in FES^S700C^ compared to WT HL-60 macrophages, which indicates that introduction of this mutation minimally disturbs gene expression.

Next, living wild-type and FES^S700C^ HL-60 cells were incubated with WEL028 (100 nM or 1 µM) during PMA-induced differentiation towards macrophages. To ensure complete FES inhibition at the moment of differentiation initiation, cells were pretreated with WEL028 2 h prior to addition of PMA. The growth medium was refreshed after 24 h to maintain FES inhibition, since recovery of active FES upon prolonged (> 48 h) exposure to WEL028 was observed, possibly due to protein resynthesis. Cells were harvested and lysed, followed by labeling of residual active FES^S700C^ by WEL033 (Figure 5B). This revealed full target engagement of engineered FES at a concentration of 100 nM WEL028 (Figure 5C), with only two prominent off-targets (∼150 kDa and ∼40 kDa). At a higher concentration of 1 µM, WEL028 was substantially less selective (Figure 5B) and inhibited the labeling of multiple proteins.

The percentage of differentiated cells after treatment with the differentiation agent was quantified by monitoring surface expression of CD11b, a receptor present on HL-60 macrophages but not on non-differentiated HL-60 cells.^47^ Strikingly, despite complete inhibition of FES^S700C^ at 100 nM WEL028 (Figure 5C), the percentage of CD11b-positive cells was unaltered (Figure 5D-E). Cell proliferation, an indirect hallmark of differentiation, was decreased to identical levels for FES^S700C^ HL-60 cells treated with vehicle or 100 nM WEL028 (Figure 5F). Accordingly, FES^S700C^ HL-60 cells treated with 100 nM WEL028 acquired macrophage morphology (*e.g.* adherence to plastic surfaces, cell clumping and cellular elongation) comparable to vehicle-treated controls (Figure 5G). Together, these results show that complete FES inhibition does not affect PMA-induced differentiation of HL-60 cells into macrophages, suggesting that FES activity is dispensable for this process.

Remarkably, FES^S700C^ cells undergoing PMA-induced differentiation in presence of a higher concentration of WEL028 (1 µM) completely failed to express CD11b, exhibited a less pronounced decrease in proliferation and displayed phenotypic characteristics similar to non-differentiated cells (Figure 5D-G). Competitive probe labeling experiments revealed that WEL028 targets multiple off-targets at a concentration of 1 µM (Figure 5B), which suggested that the observed block in differentiation might be due to off-targets. A beneficial feature of the used chemical genetic strategy is that wild-type cells can be included to account for these off-targets. Indeed, 1 µM WEL028 had similar effects on CD11b surface expression, proliferation and morphology of wild-type HL-60 cells subjected to differentiation (Figure 5D-G). This verifies that the functional effects of WEL028 at 1 µM can indeed be attributed to off-target rather than on-target effects.

Quantitative label-free chemical proteomics was used to identify the off-targets of WEL028 at this high concentration. The WEL028-labeled proteome was conjugated to biotin-azide post-lysis, followed by streptavidin enrichment, on-bead protein digestion and analysis of the corresponding peptides using mass spectrometry. Significantly enriched kinases in WEL028-treated samples compared to vehicle-treated samples were designated as targets (Figure 5H-I). Identified off-targets included protein kinases harboring native cysteines at the DFG-1 position (depicted in bold), as well as several metabolic kinases (ADK, PFKL, PFKP, ADPGK, NME1, PKM, PGK1), although the latter lack a DFG-motif. Notably, the off-target profile of HL-60 WT and FES^S700C^ cells was identical with the exception of FES, which was exclusively present in mutant cells. This highlights that, despite a limited number of off-targets, WEL028 can be effectively used in comparative target validation studies between WT and FES^S700C^ HL-60 cells.

### Step 5 - FES activates SYK by phosphorylation of its linker residue Y352 and contributes to phagocytic uptake of E. coli by HL-60 neutrophils

Neutrophils belong to the first line of defense against invading bacteria. They are recruited to the site of infection and their primary function is to phagocytize and kill the pathogens. Neutrophil phagocytosis is a complex process that can occur via (a cross-talk of) various receptors, including pathogen-associated molecular pattern (PAMP) receptors, Fcγ receptors (FcγRs) and complement receptors (CRs).^51^ Since FES was previously reported to regulate cell surface receptors, including TLR4 in macrophages^52^ and FcεRI in mast cells^11,19^, we wondered whether FES might be involved in neutrophil phagocytosis. First, we confirmed that treatment of live neutrophils with a low concentration of WEL028 (100 nM, 1 h) resulted in complete and selective inhibition of FES in the mutant cells (Figure 6A-B). Partial inhibition of two off-targets (∼80% inhibition of ∼150 kDa protein and ∼10% inhibition of ∼40 kDa protein) in both WT and mutant cells was observed (Figure 6A-B). To identify these off-targets, cells were harvested and the WEL028-bound targets were identified using chemical proteomics. This identified GAK and MAPK1 as the only significantly enriched proteins in both WT and FES^S700C^ neutrophils (Figure 6C), thereby confirming the selectivity of the probe at this low concentration. Of note, the size of these proteins (143 kDa for GAK and 41 kDa for MAPK1) corresponds to the size of the two fluorescent bands visualized on gel. FES was exclusively identified as target of WEL028 in the mutant cells.

**Figure 6.**
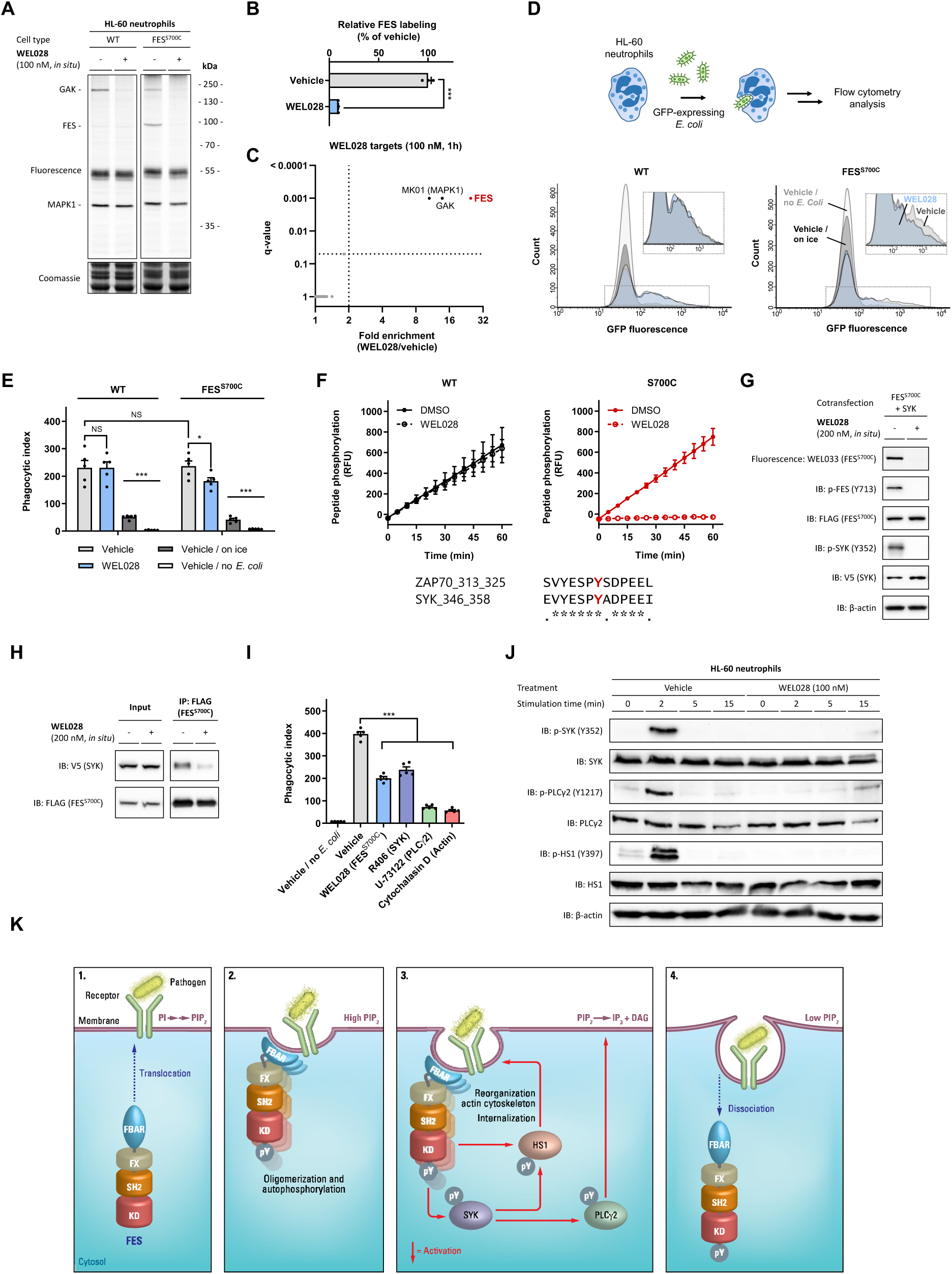
FES mediates phagocytic uptake of *E. coli* in HL-60 neutrophils by activating the tyrosine kinase SYK. (**A-B**) Complete FES^S700C^ inhibition at 100 nM WEL028 in HL-60 neutrophils *in situ*. Neutrophils were treated with WEL028 (100 nM, 1 h) and lysates were post-labeled (1 µM WEL033). Band intensities were normalized to vehicle-treated control (n = 3). (**C**) Chemical proteomics-based identification of WEL028 kinase targets at 100 nM in FES^S700C^ HL-60 neutrophils. Kinases with >2-fold enrichment compared to vehicle control (q < 0.05) were designated as targets. Values represent means of fold enrichment (n = 4). (**D-E**) WEL028 reduces phagocytic uptake in FES^S700C^ but not WT neutrophils. HL-60 neutrophils were incubated with vehicle or WEL028 (100 nM, 1 h), after which GFP-expressing *E. coli* B834 were added (MOI = 30, 1 h, 37°C). Cells were analyzed by flow cytometry (n = 5). Phagocytic index was calculated as fraction of GFP-positive cells (number of phagocytic cells) multiplied by GFP MFI (number of phagocytized bacteria). **(F)** SYK Y352 as proposed phosphorylation site of FES, based on phosphorylation of homologous ZAP70 peptide. WEL028 abolishes phosphorylation by FES^S700C^ (PamChip® microarray, n = 3). **(G)** FES phosphorylates SYK Y352 in co-transfected U2OS cells. U2OS cells co-expressing FLAG-tagged FES^S700C^ and V5-tagged SYK were incubated with vehicle or WEL028 (200 nM, 1 h) and lysed. Lysates were labeled (250 nM WEL033) and analyzed by in-gel fluorescence and immunoblot (n = 3). **(H)** SYK interacts with FES and this interaction is dependent on FES activity. Co-transfected U2OS cells were incubated as in **G**, followed by immunoprecipitation using anti-FLAG antibody and immunoblot analysis (n = 3). **(I)** Phagocytic uptake of *E. coli* by HL-60 neutrophils is dependent on FES, SYK and PLCγ2 activity and requires actin polymerization. Incubations as in **D-E** (WEL028: 100 nM, R406: 1 µM, U-73122: 5 µM, cytochalasin D: 10 µM; n = 5). **(J)** FES rapidly phosphorylates endogenous SYK Y352 and downstream substrates HS1 Y397 and PLCγ2 Y1217 in HL-60 neutrophils infected with *E. coli*. Inhibitor incubations as in **D-E**, followed by addition of GFP-expressing *E. coli* B834 (MOI = 30, 0-2-5-15 min, 37°C) and immunoblot analysis (n = 3). **(K)** Proposed simplified model for the role of FES in neutrophil phagocytosis. Phosphorylation sites Y378 and Y397 on HS1 are shown as single site for visualization purposes. For further details, see text. PI: phosphatidylinositol, PIP_2_: phosphatidylinositol 4,5-bisphosphate, IP_3_: inositol 1,4,5-trisphosphate, DAG: diacylglycerol, FX: F-BAR extension, SH2: Src Homology 2, KD: kinase domain, pY: phosphotyrosine. Data represent means ± SEM. Statistical analysis: ANOVA with Holm-Sidak’s multiple comparisons correction: *** *P* < 0.001; * *P* < 0.05; NS if *P* > 0.05.

To measure the phagocytic uptake by HL-60 neutrophils, a flow cytometry-based assay with live GFP-expressing *E. coli* was employed (Figure 6D). Both WT and FES^S700C^ neutrophils effectively internalized bacteria, with identical phagocytic indices (Figure 6E). Control cells incubated on ice were included to account for surface binding without internalization. Interestingly, WEL028 at an adequate concentration (100 nM) for complete and selective FES inactivation (Figure 6A), reduced the phagocytic index by 30-50% in FES^S700C^ expressing cells, but not in WT HL-60 neutrophils (Figure 6D, 6E and 6I; Figure S12). Since the off-targets GAK and MAPK1 are shared among WT and FES^S700C^ HL-60 neutrophils (Figure 6A and S10), these results indicate that on-target FES inhibition is responsible for the observed reduction in phagocytosis of *E. coli*.

To gain insight in the molecular mechanisms of FES-mediated phagocytosis, we examined the previously obtained substrate profile in more detail (Figure 2E, Table S1). A peptide of the non-receptor tyrosine kinase ZAP70 was identified as prominent FES substrate with high signal intensity. Incubation with WEL028 abolished peptide phosphorylation by FES^S700C^, but not FES^WT^ (Figure 6F). Although ZAP70 is predominantly linked to immune signaling in T-cells, its close homologue SYK is ubiquitously expressed in various immune cells, including neutrophils.^46^ Moreover, SYK is part of signaling pathways linked to pathogen recognition and involved in bacterial uptake by neutrophils.^53^ The identified ZAP70 peptide substrate shows high sequence similarity to its SYK counterpart surrounding Y352 (Figure 6F). To validate that SYK is a downstream target of FES, SYK-V5 and FES^S700C^-FLAG were co-transfected in U2OS cells. First, it was confirmed that overexpression of FES^S700C^ led to autophosphorylation of FES at Y713, which was sensitive to WEL028 (Figure 6G). Subsequent immunoblot analysis using a SYK Y352 phospho-specific antibody showed that SYK was phosphorylated in a WEL028-dependent manner (Figure 6G). Of note, WEL028 did not inhibit SYK in the kinome screen (Figure 3E and S3) and did not affect SYK pY352 levels when FES^S700C^ was omitted (Figure S11). In addition, immunoprecipitation against FES^S700C^ using an anti-FLAG antibody revealed a physical interaction between FES and SYK as witnessed by immunoblot against the V5-tag of SYK (Figure 6H). Importantly, this interaction was dependent on the activation status of FES, because WEL028 inhibited the co-precipitation of SYK with FES. Taken together, these results suggest that SYK Y352 is a direct substrate of FES.

To verify that endogenous SYK is also involved in phagocytic uptake of *E. coli* by neutrophils, HL-60 neutrophils were incubated with the potent SYK inhibitor R406. SYK inhibition reduced phagocytosis to similar levels as observed for WEL028 (Figure 6I). Of note, U-73122 and Cytochalasin D were taken along as positive controls to verify that the phagocytosis is mediated via phospholipase C gamma 2 (PLCγ2) and actin polymerization, respectively.^51^ Finally, we verified whether the downstream signaling pathway of SYK was modulated in a FES-dependent manner by using immunoblot analysis with phospho-specific antibodies for SYK Y352, hematopoietic cell-specific protein-1 (HS1) Y397 (an actin-binding protein) and PLCγ2 Y1217 (Figure 6J). Indeed, a transient phosphorylation of SYK, HS1 and PLCγ2 was observed, which peaked at 2 minutes after incubation with bacteria (Figure 6J). Inhibition of FES^S700C^ completely blocked the phosphorylation of these proteins, thereby demonstrating FES activation is an early event in the phagocytosis of *E. coli* in HL-60 neutrophils.

## Discussion

In this report, we present a novel chemical genetics strategy to acutely modulate kinase activity and visualize kinase target engagement, combining precise gene editing with the design and application of complementary covalent inhibitors. The field of chemical genetics has previously generated tools to aid in kinase target validation^54,55^, such as the powerful “analog-sensitive” *(AS)* technology, where the gatekeeper residue is changed into a less bulky residue, enabling the kinase of interest to accommodate bulky ATP analogs in its active site.^56^ However, these analogs do not form covalent adducts with the kinase and therefore do not readily allow visualization of target engagement. Furthermore, mutagenesis of gatekeeper residues may result in impaired catalytic activity and the suboptimal pharmacokinetic properties of ATP analogs used in the *AS* technology limit their applicability for *in vivo* target validation studies.^57^ The concept of “covalent complementarity” is based on mutagenesis of the gatekeeper^36^ or gatekeeper+6^38^ residue into a cysteine to function as nucleophile. Although this allows the development of covalent probes and thereby target engagement studies, a second mutation in the active site was required to improve gatekeeper cysteine reactivity or compound selectivity and potency.^36,37^ This is particularly challenging when moving to an endogenous model system, since it would involve two independent CRISPR/Cas9 gene editing events to introduce these two point mutations.

Here, we identified the DFG-1 residue as an excellent position for introduction of a nucleophilic cysteine to react with an acrylamide as a complementary warhead, with no need for second point mutations to improve cysteine reactivity or inhibitor selectivity. Furthermore, mutagenesis of the DFG-1 position into a cysteine is functionally silent: it does not affect FES catalytic activity nor its substrate recognition and SH2 domain binding profile. FES^S700C^ showed a minor increase in ATP binding affinity, but this difference in K_M_ is unlikely to have any consequences at physiologically relevant ATP concentrations, which are typically in the millimolar range.^41^ Although nearly 10% of all known kinases have a native DFG-1 cysteine residue, many kinases harboring a DFG-1 cysteine showed limited or no inhibition by WEL028 (Figure S3 and Figure 6C), suggesting that the choice of the chemical scaffold constitutes an additional selectivity filter. An acrylamide group was selected as the electrophile to react with the intended cysteine, since it exhibits sufficient reactivity towards cysteines only when appropriately positioned for a Michael addition reaction and has limited reactivity to other intracellular nucleophiles.^58^ This may additionally prevent non-specific interactions with targets outside of the kinase cysteinome. Given the favorable selectivity profile and cellular permeability of WEL028, we postulate that the diaminopyrimidine scaffold is a useful addition to the toolbox of covalent complementary probes applied in chemical genetic strategies, which previously consisted of mainly quinazolines and pyrazolopyrimidines.^36,38^

Previously reported chemical genetic methods are relying on overexpression systems that disturb signal transduction cascades.^40^ In contrast, a major benefit of our chemical genetics strategy lies in the unprecedented control that our method allows to exert over a biological system without disturbing the cellular homeostasis. We replaced one single atom out of 13.171 atoms in a protein by changing only one base-pair out of over 3.000.000.000 base-pairs in the human genome. Yet, this allowed us to rationally design and synthesize a chemical probe that visualizes and inhibits the engineered kinase activity in human cells, thereby enabling target engagement and validation studies. Arguably, this minimal change at the genome and protein level ensures that its regulation at transcriptional and (post)-translational level activity are minimally disturbed. In fact, we could demonstrate that the mutant protein activity, substrate preference and protein-protein interactions were similar to the WT protein. Furthermore, WT and mutant cells behaved similarly in various functional assays (e.g. proliferation, differentiation and phagocytosis). Targeted sequencing experiments further confirmed minimal transcriptional differences (0.03% of read transcripts) between the clonal FES^S700C^ cell line and parental WT cell line. Although the low efficiency of HDR-mediated mutagenesis did not allow us to identify more than one homozygous mutant clone, recent developments in CRISPR/Cas9 gene editing, base editing^59^ and prime editing^60^ will undoubtedly improve this efficiency in the future.

Our chemical genetics strategy allows temporal control in modulating kinase activity, opposed to conventional genetic approaches. In this report, we leverage this advantage by inactivating FES activity both during myeloid differentiation and in terminally differentiated neutrophils. In contrast to previous studies relying on overexpression of (constitutively active) FES or genetic knockout, we found that FES activity is dispensable for differentiation towards macrophages in HL-60 cells. Caution should thus be taken when using overexpression systems, especially in case of artificial mutant kinases with constitutive activity, as these may induce artefacts in cellular physiology. It remains to be investigated whether FES plays a role in myeloid differentiation in other cell types, or independently of its kinase activity, *e.g.* as a interaction partner for other proteins.

We illustrated that comparative target engagement profiling in mutant and wild-type cells is a powerful approach to distinguish on-target from off-target effects. Our results highlight the relevance of visualizing target engagement to select a dose that is sufficient to completely inactivate the kinase of interest, and avoid doses that induce off-target effects. For example, differentiation of HL-60 cells was prevented at a higher concentration of WEL028 than required for complete FES inhibition. Chemical proteomics identified various members of the MAP kinase family as prominent off-targets under these conditions. Interestingly, MAPK1/3 and MAP2K1/2 are part of signaling pathways activated upon PMA treatment and are reported to be essential for HL-60 cell differentiation along the monocyte/macrophage lineage.^47^ Although MAPK1 and MAPK3 were no prominent targets of WEL028 in the single-dose kinome screen at 1 μM (< 50% inhibition) (Figure 3E and S3), both were identified as off-targets at this concentration *in situ*. This could result from different *in vitro* and *in situ* inhibitory potencies^61^ or high total abundance of these kinases in cells. MAP2K1 and MAP2K2 were already identified in the *in vitro* selectivity assay and although WEL028 showed only a moderate potency on these targets (pIC_50_ = 6.6 ± 0.06), both are likely to be inhibited at a concentration of 1 µM.

Furthermore, our chemical genetics strategy enabled us to uncover a novel biological role for FES in neutrophils. FES was demonstrated to play a key role in the phagocytic uptake of bacteria in neutrophils by activating SYK and downstream substrates HS1 and PLCγ2. Our results indicate that FES is involved in internalization of the immunoreceptor-bacterium complex, which is in line with previous studies that reported a role for FES in regulating surface expression of TLR4 in macrophages^52^ and adhesion molecule PSGL-1 in leukocytes.^22^ In combination with previous data reported in the literature (reviewed in ref. 19), we can propose the following model (Figure 6K). One of the first events in response to bacterial recognition by neutrophil surface receptors is the formation of the lipid phosphatidylinositol 4,5-bisphosphate (PIP^2^) in the membrane (Figure 6K, panel 1).^62^ FES normally resides in the cytosol in an inactive conformation, but translocation to the PIP_2_−rich membrane may occur by binding via its F-BAR domain. This triggers the formation of oligomers and auto-activation by phosphorylation on FES Y713 and induces membrane curvature required for particle internalization.^19^ (Figure 6K, panel 2). FES subsequently activates SYK by phosphorylation of Y352, which poses an alternative activation mechanism of SYK compared to the traditional activation via binding to ITAM domains of immunoreceptors.^63^ Y352 is located in a linker region of SYK, which separates its two N-terminal SH2 domains from its C-terminal catalytic domain. Phosphorylation of this linker residue was previously shown to perturb auto-inhibitory interactions, resulting in SYK activation.^63^ SYK is known to phosphorylate HS1, an actin-binding protein involved in reorganization of the actin cytoskeleton. It can be phosphorylated on multiple tyrosine residues that all contribute to its actin remodeling function.^64^ Of note, its Y378 and Y397 residues are phosphorylated by FES in mast cells, but both sites have also been identified as substrate sites for other kinases, including SYK.^19^ Phosphorylation of HS1 by FES and/or SYK drives reorganization of the actin cytoskeleton required for internalization of the bacterium-receptor complex (Figure 6K, panel 3). Concomitantly, the phosphorylated Y352 residue in SYK could serve as binding site for the SH2 domain of PLCγ2, followed by SYK-mediated PLCγ2 activation. In turn, this would allow for degradation of PIP_2_ into diacylglycerol (DAG) and inositoltriphosphate (IP_3_), altering the membrane composition and returning it to the non-activated state: FES dissociates from the membrane and the signaling process is terminated (Figure 6K, panel 4). This model thus proposes a feedback mechanism in which FES indirectly regulates its own localization and activation by modulating PLCγ2 activity via SYK. It is also relevant to note that FES possesses an FX domain that binds phosphatidic acid (PA) and activates its kinase activity.^65^ PA is synthesized by phospholipase D (PLD) in response to activation of various immune receptors. PLD activation also leads to elevated intracellular Ca^2+^ levels, but the exact mechanistic pathways underlying this event remain poorly understood.^66^ FES could possibly be a molecular link that activates PLCγ2 to increase Ca^2+^ in response to PA production by PLD. Further studies are necessary to investigate the activation mechanism of FES by PA and the role of FES in the complex cross-talk between PLD and PLCγ2.

The identification of FES as a potential activator of SYK also provides new insights in previous studies reporting on FES and/or SYK function. For example, FES is involved in the development and function of osteoclasts, multinucleated cells responsible for bone resorption.^29^ Accordingly, SYK-deficient osteoclasts exhibit major defects in the actin cytoskeleton, resulting in reduced bone resorption.^67^ It remains to be determined whether FES inhibition disrupts the osteoclast cytoskeleton in a similar manner, which would make FES a potential target to treat osteoporosis and cancer-associated bone disease. Moreover, FES inhibition was shown to suppress growth of acute myeloid leukemia (AML) cells that express the FLT3-ITD mutation, but not cells expressing wild-type FLT3.^7^ Similarly, FLT3-ITD AML is more vulnerable to SYK suppression than FLT3-WT AML.^68^ It would be interesting to investigate whether FES activates SYK in these AML cells and whether altered internalization of FLT3-ITD compared to FLT3-WT perhaps explains the increased susceptibility to FES and SYK inhibition.

Two SYK inhibitors have recently been approved by the FDA and more are currently in clinical trials for the treatment of various malignancies and inflammatory diseases.^6^ The question thus rises in what other cell types and cellular processes SYK activation is dependent on FES. Notably, FES is expressed in many cell types that contribute to the pathogenesis of inflammatory diseases, such as macrophages, mast cells, neutrophils and B-cells, but future studies will prove whether FES inhibitors may be of therapeutic value in these disorders. We believe that our chemical genetics strategy may provide valuable tools to investigate FES-associated physiological processes and aid in its validation as drug target.

On a final note, our chemical genetics strategy is not limited to FES, but it is envisioned that it can be applied to other kinases, either structurally related or distinct. The selectivity acquired by combining gene editing and a complementary probe brings the advantages of acute, pharmacological inhibition without the need for extensive hit optimization programs to identify compounds of adequate potency and selectivity. Although we provided a proof-of-concept using a cell line as model system, tremendous advancements in gene editing technologies also provide means to generate mutant (stem) cells or animal models. In conclusion, we envision that the presented methodology could provide powerful pharmacological tools to study the function of poorly characterized kinases and aid in their validation as therapeutic targets.

## Acknowledgements

The Cloning & Purification Facility and Hans van den Elst are acknowledged for technical support. We thank Richard J.B.H.N. van den Berg for critically reading the Material & Methods and Jessica Schlachter for preparing the PAK4 plasmids. Prof. dr. John Kuriyan is acknowledged for providing the YopH plasmid. The department of Toxicology (Leiden Academic Centre for Drug Research) is kindly acknowledged for access to the TempO-Seq consortium. We acknowledge ChemAxon for kindly providing the Instant JChem software to manage our compound library. T.v.d.W., A.K, T.B. and M.v.d.S. would like to acknowledge NWO (VICI) & Topsector Chemistry (TKI-project “OncoDrugs”) for financial support. E.B.L. would like to thank the Agentschap Innoveren en Ondernemen (AIO) for financial support (AIO project 155028). G.J.P.v.W. thanks the Dutch Research Council (NWO-TTW Veni 14410) for funding.

## Author contributions

T.v.d.W., A.K., T.B., and M.v.d.S. designed research approach. T.v.d.W., H.d.D., R.H., T.v.d.H., N.M.P., B.I.F. and E.B.L. performed experiments and analyzed data. T.v.d.W. and M.v.d.S. wrote the manuscript. R.R., G.J.P.v.W. H.S.O., A.K. and T.B. provided useful comments and feedback to improve the manuscript.

## Data availability

Data are available from authors upon request. Raw TempO-Seq data associated with Figure 5A and raw proteomics data associated with Figure 5H-I and 6C are included as Supplementary Data files. Uncropped gel and immunoblot images are included in Figure S13.

