## Supplementary Information for "Chemical genetics strategy to profile kinase target engagement reveals role of FES in neutrophil phagocytosis via SYK activation"

**Figure S1. Determination of ATP $K_M$ for other active FES mutants and concentration-response curves of inhibitors against $FES^{WT}$ and $FES^{S700C}$.**
**Figure S2. LC-MS elution profiles of $FES^{S700C}$ incubated with WEL028 (A) or vehicle (B).**
**Figure S3. Single-point selectivity screen of WEL028 on a panel of 279 kinases.**
**Figure S4. Concentration-response curves of WEL028 against kinases with >50 % inhibition at 1 µM in initial single-dose screen.**
**Figure S5. Two-step labeling of $FES^{S700C}$ by WEL028 conjugated to Cy5-azide in vitro using click chemistry.**
**Figure S6. Characterization of irreversible, covalent binding mode of WEL028 by probe displacement and inhibitor washout experiments.**
**Figure S7. Determination of ATP $K_M$ for $FER^{WT}$ and $FER^{S701C}$ and concentration-response curves of inhibitors against $FER^{WT}$ and $FER^{S701C}$.**
**Figure S8. Applicability of chemical genetic strategy on various kinases harboring DFG-1 residues mutated into cysteines.**
**Figure S9. Analysis of putative sgRNA off-target site.**
**Figure S10. Chemical proteomics-based identification of WEL028 kinase targets at 100 nM in WT HL-60 neutrophils.**
**Figure S11. WEL028 does not affect SYK Y352 phosphorylation in U2OS cells in absence of $FES^{S700C}$.**
**Figure S12. Phagocytic uptake of E. coli by HL-60 $FES^{S700C}$ neutrophils at variable multiplicity of infection (MOI) and infection time.**
**Figure S13. Uncropped gel and blot images.**

**Table S1: Substrate identification using PamChip® activity assay.**
**Table S2: SH2 binding partner identification using PamChip® binding assay.**
**Table S3: Inhibitory potency of WEL028 against kinases with >50 % inhibition at 1 µM in initial single-dose screen.**
**Table S4: Inhibitory potency of synthesized TAE684 derivatives against FERWT and FERS701C.**
**Table S5: List of putative off-target cleavage sites for sgRNA employed in CRISPR/Cas9-mediated mutagenesis of FES.**
**Table S6: Complete list of oligonucleotide sequences.**

**Supplementary Materials & Methods – Biology**

**Supplementary Materials & Methods – Chemistry**

**Uploaded as separate file: Supplementary Data (TempO-Seq and Proteomics)**

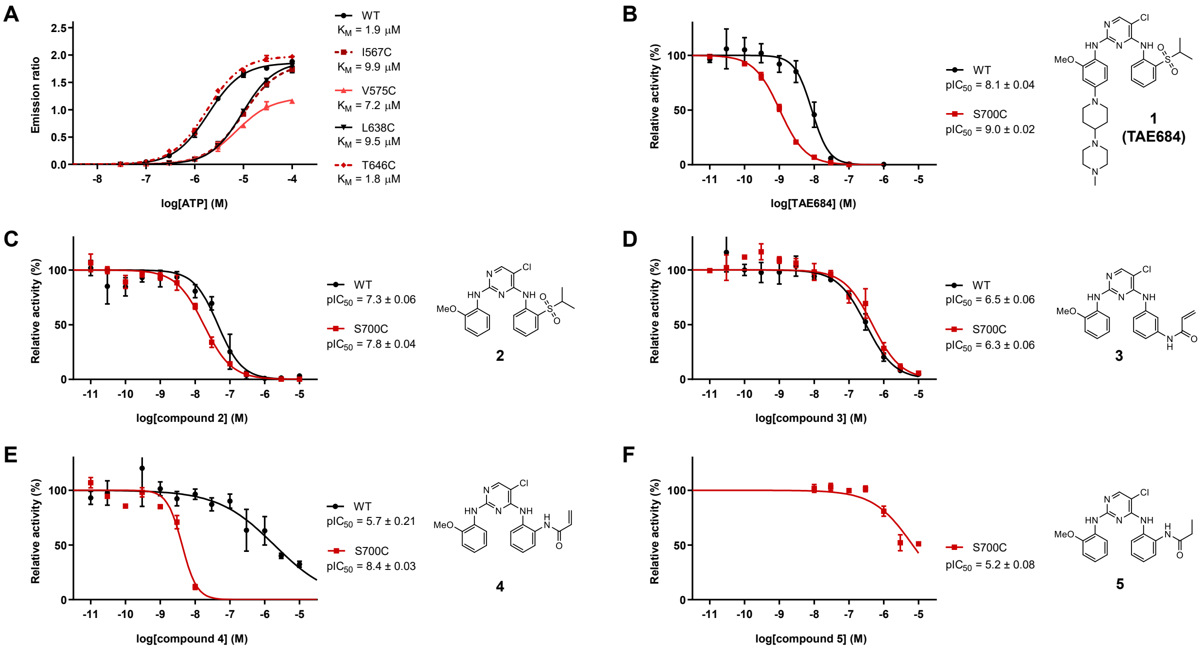


**Figure S1. Determination of ATP $K_M$ for other active FES mutants and concentration-response curves of inhibitors against $FES^{WT}$ and $FES^{S700C}$.**

(**A**) Determination of ATP $K_M$ for $FES^{WT}$ and other FES mutants with >50% relative activity as determined in Fig. 2b.

(**B-F**) Concentration-response curves of inhibitors against $FES^{WT}$ and $FES^{S700C}$ as determined in TR-FRET assay. Compound code and structure is depicted right of the corresponding curve. Data represent means ± SEM (n = 3).

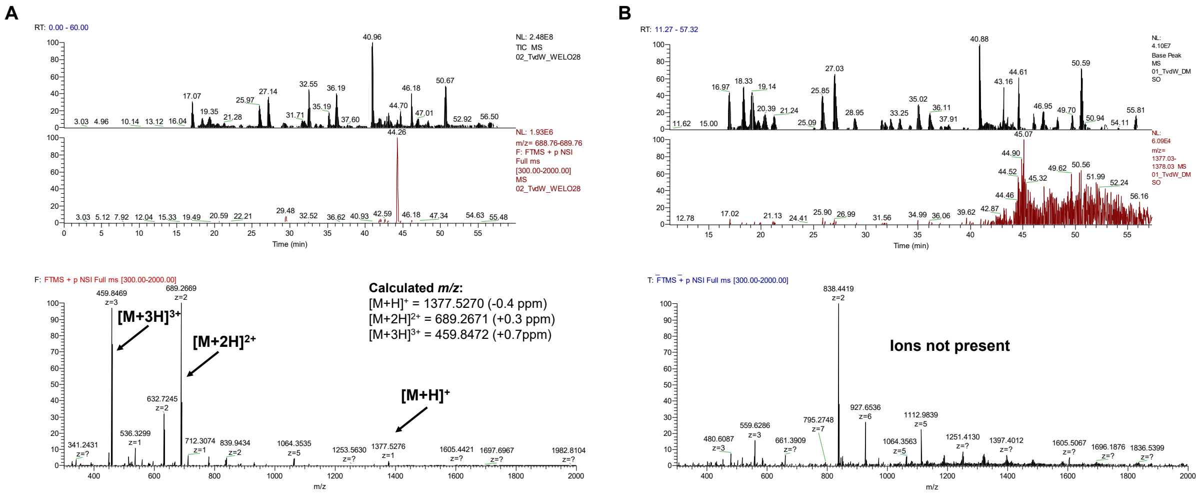


**Figure S2. LC-MS elution profiles of FES$^{S700C}$ incubated with WEL028 (A) or vehicle (B).** Expected precursor ions and corresponding calculated m/z values are indicated. Ions were not present in vehicle-treated control sample.

| Kinase | Inhibition (%) | |
|---|---|---|
| ABL1 | -3 | |
| ABL2 (Arg) | -5 | |
| ACVR1 (ALK2) | 2 | † |
| ACVR1B (ALK4) | 1 | |
| ACVR2B | 9 | † |
| ADRBK1 (GRK2) | 4 | |
| ADRBK2 (GRK3) | 7 | |
| AKT1 (PKB alpha) | -3 | |
| AKT2 (PKB beta) | -1 | |
| AKT3 (PKB gamma) | 1 | |
| AMPK A1/B1/G1 | 13 | |
| AMPK A2/B1/G1 | 4 | |
| AURKA (Aurora A) | 74 | |
| AURKB (Aurora B) | 37 | |
| AURKC (Aurora C) | 37 | |
| AXL | -2 | |
| BLK | 0 | |
| BMPR1A (ALK3) | -2 | † |
| BMX | 16 | |
| BRAF | 94 | * |
| BRSK1 (SAD1) | 2 | |
| BTK | 11 | |
| CAMK1 (CaMK1) | 6 | * |
| CAMK1D (CaMKI delta) | 6 | |
| CAMK2A (CaMKII alpha) | -6 | |
| CAMK2B (CaMKII beta) | 0 | |
| CAMK2D (CaMKII delta) | 8 | |
| CAMK4 (CaMKIV) | -5 | |
| CAMKK1 (CAMKKA) | 22 | † |
| CAMKK2 (CaMKK beta) | 23 | † |
| CDC42 BPA (MRCKA) | -7 | |
| CDC42 BPB (MRCKB) | 6 | |
| CDK1/cyclin B | 1 | |
| CDK2/cyclin A | -3 | |
| CDK5/p25 | 2 | |
| CDK5/p35 | -4 | |
| CDK7/cyclin H/MNAT1 | 6 | |
| CDK8/cyclin C | 0 | † |
| CDK9/cyclin K | 5 | † |
| CDK9/cyclin T1 | 16 | |
| CHEK1 (CHK1) | 0 | |
| CHEK2 (CHK2) | 4 | |
| CHUK (IKK alpha) | 7 | |
| CLK1 | 5 | |
| CLK2 | 1 | |
| CLK3 | 11 | |
| CLK4 | 53 | † |
| CSF1R (FMS) | 11 | |
| CSK | 5 | |
| CSNK1A1 (CK1 alpha 1) | 9 | |

| Kinase | Inhibition (%) | |
|---|---|---|
| CSNK1D (CK1 delta) | 18 | |
| CSNK1E (CK1 epsilon) | 9 | |
| CSNK1G1 (CK1 gamma 1) | 4 | |
| CSNK1G2 (CK1 gamma 2) | 4 | |
| CSNK1G3 (CK1 gamma 3) | 0 | |
| CSNK2A1 (CK2 alpha 1) | 4 | |
| CSNK2A2 (CK2 alpha 2) | 6 | |
| DAPK1 | 12 | |
| DAPK3 (ZIPK) | 4 | |
| DDR1 | -2 | † |
| DDR2 | 3 | † |
| DMPK | 3 | † |
| DNA-PK | 6 | |
| DYRK1A | 1 | |
| DYRK1B | 0 | |
| DYRK3 | 2 | |
| DYRK4 | 0 | |
| EEF2K | 9 | |
| EGFR (ErbB1) | 6 | |
| EPHA1 | 5 | |
| EPHA2 | 7 | |
| EPHA3 | 9 | † |
| EPHA4 | 5 | |
| EPHA5 | 0 | |
| EPHA7 | 4 | † |
| EPHA8 | -7 | |
| EPHB1 | 0 | |
| EPHB2 | 4 | |
| EPHB3 | 1 | |
| EPHB4 | 1 | |
| ERBB2 (HER2) | 2 | |
| ERBB4 (HER4) | 8 | |
| FER | 23 | |
| FES (FPS) | 15 | |
| FGFR1 | 3 | |
| FGFR2 | 7 | |
| FGFR3 K650E | 8 | |
| FGFR3 | 6 | |
| FGFR4 | 34 | |
| FGR | 10 | |
| FLT1 (VEGFR1) | 36 | |
| FLT3 | 61 | |
| FLT4 (VEGFR3) | 120 | |
| FRAP1 (mTOR) | 11 | |
| FRK (PTK5) | 7 | |
| FYN | 4 | |
| GRK4 | -1 | |
| GRK5 | 4 | |
| GRK6 | 0 | |
| GRK7 | 11 | |

| Kinase | Inhibition (%) | |
|---|---|---|
| GSG2 (Haspin) | 6 | |
| GSK3A (GSK3 alpha) | 6 | |
| GSK3B (GSK3 beta) | 4 | |
| HCK | 6 | |
| HIPK1 (Myak) | -4 | |
| HIPK2 | 1 | |
| HIPK3 (YAK1) | -3 | |
| HIPK4 | 14 | |
| IGF1R | 15 | |
| IKBKB (IKK beta) | 1 | |
| IKBKE (IKK epsilon) | 2 | |
| INSR | 11 | |
| INSRR (IRR) | 9 | |
| IRAK1 | 26 | |
| IRAK4 | 0 | |
| ITK | -7 | |
| JAK1 | -3 | |
| JAK2 | 0 | |
| JAK3 | 14 | |
| KDR (VEGFR2) | 67 | |
| KIT | 18 | |
| LCK | 5 | |
| LIMK1 | 14 | † |
| LIMK2 | -2 | † |
| LRRK2 FL | 97 | |
| LRRK2 | 94 | |
| LTK (TYK1) | 43 | |
| LYN A | 5 | |
| LYN B | 1 | |
| MAP2K1 (MEK1) | 110 | * |
| MAP2K2 (MEK2) | 106 | * |
| MAP2K3 (MEK3) | 46 | † |
| MAP2K6 (MKK6) | 73 | * |
| MAP3K10 (MLK2) | 4 | † |
| MAP3K11 (MLK3) | 7 | † |
| MAP3K14 (NIK) | 3 | † |
| MAP3K2 (MEKK2) | 3 | † |
| MAP3K3 (MEKK3) | 7 | † |
| MAP3K5 (ASK1) | 6 | † |
| MAP3K8 (COT) | 77 | * |
| MAP3K9 (MLK1) | -11 | |
| MAP4K2 (GCK) | -12 | |
| MAP4K4 (HGK) | 7 | |
| MAP4K5 (KHS1) | -5 | |
| MAPK1 (ERK2) | 26 | |
| MAPK10 (JNK3) | -7 | * |
| MAPK11 (p38 beta) | 6 | |
| MAPK12 (p38 gamma) | 10 | |
| MAPK13 (p38 delta) | 5 | |
| MAPK14 (p38 alpha) | 7 | |

| Kinase | Inhibition (%) | |
|---|---|---|
| MAPK3 (ERK1) | 19 | |
| MAPK8 (JNK1) | 16 | * |
| MAPK9 (JNK2) | 2 | * |
| MAPKAPK2 | 7 | |
| MAPKAPK3 | 9 | |
| MAPKAPK5 (PRAK) | 66 | |
| MARK1 (MARK) | 1 | |
| MARK2 | 8 | |
| MARK3 | 6 | |
| MARK4 | 8 | |
| MATK (HYL) | 8 | |
| MELK | -2 | |
| MERTK (cMER) | 4 | |
| MET (cMet) | -4 | |
| MINK1 | 20 | |
| MKNK1 (MNK1) | 13 | |
| MKNK2 (MNK2) | 93 | † |
| MLCK (MLCK2) | 9 | † |
| MST1R (RON) | 5 | |
| MST4 | 19 | |
| MUSK | 26 | |
| MYLK (MLCK) | 15 | † |
| MYLK2 (skMLCK) | -3 | |
| NEK1 | 6 | |
| NEK2 | 12 | |
| NEK6 | 9 | |
| NEK7 | -3 | |
| NEK9 | -8 | |
| NLK | 3 | † |
| NTRK1 (TRKA) | 17 | |
| NTRK2 (TRKB) | 5 | |
| NTRK3 (TRKC) | 4 | |
| NUAK1 (ARK5) | 35 | |
| PAK1 | 8 | |
| PAK2 (PAK65) | 29 | |
| PAK3 | 7 | |
| PAK4 | -4 | |
| PAK6 | -7 | |
| PAK7 (KIAA1264) | 12 | |
| PASK | 2 | |
| PDGFRA (PDGFR alpha) | 63 | |
| PDGFRB (PDGFR beta) | 22 | |
| PDK1 Direct | 5 | |
| PDK1 | 36 | |
| PHKG1 | -2 | |
| PHKG2 | 0 | |
| PI4KA (PI4K alpha) | 7 | ** |
| PI4KB (PI4K beta) | 34 | |
| PIK3C2A (PI3K-C2 alpha) | 5 | |
| PIK3C2B (PI3K-C2 beta) | -1 | ** |

| Kinase | Inhibition (%) | |
|---|---|---|
| PIK3C3 (hVPS34) | 11 | |
| PIK3CA/PIK3R1 (p110a/p85a) | 14 | |
| PIK3CD/PIK3R1 (p110d/p85a) | 8 | |
| PIK3CG (p110 gamma) | 13 | |
| PIM1 | -8 | |
| PIM2 | -4 | |
| PKN1 (PRK1) | 7 | |
| PLK1 | 12 | |
| PLK2 | 12 | |
| PLK3 | -1 | |
| PRKACA (PKA) | 2 | |
| PRKCA (PKC alpha) | 14 | |
| PRKCB1 (PKC beta I) | 16 | |
| PRKCB2 (PKC beta II) | 0 | |
| PRKCD (PKC delta) | 14 | |
| PRKCE (PKC epsilon) | -13 | |
| PRKCG (PKC gamma) | 1 | |
| PRKCH (PKC eta) | -4 | |
| PRKCI (PKC iota) | 10 | |
| PRKCN (PKD3) | 31 | |
| PRKCQ (PKC theta) | 8 | |
| PRKCZ (PKC zeta) | -1 | |
| PRKD1 (PKC mu) | 11 | |
| PRKD2 (PKD2) | 3 | |
| PRKG1 | -3 | |
| PRKG2 (PKG2) | 4 | |
| PRKX | -3 | |
| PTK2 (FAK) | 78 | |
| PTK2B (FAK2) | 34 | |
| PTK6 (Brk) | 31 | |
| RET | 7 | |
| RIPK2 | 1 | † |
| ROCK1 | -3 | |
| ROCK2 | 0 | |
| ROS1 | 56 | |
| RPS6KA1 (RSK1) | 5 | |
| RPS6KA2 (RSK3) | 19 | |
| RPS6KA3 (RSK2) | 13 | |
| RPS6KA4 (MSK2) | 1 | |
| RPS6KA5 (MSK1) | -7 | |
| RPS6KA6 (RSK4) | 5 | |
| RPS6KB1 (p70S6K) | 4 | |
| SGK (SGK1) | 10 | |
| SGK2 | 2 | |
| SGKL (SGK3) | -7 | |
| SLK | 4 | † |
| SNF1LK2 | -6 | |
| SPHK1 | 2 | |
| SPHK2 | 3 | ** |
| SRC | 9 | |

| Kinase | Inhibition (%) | |
|---|---|---|
| SRMS (Srm) | 7 | |
| SRPK1 | 7 | |
| SRPK2 | -1 | |
| STK16 (PKL12) | 32 | † |
| STK17A (DRAK1) | 9 | † |
| STK22B (TSSK2) | 18 | |
| STK22D (TSSK1) | 19 | |
| STK23 (MSSK1) | -1 | |
| STK24 (MST3) | 12 | |
| STK25 (YSK1) | 5 | |
| STK3 (MST2) | 5 | |
| STK33 | 60 | † |
| STK4 (MST1) | 3 | |
| SYK | 3 | |
| TAOK2 (TAO1) | 4 | |
| TAOK3 (JIK) | 1 | † |
| TBK1 | 7 | |
| TEC | 5 | † |
| TEK (Tie2) | 7 | |
| TGFBR1 (ALK5) | 6 | † |
| TNK2 (ACK) | 73 | † |
| TXK | 7 | |
| TYK2 | -1 | |
| TYRO3 (RSE) | 5 | |
| WEE1 | -5 | † |
| WNK2 | 5 | † |
| YES1 | 8 | |
| ZAK | 4 | † |
| ZAP70 | 2 | |

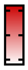


**Figure S3. Single-point selectivity screen of WEL028 on a panel of 279 kinases.** All data were obtained from SelectScreen™ selectivity profiling service. Assays were performed at 1 µM WEL028 with 1 h preincubation. The ATP concentration was equal to the kinase $K_M$, except for those indicated († = LanthaScreen technology, no ATP; * = 100 µM ATP; ** = 10 µM ATP). Values represent mean percentage inhibition compared to vehicle-treated control (n = 2).

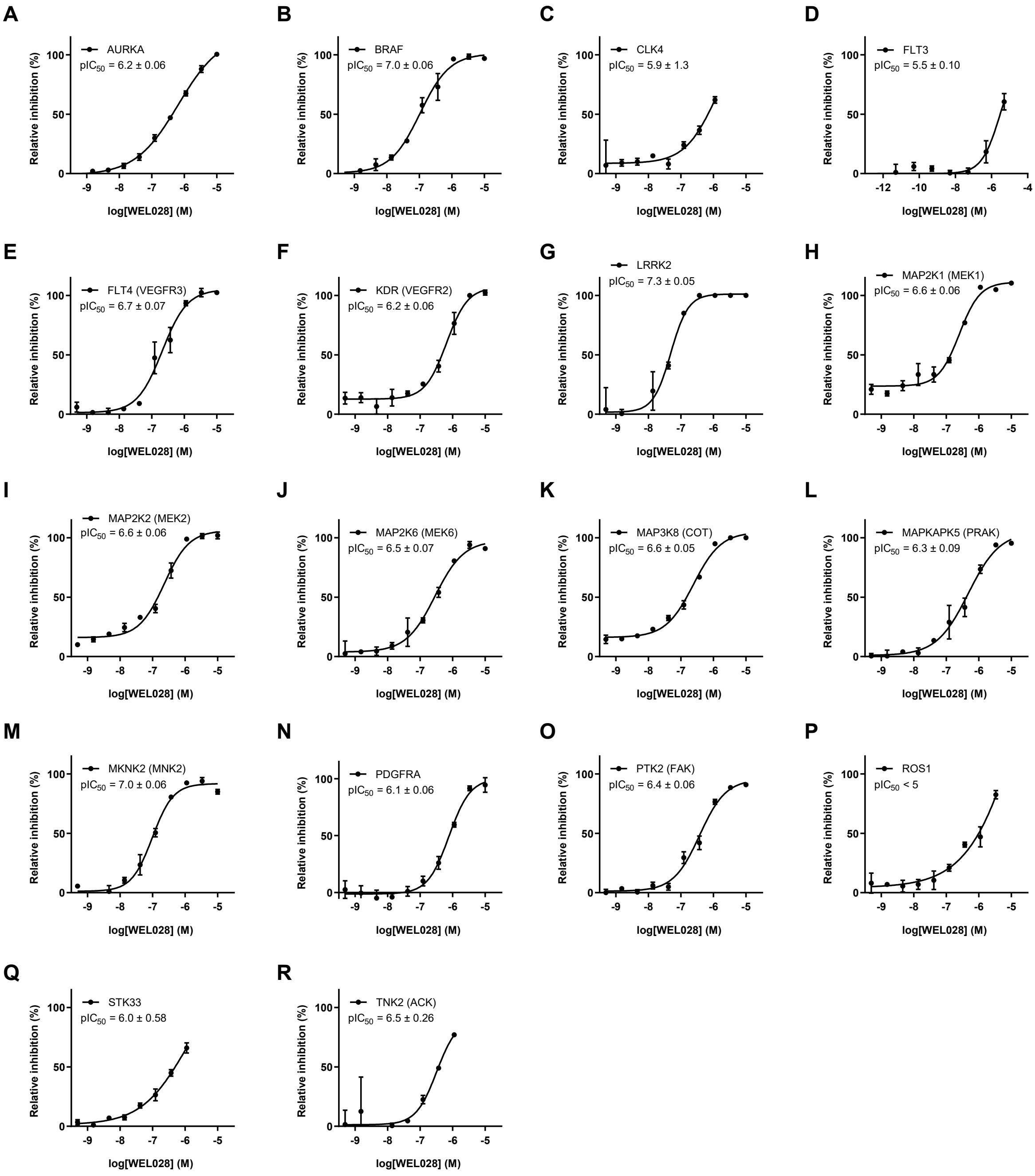


**Figure S4. Concentration-response curves of WEL028 against kinases with >50 % inhibition at 1 μM in initial single-dose screen.** Data (means ± SD, n = 2) were obtained from SelectScreen™ selectivity profiling service. Assays were performed with 1 h preincubation and concentration of ATP was selected to be equal to the $K_M$, unless indicated otherwise in Figure S3.

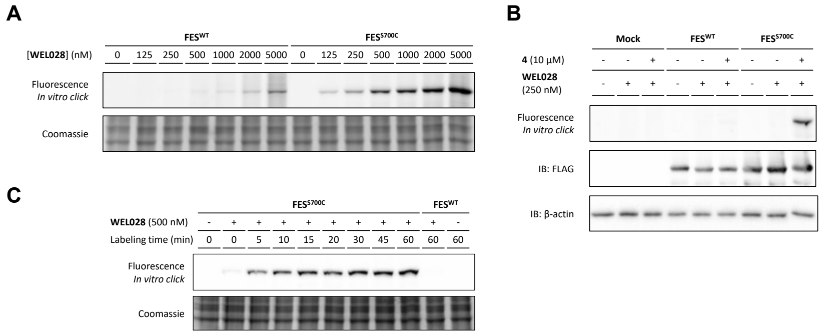


**Figure S5. Two-step labeling of FES$^{S700C}$ by WEL028 conjugated to Cy5-azide *in vitro* using click chemistry.**

(**A**) Dose-dependent labeling of recombinantly expressed full-length FES$^{S700C}$ but not FES$^{WT}$ in HEK293T cell lysate. Lysates were incubated with WEL028 (indicated concentration, 30 min, rt), followed by addition of click mix containing Cy5-azide (2 eq., 30 min, rt). Samples were resolved by SDS-PAGE, followed by in-gel fluorescence scanning.

(**B**) Two-step labeling by WEL028 is specific and exclusive for FES$^{S700C}$. Recombinantly expressed FES in HEK293T cell lysate was preincubated with vehicle or compound **4** (10 µM, 30 min, rt), followed by incubation with two-step probe WEL028 (250 nM, 30 min, rt) and click mix containing Cy5-azide (2 eq., 30 min, rt). Samples were resolved by SDS-PAGE, followed by in-gel fluorescence scanning. Protein expression was verified by immunoblot against a C-terminal FLAG-tag and β-actin as loading control.

(**C**) WEL028 labeling kinetics for FES$^{S700C}$. Lysates were incubated with WEL028 (500 nM, indicated time, rt) and processed as in **A**. Complete labeling was achieved after 15 min and this labeling was stable up to 60 min.

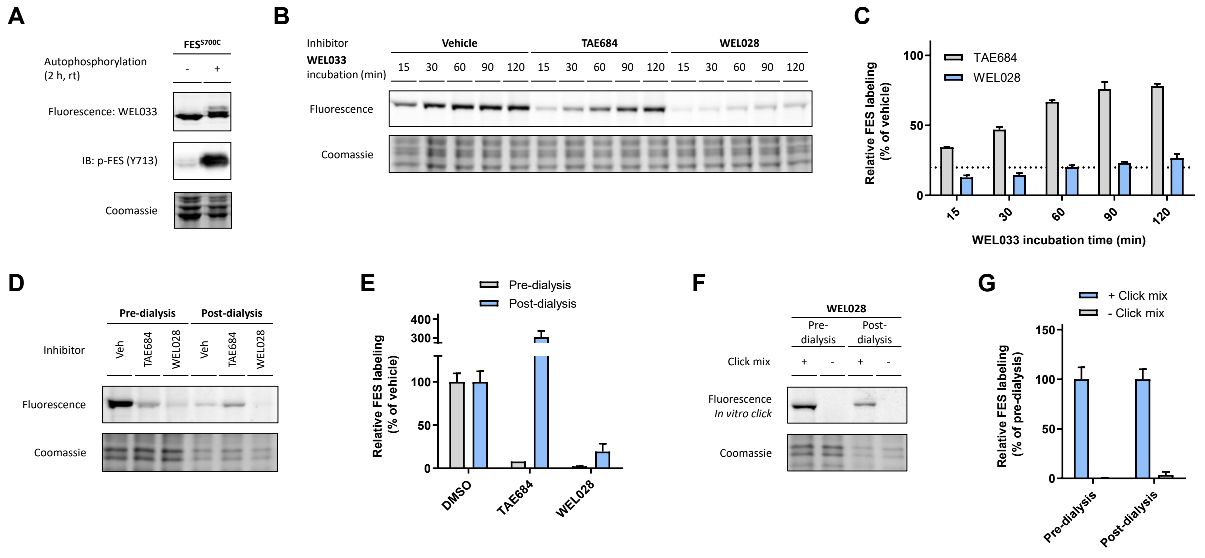


**Figure S6. Characterization of irreversible, covalent binding mode of WEL028 by probe displacement and inhibitor washout experiments.**

(**A**) WEL033 labels both non-autophosphorylated and autophosphorylated $FES^{S700C}$. *E. coli* BL21(DE3) recombinantly co-expressing truncated FESS700C (SH2-KD) and YopH were lysed, after which part of the lysate was subjected to autophosphorylation (2 h, rt). Lysates were labeled with WEL033 (250 nM, 30 min, rt) and analyzed by in-gel fluorescence and immunoblot using anti-phospho-FES Y713 antibody.

(**B**) WEL033 outcompetes TAE684 but not WEL028 binding over time. Lysate was incubated with vehicle, TAE684 or WEL028 at the respective $IC_{80}$-concentration (20% remaining activity; TAE684: 82 nM, WEL028: 27 nM; 30 min, rt), followed by incubation with WEL033 (1 µM, indicated time, rt).

(**C**) Quantification of band intensity as shown in panel **B**, normalized to vehicle-treated control at same time point.

(**D**) Sustained $FES^{S700C}$ inhibition by WEL028 but not TAE684 after overnight dialysis. Lysates were treated with vehicle, TAE684 or WEL028 as in panel **B** and pre-dialysis samples were directly flash-frozen after incubation. Residual lysate was dialyzed overnight at 4°C. Pre- and post-dialysis samples were then treated with WEL033 (250 nM, 30 min, rt).

(**E**) Quantification of band intensity as shown in panel **D**, normalized to vehicle-treated control.

(**F**) Two-step labeling of WEL028-bound $FES^{S700C}$ before and after dialysis. Samples were processed as in panel **D**, but conjugated to BODIPY-azide using click chemistry (2 eq., 30 min, rt).

(**G**) Quantification of band intensity as shown in panel **F**, normalized to pre-dialysis control. Data represent means ± SEM (n = 3).

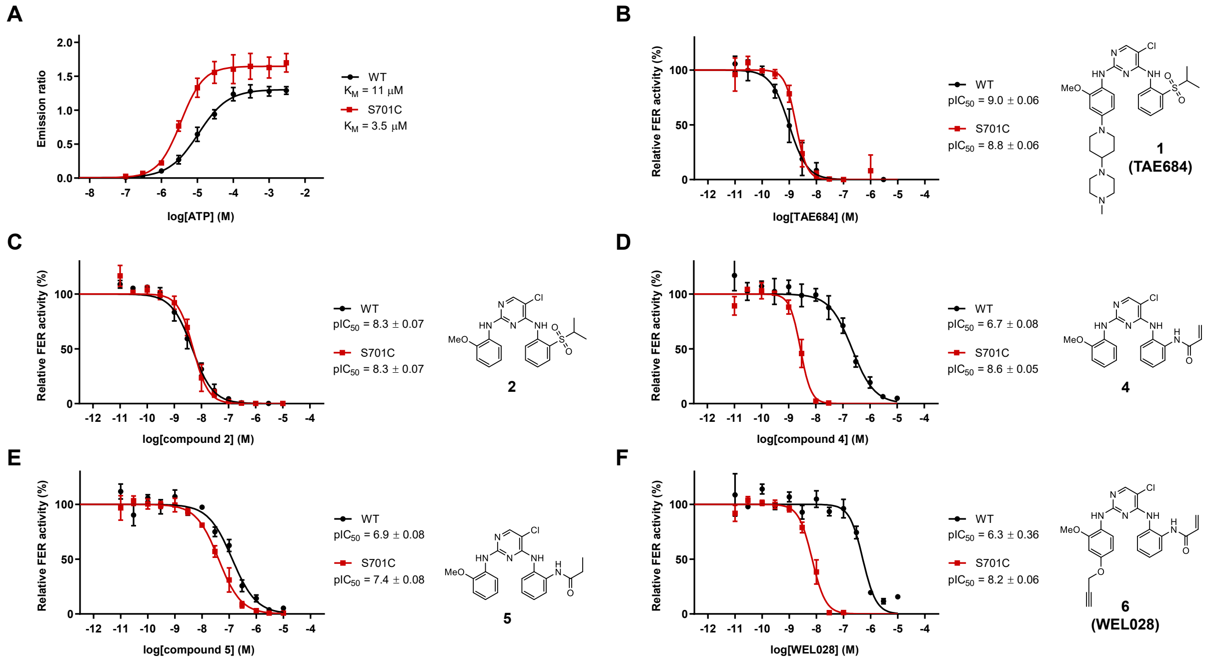


**Figure S7. Determination of ATP $K_M$ for $FER^{WT}$ and $FER^{S701C}$ and concentration-response curves of inhibitors against $FER^{WT}$ and $FER^{S701C}$.**

(**A**) Determination of ATP $K_M$ for $FER^{WT}$ and $FER^{S701C}$.

(**B-F**) Concentration-response curves of inhibitors against $FER^{WT}$ and $FER^{S701C}$ as determined in TR-FRET assay. Final ATP concentration was 12 μM and 1 μM for $FER^{WT}$ and $FER^{S701C}$, respectively. Compound code and structure is depicted right of the corresponding curve. Data represents means ± SEM (n = 3).

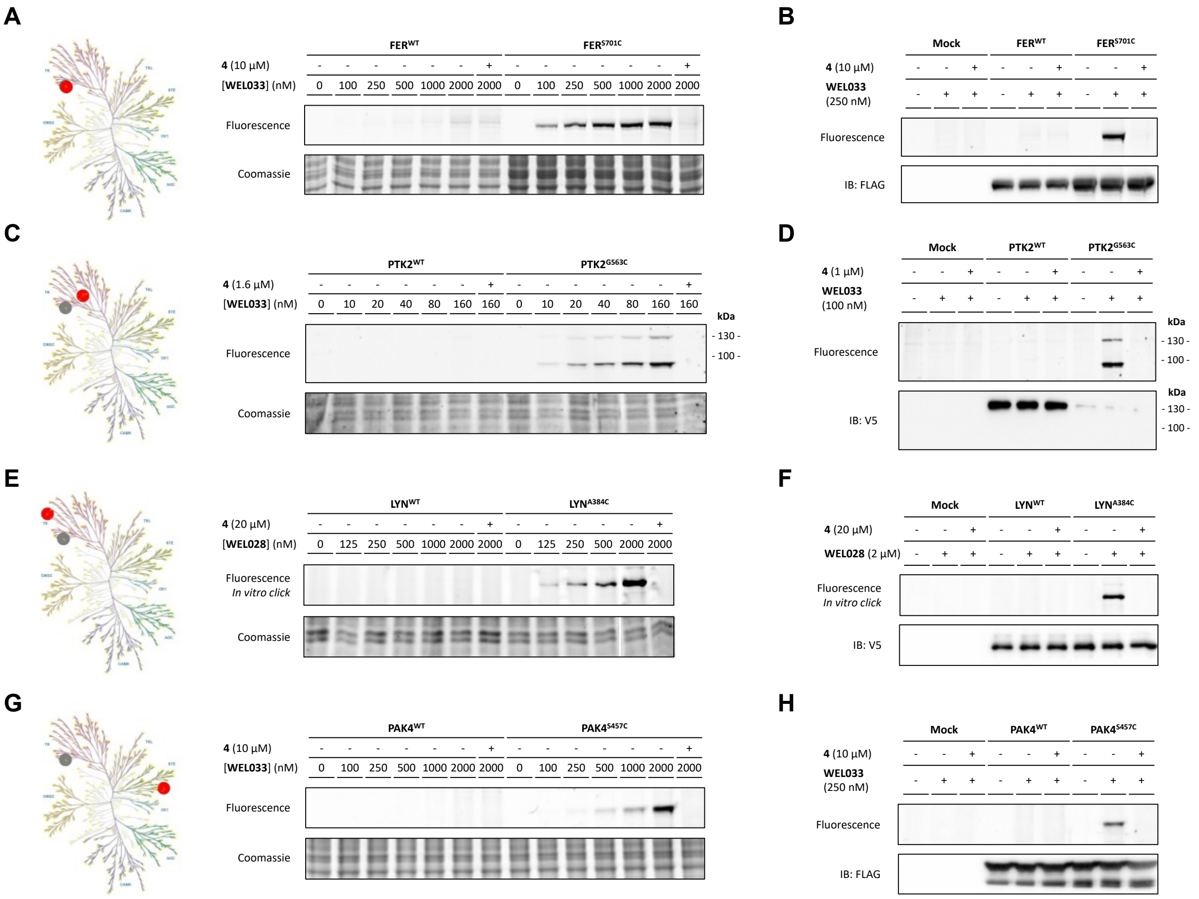

**Figure S8. Applicability of chemical genetic strategy on various kinases harboring DFG-1 residues mutated into cysteines.**

(**A**, **C**, **E**, **G**) Dose-dependent and specific labeling of DFG-1 cysteine mutants but not wild-type kinases by complementary probes. Recombinantly expressed kinase (**A**: FER, **C**: PTK2, **E**: LYN, **G**: PAK4) in HEK293T cell lysate were preincubated with vehicle or **4** (indicated concentration, 30 min, rt), followed by incubation with probe (**A**, **C**, **G**: one-step probe WEL033, **E**: two-step probe WEL028; indicated concentrations, 30 min, rt). For **E**, samples were then conjugated to Cy5-azide using click chemistry. Homology-based similarity of corresponding kinase (red) to FES (gray) is visualized in kinome tree illustrations.

(**B**, **D**, **F**, **H**) Labeling by complementary probes is specific and exclusive for DFG-1 cysteine mutants. Samples were treated as aforementioned, but at the optimal probe concentration (indicated, 30 min, rt). Protein expression was verified by immunoblot against a C-terminal FLAG-tag or V5-tag. Of note, PTK2 migrated a two bands of which only the upper band was detected by immunoblot. Kinome illustrations were rendered using KinMap (www.kinhub.org/kinmap), reproduced courtesy of Cell Signaling Technology, Inc. www.cellsignal.com).

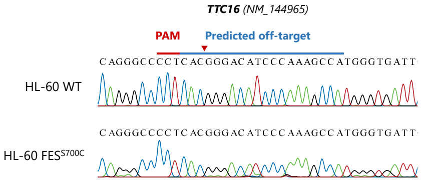


**Figure S9. Analysis of putative sgRNA off-target site.** Analysis of the only predicted coding off-target of the sgRNA target sequence employed for FES$^{S700C}$ mutagenesis, located in the *TTC16* gene. Genomic region surrounding putative off-target site was amplified by PCR, followed by Sanger sequencing analysis. No off-target gene editing events were observed.

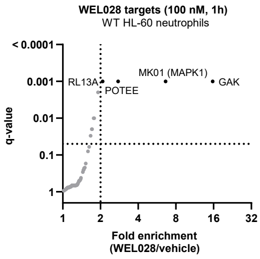


**Figure S10. Chemical proteomics-based identification of WEL028 kinase targets at 100 nM in WT HL-60 neutrophils.** Kinases with > 2-fold enrichment compared to vehicle control (q < 0.05) were designated as targets. Values represent means of fold enrichment (n = 3).

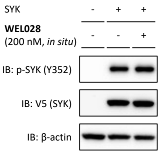


**Figure S11. WEL028 does not affect SYK Y352 phosphorylation in U2OS cells in absence of FES$^{S700C}$.** U2OS cells were transfected with only V5-tagged SYK. After 48 h, cells were incubated with vehicle or WEL028 (200 nM, 1 h) and lysed. Lysates were incubated with WEL033 (250 nM, 30 min, rt) and analyzed by in-gel fluorescence and immunoblot (n = 3).

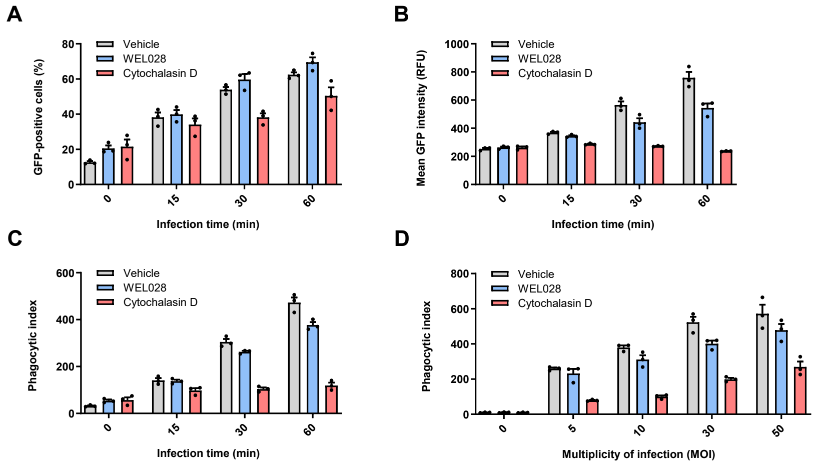


**Figure S12. Phagocytic uptake of *E. coli* by HL-60 FES$^{S700C}$ neutrophils at variable multiplicity of infection (MOI) and infection time.**

(**A-C**) HL-60 FES$^{S700C}$ neutrophils were incubated with vehicle, WEL028 (100 nM) or Cytochalasin D (10 µM) for 1 h, after which GFP-expressing E. coli were added at MOI of 30. After indicated times, cells were washed and fixed (1% PFA, 15 min, 4 °C), followed by flow cytometry analysis. Phagocytic index (**C**) was calculated as fraction of GFP-positive cells (number of phagocytic cells, **A**) multiplied by GFP MFI (number of phagocytized bacteria, **B**).

(**D**) Neutrophils were treated as in panel **A-C**, but with a variable MOI for 1 h. Data represent means ± SEM (N = 3).

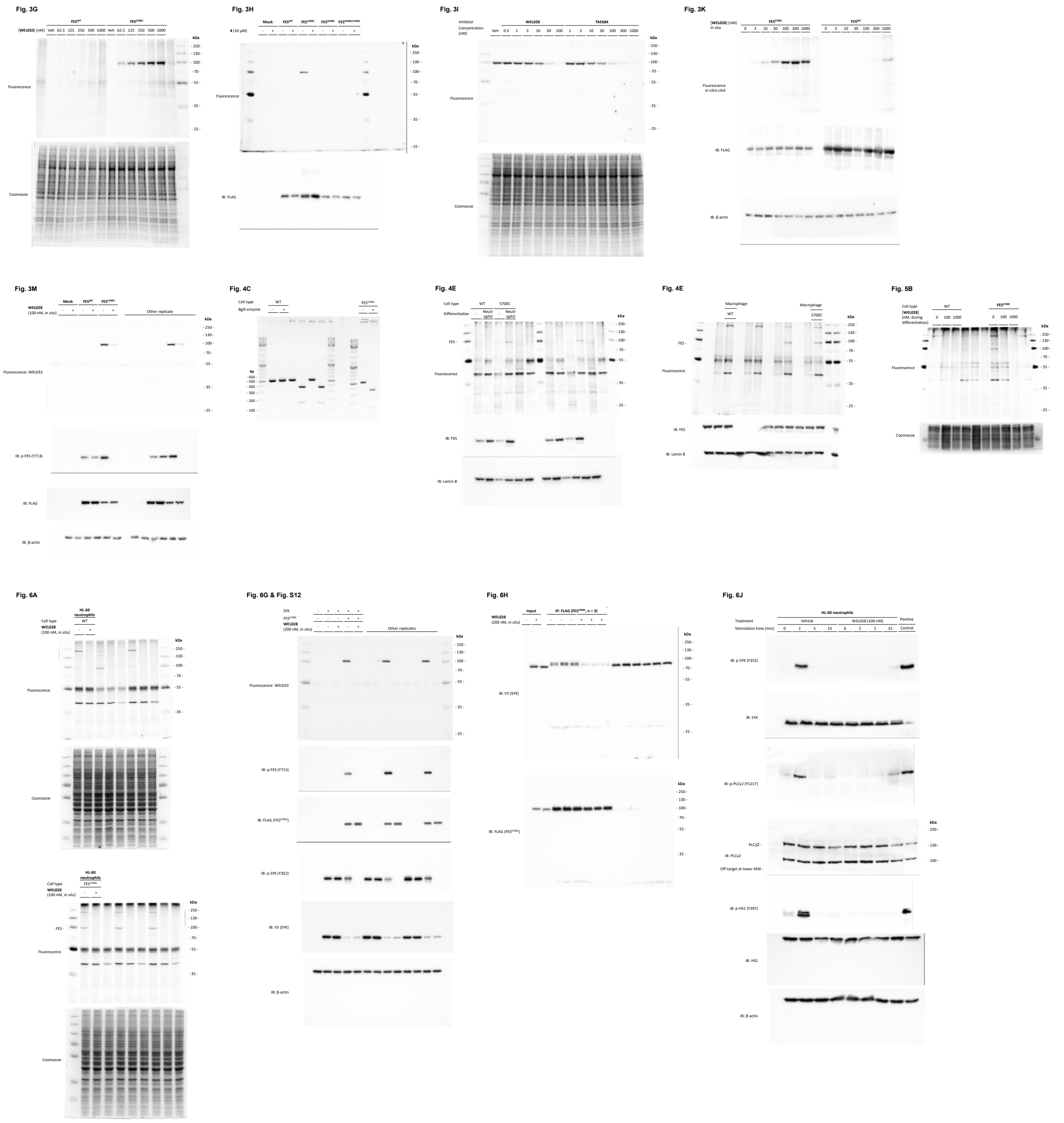


**Figure S13. Uncropped gel and blot images (continues on next page).** Unmarked lanes are not part of this study. Membranes were generally cut in two and developed as separate immunoblots.

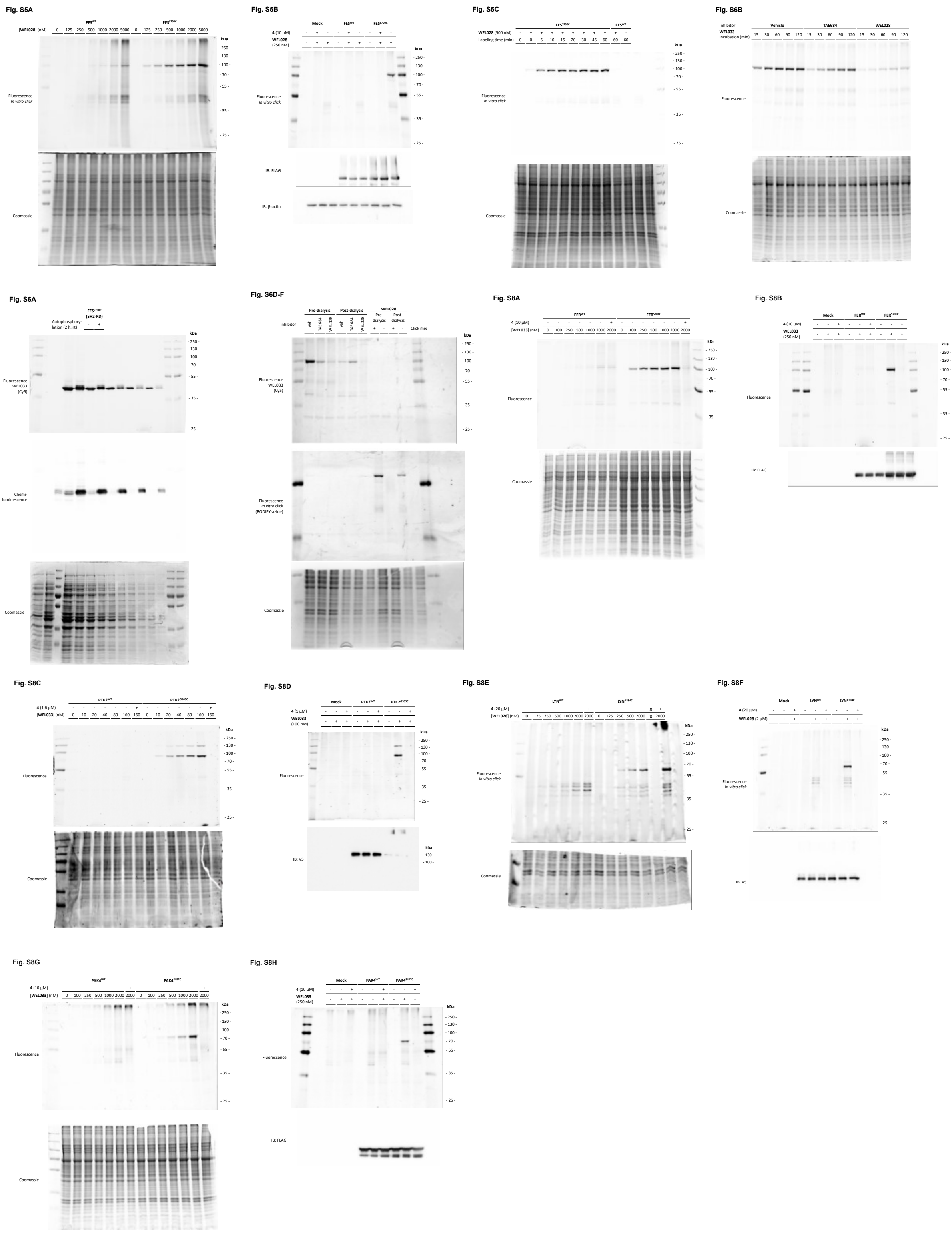


**Figure S13. Uncropped gel and blot images (continued).**

**Table S1: Substrate identification using PamChip® activity assay.** Top 30 of peptides with highest signal intensity are shown. Numbers behind substrate names indicate the amino acid residue numbers within the corresponding protein sequence. Predicted phosphorylated tyrosine residues are indicated in bold red.

| Substrate | Peptide | log2 of signal intensity | |
|---|---|---|---|
| | | $FES^{WT}$ | $FES^{S700C}$ |
| CD79A_181_193 | EYEDENL**Y**EGLNL | 14.59 | 14.18 |
| ENOG_37_49 | SGASTGI**Y**EALEL | 14.39 | 14.13 |
| **ZAP70_313_325** | SVYESP**Y**SDPEEL | 12.83 | 12.51 |
| EFS_246_258 | GGTDEGI**Y**DVPLL | 12.68 | 12.43 |
| PLCG1_764_776 | IGTAEPD**Y**GALYE | 12.56 | 12.48 |
| RET_1022_1034 | TPSDSLI**Y**DDGLS | 12.53 | 12.29 |
| IRS2_626_638 | HPYPED**Y**GDIEIG | 12.38 | 12.20 |
| EPHA1_774_786 | LDDFDGT**Y**ETQGG | 12.35 | 12.24 |
| PGFRB_572_584 | VSSDGHE**Y**IYVDP | 12.33 | 12.09 |
| P85A_600_612 | NENTEDQ**Y**SLVED | 12.27 | 12.19 |
| LAT_249_261 | EEGAPD**Y**ENLQEL | 12.14 | 12.23 |
| PLCG2_1191_1203_C1200S | ESEEEL**Y**SSSRQL | 12.14 | 12.04 |
| PDPK1_2_14 | ARTTSQL**Y**DAVPI | 12.10 | 11.97 |
| FRK_380_392 | KVDNEDI**Y**ESRHE | 11.99 | 11.88 |
| PTN11_57_67 | QNTGD**Y**YDLYG | 11.89 | 11.89 |
| PTN11_580_590 | SARV**Y**ENVGLM | 11.71 | 11.71 |
| KIT_930_942_C942S | ESTNHI**Y**SNLANS | 11.68 | 11.59 |
| PDPK1_369_381 | DEDCYGN**Y**DNLLS | 11.55 | 11.57 |
| EGFR_1165_1177 | ISLDNPD**Y**QQDFF | 11.40 | 11.59 |
| PECA1_708_718 | DTETV**Y**SEVRK | 11.36 | 11.53 |
| PGFRB_709_721 | RPPSAEL**Y**SNALP | 11.35 | 11.53 |
| EPHA7_607_619 | TYIDPET**Y**EDPNR | 11.30 | 11.43 |
| VGFR2_989_1001 | EEAPEDL**Y**KDFLT | 11.26 | 11.38 |
| PGFRB_1002_1014 | LDTSSVL**Y**TAVQP | 11.25 | 11.33 |
| PTN6_558_570 | KHKEDV**Y**ENLHTK | 11.18 | 11.16 |
| EPHA2_765_777 | EDDPEAT**Y**TTSGG | 11.16 | 11.32 |
| PTN6_531_541 | GQESE**Y**GNITY | 11.15 | 11.29 |
| FES_706_718 | REEADGV**Y**AASGG | 11.07 | 11.21 |
| FER_707_719 | RQEDGGV**Y**SSSGL | 10.98 | 11.13 |
| IRS1_890_902 | PKSPGE**Y**VNIEFG | 10.93 | 11.11 |

**Table S2: SH2 binding partner identification using PamChip® binding assay.** Top 30 of peptides with highest signal intensity are shown. Numbers behind substrate names indicate the amino acid residues within the corresponding protein.

| Binding partner | Peptide | log2 of signal intensity | |
|---|---|---|---|
| | | $FES^{WT}$ | $FES^{S700C}$ |
| PGFRB_572_584 | VSSDGHEYIYVDP | 12.44 | 13.06 |
| PGFRB_1014_1028 | PNEGDNDYIIPLPDP | 12.17 | 12.94 |
| LAT_249_261 | EEGAPDYENLQEL | 10.79 | 12.19 |
| VGFR2_1168_1180 | AQQDGKDYIVLPI | 10.69 | 11.64 |
| ENOG_37_49 | SGASTGIYEALEL | 10.67 | 12.29 |
| PTN11_57_67 | QNTGDYYDLYG | 10.66 | 12.05 |
| CD3E_182_194 | PVPNPDYEPIRKG | 10.03 | 11.48 |
| CD79A_181_193 | EYEDENLYEGLNL | 10.02 | 12.05 |
| IRS1_890_902 | PKSPGEYVNIEFG | 9.97 | 11.28 |
| LAT_194_206 | MESIDDYVNVPES | 9.51 | 11.14 |
| MAPK3_198_210_C203S | ALQTPSYTPYYVA | 9.32 | 10.68 |
| MK12_180_189_M182B | SEBTGYVVTR | 9.16 | 10.58 |
| PTN6_558_570 | KHKEDVYENLHTK | 8.74 | 10.88 |
| FGFR2_762_774 | TLTTNEEYLDLSQ | 8.26 | 9.86 |
| TYK2_1048_1060 | VPEGHEYYRVRED | 7.88 | 9.60 |
| JAK3_974_986 | LPLDKDYYVVREP | 7.87 | 9.77 |
| RON_1346_1358 | SALLGDHYVQLPA | 7.75 | 9.04 |
| FGFR3_753_765 | TVTSTDEYLDLSA | 7.66 | 9.57 |
| MET_1227_1239 | RDMYDKEYYSVHN | 7.59 | 9.42 |
| PTN11_580_590 | SARVYENVGLM | 7.27 | 9.94 |
| EGFR_1190_1202 | STAENAEYLRVAP | 7.12 | 9.01 |
| MK14_173_185 | RHTDDEMTGYVAT | 7.08 | 9.11 |
| FAK2_572_584 | RYIEDEDYYKASV | 7.00 | 9.11 |
| JAK1_1027_1039 | AIETDKEYYTVKD | 6.53 | 8.78 |
| MK03_199_208 | GFLTEYVATR | 6.49 | 8.31 |
| EPOR_419_431 | ASAASFEYTILDP | 6.42 | 7.97 |
| MK12_178_190 | ADSEMTGYVVTRW | 6.15 | 8.13 |
| 41_654_666 | LDGENIYIRHSNL | 6.08 | 8.02 |
| MK07_212_224 | AEHQYFMTEYVAT | 4.96 | 6.25 |
| HAVR2_257_267 | GIRSEENIYTI | 4.95 | 4.67 |

**Table S3: Inhibitory potency of WEL028 against kinases with >50 % inhibition at 1 μM in initial single-dose screen.** All data ($pIC_{50}$ ± SD, n = 2) were obtained from SelectScreen™ selectivity profiling service except for $FES^{S700C}$, which was determined in-house. Kinases exhibiting >50% inhibition at 1 μM in an initial screen on 279 kinases (Figure S3) were selected for dose-response experiments. Assays were performed with 1 h preincubation. Indicated molecular weight is based on UniProt database records. Location of native cysteine residues in the kinase active site is indicated if applicable, with nomenclature as previously described.[1] Apparent fold selectivity was calculated as $IC_{50}$ on that kinase divided by $IC_{50}$ on $FES^{S700C}$.

| Kinase | Molecular weight (kDa) | Native cysteine | $pIC_{50}$ | Apparent fold selectivity |
|---|---|---|---|---|
| AURKA | 45.8 | None | 6.2 ± 0.06 | 155 |
| BRAF | 84.4 | Hinge2 | 7.0 ± 0.06 | 25 |
| CLK4 | 57.5 | None | 5.9 ± 1.3 | 272 |
| **$FES^{S700C}$** | **93.5** | **DFG-1** | **8.4 ± 0.03** | **N/A** |
| FLT3 | 112.9 | DFG-1 | 5.5 ± 0.10 | 718 |
| FLT4 (VEGFR3) | 152.8 | DFG-1, Hinge2 | 6.7 ± 0.07 | 50 |
| KDR (VEGFR2) | 151.5 | DFG-1, Hinge2 | 6.2 ± 0.06 | 153 |
| LRRK2 | 286.1 | None | 7.3 ± 0.05 | 12 |
| MAP2K1 (MEK1) | 43.4 | DFG-1, GK-1 | 6.6 ± 0.06 | 60 |
| MAP2K2 (MEK2) | 44.4 | DFG-1, GK-1 | 6.6 ± 0.06 | 54 |
| MAP2K6 (MEK6) | 37.5 | DFG-1, GK-1 | 6.5 ± 0.07 | 68 |
| MAP3K8 (COT) | 52.9 | DFG+1 | 6.6 ± 0.05 | 57 |
| MAPKAPK5 (PRAK) | 54.2 | DFG-1 | 6.3 ± 0.09 | 112 |
| MKNK2 (MNK2) | 51.9 | DFG-1 | 7.0 ± 0.06 | 23 |
| PDGFRA | 122.7 | DFG-1, Hinge2 | 6.1 ± 0.06 | 184 |
| PTK2 (FAK) | 119.2 | Hinge2 | 6.4 ± 0.06 | 89 |
| ROS1 | 263.9 | None | < 5 | > 238 |
| STK33 | 57.8 | Hinge2 | 6.0 ± 0.58 | 254 |
| TNK2 (ACK) | 114.6 | None | 6.5 ± 0.26 | 76 |

**Table S4: Inhibitory potency of synthesized TAE684 derivatives against FER$^{WT}$ and FER$^{S701C}$.** Half maximal inhibitory concentrations (expressed as $pIC_{50}$) were determined using recombinantly expressed FER$^{WT}$ and FER$^{S701C}$ in a TR-FRET assay. Final ATP concentration was 12 µM and 1 µM for FER$^{WT}$ and FER$^{S701C}$, respectively. Apparent fold selectivity was calculated as $IC_{50}$ on FER$^{WT}$ divided by $IC_{50}$ on FER$^{S701C}$. Data represent means ± SEM (n = 3). ND: not determined. Dose-response curves can be found in Figure S7.

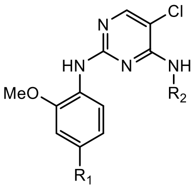


| Compound | $R_1$ | $R_2$ | $pIC_{50}$ Fer$^{WT}$ | $pIC_{50}$ Fer$^{S701C}$ | Apparent fold selectivity |
|---|---|---|---|---|---|
| **1** (**TAE684**) | 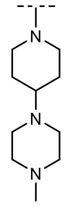 | 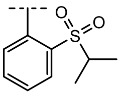 | 9.0 ± 0.06 | 8.8 ± 0.06 | 0.54 |
| **2** | H | 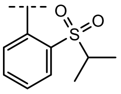 | 8.3 ± 0.07 | 8.3 ± 0.07 | 0.92 |
| **4** | H | 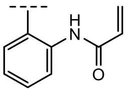 | 6.7 ± 0.08 | 8.6 ± 0.05 | 81 |
| **5** | H | 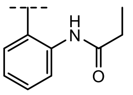 | 6.9 ± 0.08 | 7.4 ± 0.08 | 3.2 |
| **6** (**WEL028**) | 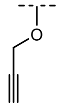 | 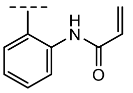 | 6.3 ± 0.06 | 8.2 ± 0.06 | 74 |

**Table S5: List of putative off-target cleavage sites for sgRNA employed in CRISPR/Cas9-mediated mutagenesis of FES.** Specificity of sgRNA was assessed using DESKGEN™ online web tool (www.deskgen.com). Sites with 3 or less mismatches compared to the sgRNA target were included. Only 1 putative off-target is located in the coding region of a gene.

| Potential off-target sequence | PAM | Similarity | Mismatches | Gene | Locus |
|---|---|---|---|---|---|
| **TGGCTTTGGGATGTCCCGTG** | **AGG** | **5** | **3,19** | **Yes** | **chr9@ 127716868-127716891** |
| TGACTTTGGCATGTCCTGAG | AGG | 3 | 10,17 | No | chr16@ 50371486-50371509 |
| TTAGTTTGGGATGTCCAGAG | GGG | 1 | 2,4,17 | No | chr2@ 147874374-147874397 |
| TGCCTTTGGTATGTCCAGAG | TAG | 1 | 3,10,17 | No | chr13@ 25630797-25630820 |
| AGACTTTGGGAGGTCCCGTG | CAG | 1 | 1,12,19 | No | chr12@ 11447626-11447649 |
| TGGCTGTGGGATGTCCCCAG | GAG | 0 | 3,6,18 | No | chr3@ 13295903-13295926 |
| TGAGTCTGGGATGTCCCTAG | AGG | 0 | 4,6,18 | No | chr5@ 137202156-137202179 |
| TGGCTTTGGGATGTCGGGAG | AAG | 0 | 3,16,17 | No | chr20@ 61773235-61773258 |
| TGAATTTGGGATGTCCCATG | TAG | 0 | 4,18,19 | No | chr5@ 135589616-135589639 |
| TGAATTTGGGATGCCCTGAG | AGG | 0 | 4,14,17 | No | chr5@ 61639561-61639584 |
| TGACGTTGGGATGACCAGAG | CAG | 0 | 5,14,17 | No | chr19@ 35077359-35077382 |
| TGACTTTGGGTTGTCCCCAT | GAG | 0 | 11,18,20 | No | chrX@ 112142232-112142255 |
| TGACTTTGGGGTCTCCCAAG | AGG | 0 | 11,13,18 | No | chr2@ 95278649-95278672 |

| Mismatches | 0 | 1 | 2 | 3 |
|---|---|---|---|---|
| Coding | 0 | 0 | **1** | 0 |
| Non-coding | 0 | 0 | 1 | 12 |

### Table S6: Complete list of oligonucleotide sequences.

| ID | Application | Name | Sequence |
|---|---|---|---|
| P1 | Cloning | FES_SH2-KD_forw | AGGGCGCCATGGGGATTCCGGAGGTGCAGAAGC |
| P2 | Cloning | FES_SH2-KD_rev | CACTCGAGCACCGCGGCCGCTTACCGATGCCGCTTTCGGAT |
| P3 | Cloning | FES_I567C_forw | GTGTTGGGTGAGCAGTGTGGACGGGGGAACTTT |
| P4 | Cloning | FES_G570C_forw | GAGCAGATTGGACGGTGCAACTTTGGCGAAGTG |
| P5 | Cloning | FES_V575C_forw | GGGAACTTTGGCGAATGCTTCAGCGGACGCCTG |
| P6 | Cloning | FES_L638C_forw | TACATCGTCATGGAGTGTGTGCAGGGGGGCGAC |
| P7 | Cloning | FES_G642C_forw | GAGCTTGTGCAGGGGTGCGACTTCCTGACCTTC |
| P8 | Cloning | FES_T646C_forw | GGGGGCGACTTCCTGTGCTTCCTCCGCACGGAG |
| P9 | Cloning | FES_N688C_forw | GACCTGGCTGCTCGGTGCTGCCTGGTGACAGAG |
| P10 | Cloning | FES_L690C_forw | GCTGCTCGGAACTGCTGCGTGACAGAGAAGAAT |
| P11 | Cloning | FES_S700C_forw | AATGTCCTGAAGATCTGTGACTTTGGGATGTCC |
| P12 | Cloning | FES_forw | CTTAAGCTTTGGTACCGCCGCCACCATGGGCTTCTCTTCTGAGC |
| P13 | Cloning | FES_rev | CATTCTAGATCACTCGAGACCGGTCCGATGCCGCTTTCGGAT |
| P14 | Cloning | FES_K590E_forw | ACCCTGGTGGCGGTGGAGTCTTGTCGAGAGACG |
| P15 | Cloning | FES_K590E_rev | CGTCTCTCGACAAGACTCCACCGCCACCAGGGT |
| P16 | Cloning | FER_SH2-KD_forw | CGTCTCCCATGATCTCCATCAGTGAGAAGCCTT |
| P17 | Cloning | FER_SH2-KD_rev | CACCGCGGCCGCTTATGTGAGTTTTCTCTTGAT |
| P18 | Cloning | FER_S701C_forw | AATGTTCTGAAAATCTGTGACTTTGGAATGT |
| P19 | Cloning | FER_S701C_rev | ACATTCCAAAGTCACAGATTTTCAGAACATT |
| P20 | Cloning | FER_forw | AGCCGTCTCGGTACCGCCGCCACCATGGGGTTTGGGAGTGACC |
| P21 | Cloning | FER_rev | CATTCTAGATCACTCGAGACCGGTTGTGAGTTTTCTCTTGA |
| P22 | Cloning | LYN_A384C_forw | GTCACTCATGTGCAAGATCTGTGATTTTGGCCTTGCT |
| P23 | Cloning | LYN_A384C_rev | AGCAAGGCCAAAATCACAGATCTTGCACATGAGTGAC |
| P24 | Cloning | PTK2_G563C_forw | GATTGTGTAAAATTATGCGACTTTGGTCTCTCCCGATATATGGAA |
| P25 | Cloning | PTK2_G563C_rev | TTCCATATATCGGGAGAGACCAAAGTCGCATAATTTTACACAATC |
| P26 | Cloning | PAK4_forw | CTTAAGCTTTGGTACCGCCGCCACCATGTTTGGGAAGAGGAAGAA |
| P27 | Cloning | PAK4_rev | CATTCTAGATCACTCGAGACCGGTTCTGGTGCGGTTCTGGCGCA |
| P28 | Cloning | PAK4_S457C_forw | GGCAGGGTGAAGCTGTGTGACTTTGGGTTCTGC |
| P29 | Cloning | PAK4_S457C_rev | GCAGAACCCAAAGTCACACAGCTTCACCCTGCC |
| P30 | CRISPR mutagenesis | sgRNA_hFES-MUT_S700C_top | CACCTGACTTTGGGATGTCCCGAG |
| P31 | CRISPR mutagenesis | sgRNA_hFES-MUT_S700C_bott | AAACCTCGGGACATCCCAAAGTCA |
| P32 | CRISPR mutagenesis | gPCR_hFES_S700C_forw | TTTTGTCCTTGGCTTTCCTAGA |
| P33 | CRISPR mutagenesis | gPCR_hFES_S700C_rev | GTGCTTACCCTTCTCCACAAAC |
| P34 | CRISPR off-target analysis | HDR-template_hFES_S700C | ACTGTTGGCCAAATGAGCCCCTGCCCTGTCTCACCCAGGGACCTGGCTGCTCGGAACTGCCTG<br>GTGACAGAGAAGAATGTGCTGAAGATCTGTGACTTTGGCATGTCCCGAGAAGAAGCCGATG<br>GGGTCTATGCAGCCTCAGGGGGCCTCAGACAAGTCCCCGTGAAGTGGACCGCACCTGAGGCC<br>CTTAACTA |
| P35 | CRISPR off-target analysis | gPCR_TTC16_forw | AGAACAGACGGTGTTGTAAGCAT |
| P36 | CRISPR off-target analysis | gPCR_TTC16_rev | ATTAGACAGTTGAGTTCACTGAGGC |

**Supplementary Materials & Methods – Biology**

### 1. General

All chemicals were purchased at Sigma Aldrich, unless stated otherwise. DNA oligos were purchased at Sigma Aldrich or Integrated DNA Technologies and sequences can be found in Table S6. Cloning reagents were from Thermo Fisher. TAE684 and R406 were purchased at Selleckchem and Cytochalasin D and U-73122 at Focus Biomolecules. Cy5-azide and BODIPY-azide were previously synthesized in-house and characterized by NMR and LC-MS[2]. All cell culture disposables were from Sarstedt. Bacterial and eukaryotic protease inhibitor cocktails were obtained from Amresco.

### 2. Cloning

Full-length human cDNA encoding FES and PAK4 was obtained from Source Bioscience. pDONR223-constructs with full-length human cDNA of FER, LYN, PTK2 and SYK were a gift from William Hahn & David Root (Addgene Human Kinase ORF Collection). For bacterial expression constructs, human FES cDNA encoding residues 448-822 or human FER cDNA encoding residues 448-820 was amplified by PCR and cloned into expression vector pET1a in frame of an N-terminal $His_6$-tag and Tobacco Etch Virus (TEV) recognition site. Eukaryotic expression constructs of FES, FER and PAK4 were generated by PCR amplification and restriction/ligation cloning into a pcDNA3.1 vector, in frame of a C-terminal FLAG-tag. Eukaryotic expression constructs of LYN, PTK2 and SYK were generated using Gateway™ recombinational cloning into a pcDest40 vector, in frame of a C-terminal V5-tag, according to recommended procedures (Thermo Fisher). Point mutations were introduced by site-directed mutagenesis and all plasmids were isolated from transformed XL-10 competent cells (prepared using *E. coli* transformation buffer set; Zymo Research) using plasmid isolation kits following the supplier's protocol (Qiagen). All sequences were verified by Sanger sequencing (Macrogen).

For CRISPR/Cas9 plasmids, guides were cloned into the BbsI restriction site of plasmid px330-U6-Chimeric_BB-CBh-hSpCas9 (gift from Feng Zhang, Addgene plasmid #42230) as previously described[3,4].

### 3. Protein expression and purification

Bacterial expression constructs were transformed into *E. coli* BL21(DE3) co-transformed with a pCDFDuet-1 vector encoding *Yersinia* phosphatase YopH (kindly provided by prof. dr. Kuriyan)[5]. Cells were grown in Luria Broth (LB) medium containing 50 µg/mL kanamycin and 50 µg/mL streptomycin at 37°C to an $OD_{600}$ of 0.4. The cultures were cooled on ice and protein expression was induced by addition of 50 µM isopropyl-β-D-thiogalactopyranoside (IPTG) for 16 h at 18°C. Cultures (typically 10 mL per mutant) were centrifuged (1000 *g*, 10 min, 4°C) and washed in 1 mL physiological salt solution (0.9% (w/v) NaCl). The pellet was then resuspended in 400 µL lysis buffer (100 mM $NaH_2PO_4$ pH 8.0, 500 mM NaCl, 10% glycerol, 0.5 mM tris(2-carboxyethyl)phosphine (TCEP), 20 mM imidazole, 1 x bacterial protease inhibitor cocktail) and cells were lysed by sonication on ice (15 cycles of 4'' on, 9.9'' off at 25% maximum amplitude; Vibra Cell (Sonics)). $MgCl_2$ and OmniCleave (Epicentre) were added to final concentrations of 10 mM and 125 U/mL respectively, and lysates were incubated for 10 min at rt.

Meanwhile, 200 µL Nickel Magnetic Beads (BiMake), were homogenized by vortexing and washed in lysis buffer (3 x 200 µL) using a magnetic separator. Lysate was added to beads and incubated for 1 h at 4°C with vigorous shaking. Beads were then washed with lysis buffer containing 75 mM imidazole (3 x 400 µL), after which the beads were transferred to a clean Eppendorf tube. Protein was phosphorylated on beads by addition of autophosphorylation buffer (50 mM HEPES pH 7.5, 500 mM NaCl, 5% glycerol, 10 mM dithiothreitol (DTT), 10 mM $Na_3VO_4$, 2 mM $(NH_4)_2SO_4$, 10 mM $MgCl_2$, 5 mM $MnCl_2$, 2 mM ATP) and vigorous shaking for 2 h at rt. Beads were subsequently washed with lysis buffer containing 20 mM imidazole (3 x 400 µL), after which the protein was eluted in lysis buffer containing 250 mM imidazole (2 x 200 µL). The elution fractions were combined, applied onto a 10 kDa cutoff centrifugal filter unit (Amicon) and centrifuged (14,000 *g*, 10 min, 4°C). The retentate was reconstituted in 100 µL protein storage buffer (10 mM HEPES pH 7.5, 500 mM NaCl, 5% glycerol, 10 mM DTT), aliquoted and stored at -80°C. Protein concentration was measured using Qubit fluorometric quantitation (Thermo Fisher) and protein purity was monitored by SDS-PAGE and Coomassie staining.

### 4. In vitro TR-FRET kinase assay

Assays were performed in white ProxiPlate-384 Plus™ 384-well microplates (Perkin Elmer). Incubation steps were performed at 21°C in kinase reaction buffer (50 mM HEPES pH 7.5, 150 mM NaCl, 10 mM $MgCl_2$, 1 mM EGTA, 2 mM DTT, 0.01% Tween-20). Purified proteins were diluted in kinase reaction buffer prior to use. All measurements were performed in triplicate.

Plates were read on a TECAN Infinite M1000 Pro plate reader, using fluorescence top reading settings ($\lambda_{ex}$ = 320/20 nm, $\lambda_{em,donor}$ = 615/10 nm, $\lambda_{em,acceptor}$ = 665/10 nm, 100 µs integration time, 50 µs lag time). Emission ratios were calculated as fluorescence of acceptor / fluorescence of donor. *Z'*-factors were determined for each individual assay plate as $Z' = 1 - 3(\sigma_{pc} + \sigma_{nc})/(\mu_{pc} - \mu_{nc})$, with σ = standard deviation and µ = mean of measured replicates, wild-type or untreated samples as positive control (pc) and samples incubated with no ATP as negative control (nc). All plates met the requirement of $Z' > 0.7$. Emission ratios were corrected for background signal of samples incubated with no ATP. Corrected ratios were averaged and normalized to wild-type or untreated signal as 100% reference.

For relative activity determination of mutants, purified wild-type or mutant FES (12.5 ng per well) or FER (5 ng per well) was incubated with 50 nM U*Light*-TK peptide (Perkin Elmer) and 100 µM ATP for 60 min at 21°C in a total volume of 10 µL. The reaction was quenched by addition of 10 µL development solution (20 mM EDTA, 4 nM Europium-anti-phosphotyrosine antibody (PT66, Perkin Elmer) and incubated for 60 min before fluorescence was measured.

For $K_M$ determinations, assay was performed as described above, but with variable ATP concentrations in a dilution series of 1 mM to 100 nM. For $IC_{50}$ determinations, serial dilutions of inhibitor were prepared in DMSO, followed by further dilution in kinase reaction buffer. The inhibitors were premixed with peptide and ATP (5 µM final concentration, unless indicated otherwise), after which wild-type or mutant kinase was added to initiate the reaction. The final DMSO concentration during the reaction was 1%. $K_M$ and $IC_{50}$ curves were fitted using GraphPad Prism® 7 (GraphPad Software Inc.).

### 5. PamChip® microarray assay

*Kinase activity assay*

Kinase activity profiles were determined using the PamChip® 12 protein tyrosine (PTK) peptide microarray system (PamGene International B.V., ‘s-Hertogenbosch, The Netherlands) according to the instructions of the manufacturer, essentially as described[6] with the exception that arrays were blocked with 2% BSA and the assay buffer contained EDTA instead of EGTA. Sample input was 0.25 ng purified FES (wild-type or S700C) per array and [ATP] = 400 µM. For arrays with inhibitor, recombinant FES was preincubated in assay mix without ATP with vehicle or WEL028 (100 nM, 30 min, on ice, 2% final DMSO concentration).

*SH2 domain binding assay*

PamChip® protein tyrosine kinase (PTK) arrays were blocked by pumping a 2% BSA solution up and down through the array (30 µL, 15 min, 2 cycles/min). After three washing steps, the arrays were pre-phosphorylated by incubation with 1.5 pmol of ($His_6$-tagged) JAK2 catalytic domain for 60 cycles in kinase assay buffer with 0.01% BSA. After 3 washing steps, the arrays were incubated with 0.75 µg of FES (wild-type or S700C) in PBS/0.1% Tween supplemented with 0.01% BSA. Incubations without FES served as negative control. After 3 washing steps, binding of FES was visualized by incubation with Alexa Fluor 488 labeled anti-penta-His antibody (Qiagen, #1019199), 1 µL per array, in PBS/0.01% Tween. Peptide phosphorylation of JAK2 was detected with the same anti p-Tyr antibody as used in the PTK activity assay. After incubation for 30 min (2 cycles/min), arrays were washed, and images taken at different exposure times.

*Data analysis and quality control*

Data quantification of the images at all exposure times and reaction times and visualization of the data were performed using BioNavigator software (PamGene International B.V., ‘s-Hertogenbosch, The Netherlands). Post-wash signals (local background subtracted) were used. After signal quantification and integration of exposure times, signals were log2-transformed for visualization. Peptides with no ATP-dependent signal were excluded from analysis. Identification of peptides that were significantly different between conditions was performed using a Mixed Model statistical analysis. Substrate consensus motif was generated using Enologos (http://www.benoslab.pitt.edu).

**5. Selectivity profiling**

Selectivity profiling assays were performed by the Invitrogen SelectScreen™ Services. A complete list of tested kinases can be found in Figure S3, detailed assay procedures are described in SelectScreen Assay Conditions documents located at www.invitrogen.com/kinaseprofiling. All kinases were preincubated for 1 h at indicated concentrations of inhibitor. The concentration of ATP was selected to be equal to the $K_M$, unless indicated otherwise. An initial screen was performed on 279 kinases at single dose of 1 µM. All available kinases with native cysteine residues in the active site were included in this panel. Kinases showing >50% inhibition at 1 µM were further profiled in dose-response experiments in a 10-point dilution series of 10 µM to 0.5 nM.

### 6. Docking studies

All structure-based modeling was performed in Schrödinger Suite 2017-2 (Schrödinger). The docking of compound **4** was based on the crystal structure of FES co-crystallized with TAE684 (PDB: 4e93)[7], which was prepared by the protein preparation wizard. Prior to docking, the Ser700 residue was manually changed into a cysteine. Subsequently, compound **4** was aligned to TAE684 on the basis of the diaminopyrimidine (both ligands share the same kinase/hinge binding moiety). This pose was optimized using an exhaustive hierarchical optimization procedure available in Prime[8]. The acrylamide warhead was found in proximity of Cys700 and the ligand was then covalently attached to this residue, followed by another round of hierarchical optimization[8]. Figures were rendered using PyMOL Molecular Graphics System (Schrödinger).

### 7. MS identification of covalent inhibitor-peptide adduct

Purified $FES^{S700C}$ (1.2 µg in 38 µL) was treated with vehicle or WEL028 (2 µL of 20x concentrated stock in DMSO, 1 µM final concentration) for 30 min at rt. The reaction was quenched by addition of 3x Laemmli buffer (20 µL), incubated for 5 min at 95°C and sample (400 ng, 20 µL) was resolved on 10% acrylamide SDS-PAGE gel (200 V, 60 min). Gel was stained using Coomassie Brilliant Blue R-250, bands were cut out of gel into small blocks and then destained in 500 µL of 50% MeOH in 100 mM $NH_4HCO_3$ pH 8.0 for 10 min at rt. Acetonitrile (500 µL) was added for gel blocks dehydration, after which gel blocks were digested with sequencing-grade trypsin (Promega) in 250 µL trypsin buffer (100 mM Tris pH 7.5, 100 mM NaCl, 1 mM $CaCl_2$, 10% acetonitrile) overnight at 37°C with vigorous shaking. The pH was adjusted with formic acid to pH 3, after which the sample was diluted in extraction solution (65% acetonitrile/35% MilliQ) and concentrated in a SpeedVac concentrator. Samples were subsequently desalted using stage tips with $C_{18}$ material and processed as previously described[9].

Tryptic peptides were analyzed on a Surveyor nano-LC system (Thermo) hyphenated to a LTQ-Orbitrap mass spectrometer (Thermo) as previously described[10]. General mass spectrometric conditions were: electrospray voltage of 1.8 kV, no sheath and auxiliary gas flow, ion transfer tube temperature 150°C, capillary voltage 41 V, tube lens voltage 150 V. Internal mass calibration was performed with polydimethylcyclosiloxane (m/z = 445.12002) and dioctyl phthalate ions (m/z = 391.28429) as lock mass. Samples of 10 µl were separated via a trap-elute setup at 250 nL/min flow and analyzed by data-dependent acquisition of one full scan/ top 3 method. For shotgun proteomics, fragmented precursor ions measured twice within 10 s were dynamically excluded for 60 s and ions with $z < 2$ or unassigned charges were not analyzed. Peptide ID was determined with the Mascot search engine. A parent ion list of the m/z ratios of the active-site peptides was compiled and used for LC-MS/MS analysis in a data-dependent protocol. The parent ion was electrostatically isolated in the ion trap of the LTQ and fragmented by MS/MS. Data from MS/MS experiments were validated manually.

### 8. Inhibitor washout by dialysis

$FES^{S700C}$-overexpressing HEK293T lysate (396 µL, 1.43 mg/mL) was incubated with vehicle, TAE684 or WEL028 (4 µL of 100x concentrated stock in DMSO) at final concentrations corresponding to their respective $IC_{80}$ values (TAE684: 225 nM, WEL028: 231 nM) for 30 min at rt. One fraction (200 µL) of the sample was immediately flash-frozen and stored at -80°C until use. The remaining sample was transferred to a dialysis cassette (Slide-A-Lyzer™ Dialysis Cassette, 7K MWCO, 0.5 mL; Thermo Fisher), followed by dialysis in 200 mL PBS overnight at 4°C. Pre- and post-dialysis samples were then incubated with probe WEL033 as described in section 12, or conjugated to BODIPY-$N_3$ by CuAAC as described in section 13.

### 9. Cell culture

*General cell culture*

Cell lines were purchased at ATCC and were tested on regular basis for mycoplasma contamination. Cultures were discarded after 2-3 months of use. HEK293T (human embryonic kidney) and U2OS (human osteosarcoma) cells were cultured at 37°C under 7% $CO_2$ in DMEM containing phenol red, stable glutamine, 10% (v/v) high iron newborn calf serum (Seradigm), penicillin and streptomycin (200 µg/mL each; Duchefa). Medium was refreshed every 2-3 days and cells were passaged two times a week at 80-90% confluence. HL-60 (human promyeloblast) cells were cultured at 37°C under 5% $CO_2$ in HEPES-supplemented RPMI containing phenol red, stable glutamine, 10% (v/v) fetal calf serum (Biowest), penicillin and streptomycin (200 µg/mL each), unless stated otherwise. Cell density was maintained between 0.2 x $10^6$ and 2.0 x $10^6$ cells/mL. Cell viability was assessed by Trypan Blue exclusion and quantification using a TC20™ Automated Cell Counter (Bio-Rad).

*Transfection*

One day prior to transfection, HEK293T or U2OS cells were transferred from confluent 10 cm dishes to 15 cm dishes. Before transfection, medium was refreshed (13 mL). A 3:1 (m/m) mixture of polyethyleneimine (PEI; 60 µg/dish) and plasmid DNA (20 µg/dish) was prepared in serum-free medium and incubated for 15 min at rt. The mixture was then dropwisely added to the cells, after which the cells were grown to confluence in 72 h (HEK293T) or 48 h (U2OS). Cells were then harvested by suspension in PBS, followed by centrifugation for 5 min at 200 *g*. Cell pellets were flash-frozen in liquid nitrogen and stored at -80°C until sample preparation.

Transfection of CRISPR plasmid DNA and ssODN repair template into HL-60 cells was performed using Amaxa nucleofector kit V and nucleofector I device (Lonza). One day prior to transfection, HL-60 cells were diluted to a density of 0.4 x $10^6$ cells/mL. The next day, 2 x $10^6$ cells per condition were centrifuged at 200 *g* for 5 min and resuspended in 100 µL nucleofection solution. Plasmid DNA (2 µg) and if applicable ssODN repair template (400 pmol) were added and cells were nucleofected using program T-019. Cells were allowed to recover for 10 min at rt before transfer to 12-well plates and further recovery in antibiotics-free medium at 37°C.

*Differentiation of HL-60 cells*

One day prior to induction of differentiation, cells were diluted to 0.4 x $10^6$ cells/mL. Monocyte/macrophage differentiation was induced by addition of phorbol 12-myristate 13-acetate (PMA; 16 nM) for 48 h, during which cells attached to the plastic surface and acquired monocyte/macrophage morphology. For morphological inspection, images were taken on an EVOS FL Auto 2 Imaging System (Thermo Fisher) at 20x magnification. Neutrophil differentiation was induced by addition of all-trans-retinoic acid (1 μM) and dimethylsulfoxide (1.25%) for 72 – 96 h.

*Inhibitor treatment in live cells*

The term *in situ* is used to designate experiments in which live cell cultures are treated with inhibitor, whereas the term *in vitro* refers to experiments in which the inhibitor is incubated with cell lysates. Compounds were diluted in growth medium from a 1000x concentrated stock solution in DMSO.

For *in situ* assays on live transfected cells, cells were transfected prior to treatment as described above. After 48 h, cells were treated with compound for 1 h. Cells were collected by suspension in PBS and centrifuged (1000 *g*, 5 min, rt). Pellets were flash-frozen in liquid nitrogen and stored at -80°C until use.

For *in situ* treatment post-differentiation, HL-60 cells were differentiated as described above and incubated with compound for 1 h unless specified otherwise. Cells were collected by suspension for non-adherent cells and trypsinization for adherent cells. After collection, cells were centrifuged (200 *g*, 5 min, rt) and washed in equal volume of PBS (1000 *g*, 5 min, rt). Cell pellets were flash-frozen in liquid nitrogen and stored at -80°C until use.

**10. CRISPR/Cas9 gene editing**

*sgRNA selection and ssODN homology-directed repair (HDR) donor design*

Selection of sgRNA was based on proximity to desired site of cleavage or mutagenesis as well as efficiency and specificity as predicted by CHOPCHOP v2 online web tool (http://chopchop.cbu.uib.no)[11]. sgRNA sequence can be found in Table S5. A sgRNA targeting exon 16 was selected and an ssODN HDR donor was designed that after successful HDR incorporates the desired S700C mutation along with a BglII restriction site to facilitate analysis by RFLP. Furthermore, silent mutations are incorporated to remove “NGG” protospacer adjacent motifs (PAMs), preventing recleavage of the genomic sequence after successful HDR or cleavage of the ssODN donor itself.

*Single cell isolation, expansion and genotyping*

For preparation of conditioned RPMI medium, HL-60 cells were diluted to 0.2 x $10^6$ cells/mL and grown for 48 h at 37°C. Cell suspension was then centrifuged (1,000 *g*, 5 min) and medium was transferred to a clean tube, followed by a second centrifugation step (3,500 *g*, 10 min). The medium was subsequently filtered through a 0.2 μm sterile filter and stored at -80°C until use, for no longer than 3 months.

Single cell cultures were obtained approximately 7 days post-nucleofection by dilution in 1:1 conditioned and fresh RPMI medium and expanded in 96-well plates (100 μL per well). After 14 days, plates were inspected for cell growth, clones were collected in new plates and half of the volume was transferred to 96-well PCR-plates, followed by centrifugation (1,000 *g*, 10 min). Medium was removed and cell pellets were suspended in 25 μL QuickExtract™ (Epicentre). The samples were incubated at 65°C for 6 min, mixed by vortexing and then incubated at 98°C for 2 min. Genomic DNA extracts were diluted in sterile water and directly used in PCR reactions. Genomic PCR reactions were performed on 5 µL isolated genomic DNA extract using Phusion High-Fidelity DNA Polymerase (Thermo Fisher) in Phusion HF buffer in a final volume of 20 µL.

Restriction Fragment Length Polymorphism (RFLP) assays were performed by combining genomic PCR product (8.5 µL) with FastDigest buffer (1 µL) and FastDigest BglII (0.5 µL; Thermo Fisher), followed by incubation at 37°C for 30 min. Digested PCR products were analyzed using agarose gel electrophoresis with ethidium bromide staining.

For sequencing analysis, genomic PCR products were purified using NucleoSpin® Gel and PCR Clean-up kit (Macherey-Nagel), followed by Sanger sequencing. The efficiency for obtaining homozygous mutant clones was approximately 0.5-1% (typically 1 out of 100-200 clones).

*Off-target analysis*

Potential off-target cleavage sites of the used sgRNA were predicted using DESKGEN™ online web tool (www.deskgen.com). Top-ranked potential coding off-target sequences containing 3 or less mismatches to the employed sgRNA sequence were selected for validation. The genomic region surrounding the potential off-target site was amplified by PCR and analyzed by Sanger sequencing.

### 11. Preparation of cell lysates

*Lysates of overexpressing HEK293T and U2OS cells*

Pellets were thawed on ice and suspended in lysis buffer (50 mM HEPES pH 7.2, 150 mM NaCl, 1 mM $MgCl_2$, 0.1% (w/v) Triton X-100, 2 mM $Na_3VO_4$, 20 mM NaF, 1 x mammalian protease inhibitor cocktail, 25 U/mL benzonase). Cells were lysed by sonication on ice (15 cycles of 4'' on, 9.9'' off at 25% maximum amplitude). Protein concentration was determined using Quick Start™ Bradford Protein Assay (Bio-Rad) and diluted to appropriate concentration in dilution buffer (50 mM HEPES pH 7.2, 150 mM NaCl). Lysates were aliquoted, flash-frozen and stored at -80°C until use.

*Lysates of HL-60 cells*

Pellets were thawed on ice and suspended in M-PER buffer supplemented with 1x Halt™ phosphatase and protease inhibitor cocktail (Thermo Fisher), followed by centrifugation (14,000 *g*, 10 min, 4°C). The resulting clear lysate was further processed as described above.

### 12. One-step probe labeling experiments

For *in vitro* inhibition experiments, cell lysate (14 µL) was preincubated with inhibitor (0.5 µL, 29 x concentrated stock in DMSO, 30 min, rt), followed by incubation with WEL033 (0.5 µL, 30 x concentrated stock in DMSO, 30 min, rt). For *in situ* inhibition experiments, treated cell lysate (14.5 µL) was directly incubated with WEL033 (0.5 µL, 30 x concentrated stock in DMSO, 30 min, rt). Final concentrations of inhibitors and/or WEL033 are indicated in the main text and figure legends. Reactions were quenched with 4x Laemmli buffer (5 µL, final concentrations 60 mM Tris pH 6.8, 2% (w/v) SDS, 10% (v/v) glycerol, 5% (v/v) β-mercaptoethanol, 0.01% (v/v) bromophenol blue) and boiled for 5 min at 95°C. Samples were resolved on 10% acrylamide SDS-PAGE gel (180 V, 75 min). Gels were scanned using Cy3 and Cy5 multichannel settings (605/50 and 695/55 filters, respectively; ChemiDoc™ MP System, Bio-Rad). Fluorescence intensity was corrected for protein loading determined by Coomassie Brilliant Blue R-250 staining and quantified with Image Lab (Bio-Rad). $IC_{50}$ curves were fitted with Graphpad Prism® 7 (Graphpad Software Inc.).

### 13. Two-step probe labeling experiments

For *in vitro* experiments, cell lysate (12 µL) was preincubated with inhibitor (0.5 µL, 25x concentrated stock in DMSO, 30 min, rt), followed by incubation with WEL028 (0.5 µL, 26x concentrated stock in DMSO, 30 min, rt). Meanwhile, "click mix" was prepared freshly by combining $CuSO_4$ (1 µL of 15 mM stock), sodium ascorbate (0.6 µL of 150 mM stock), THPTA (0.2 µL of 15 mM stock) and fluorophore-azide (0.2 µL of 150x concentrated stock in DMSO, 2 eq.). Click mix was added to the reaction, followed by incubation for 30 min at rt, after which the reaction was quenched and further processed as described above.

For *in situ* inhibition experiments, WEL028-treated cells were lysed and directly incubated with click mix.

### 14. Proteomics

Proteomics was performed based on previously described procedures.[12] In summary, full cell lysates of $FES^{WT}$ and $FES^{S700C}$ HL-60 macrophages or neutrophils incubated *in situ* with vehicle or WEL028 were prepared as aforementioned. Lysates (250 µL of 2 mg/mL) were conjugated to biotin-azide using click chemistry (25 µL click mix, 1 h, 37°C). The reaction was quenched and excess biotin-azide was removed by chloroform/methanol precipitation. Precipitated proteome was suspended in 6 M urea in 25 mM ammonium bicarbonate, reduced (10 mM DTT, 15 min, 65°C) and alkylated (40 mM iodoacetamide, 30 min, rt, in the dark). SDS was added (2% final concentration, 5 min, 65°C), samples were diluted in PBS and incubated with avidin beads (from 50% slurry, 3 h, rt, in overhead rotator). Beads were washed with 0.5% SDS in PBS, followed by 3 washes with PBS, and then transferred to low-binding Eppendorf tubes. Proteins were digested with trypsin overnight at 37°C and resulting peptides were desalted using stage tips with $C_{18}$ material as described in section 7. Samples were analyzed on an LC-IMS-MS system with a Synapt G2-Si instrument (Waters) as previously described [12]. Data processing was performed with ISOQuant software as previously described.[13,14] The following cut-offs were used for target identification: unique peptides ≥ 1, identified peptides ≥ 2, ratio WEL028-treated over vehicle-treated ≥ 2 with q-value < 0.05 based on *t*-test with Benjamini-Hochberg multiple comparison correction (FDR = 10%), kinase annotation in Uniprot database.

### 15. Immunoblot

Samples were resolved by SDS-PAGE as described above, but transferred to 0.2 µm polyvinylidene difluoride membranes by Trans-Blot Turbo™ Transfer system (Bio-Rad) directly after fluorescence scanning. Membranes were washed with TBS (50 mM Tris pH 7.5, 150 mM NaCl) and blocked with 5% milk in TBS-T (50 mM Tris pH 7.5, 150 mM NaCl, 0.05% Tween-20) for 1 h at rt.

Membranes were then incubated with primary antibody in 5% milk in TBS-T (FLAG, V5, β-actin; o/n at 4°C) or washed three times with TBS-T, followed by incubation with primary antibody in 5% BSA in TBS-T (other antibodies, o/n at 4°C). Membranes were washed three times with TBS-T, incubated with matching secondary antibody in 5% milk in TBS-T (1:5000, 1 h at rt) and then washed three times with TBS-T and once with TBS. Luminol development solution (10 mL of 1.4 mM luminol in 100 mM Tris pH 8.8 + 100 µL of 6.7 mM *p*-coumaric acid in DMSO + 3 µL of 30% (v/v) $H_2O_2$) or Clarity Max™ ECL Substrate (Bio-Rad) was added and chemiluminescence was detected on ChemiDoc™ MP System.

Following detection of phospho-proteins, membranes were stripped (Restore™ Plus Stripping Buffer, Thermo Fisher) for 20 min, washed three times with TBS, and blocked and incubated with the control antibodies as described above.

Primary antibodies: monoclonal mouse anti-FLAG M2 (1:5000, Sigma Aldrich, F3156), monoclonal anti-V5 (1:5000, Thermo Fisher, R960-25), monoclonal mouse anti-β-actin (1:1000, Abcam, ab8227), polyclonal rabbit anti-Lamin B1 (1:5000, Thermo Fisher, PA5-19468), polyclonal rabbit anti-phospho-FES Y713 (1:1000, Thermo Fisher, PA5-64504), monoclonal rabbit anti-FES (1:1000, Cell Signaling Technology (CST), #85704), polyclonal rabbit anti-phospho-SYK Y352 (1:1000, CST, #2701), monoclonal rabbit anti-SYK (1:1000, CST, #13198), polyclonal rabbit anti-phospho-HS1 Y397 (1:1000, CST, #4507), polyclonal rabbit anti-HS1 (1:1000, CST, #4503), polyclonal rabbit anti-phospho-PLCγ2 Y1217 (1:1000, CST, #3871), polyclonal rabbit anti-PLCγ2 (1:1000, CST, #3872). Secondary antibodies: goat anti-mouse-HRP (1:5000, Santa Cruz, sc-2005), goat anti-rabbit-HRP (1:5000, Santa Cruz, sc-2030).

#### 16. CD11b expression analysis by flow cytometry

Cells ($1x10^6$ per sample) were centrifuged (500 *g*, 3 min) and suspended in human FcR blocking solution (Miltenyi Biotec, 25x diluted in FACS buffer (1% BSA, 1% FCS, 0.1% $NaN_3$, 2 mM EDTA in PBS)), transferred to a V-bottom 96-well plate and incubated for 10 min at 4°C. Next, monoclonal rat CD11b-APC antibody (1:100, Miltenyi Biotec, 130-113-231) or rat anti-IgG2b-APC isotype control antibody (1:100, Miltenyi Biotec, 130-106-728) was added along with 7-AAD (1 µg/mL) and samples were incubated for 30 min 4°C in the dark. Samples were washed once in PBS and fixed in 1% PFA in PBS for 15 min at 4°C in the dark, followed by two washing steps in PBS and resuspension in FACS buffer to a density of approximately 500 cells/µL. Cell suspensions were measured on a Guava easyCyte HT and data was processed using GuavaSoft InCyte 3.3 (Merck Millipore). Events (generally 20,000 per condition) were gated by forward and side scatter (cells), side scatter area (singlets) and viability (live cells) and the percentage of CD11b-positive cells was determined based on background fluorescence for isotype control antibody and non-differentiated cells. The RED-R channel (661/15 filter) and RED-B channel (695/50 filter) were used to detect CD11b-APC and 7-AAD, respectively.

#### 17. Neutrophil oxidative burst assay

Cells were centrifuged (500 *g*, 3 min), suspended in PBS and seeded in triplicate per condition in 96-well plates (100,000 cells per well in 100 µL). To each well, 100 µL of PBS containing nitroblue tetrazolium (NBT) with vehicle or PMA was added, bringing final concentrations to 0.1% and 1.6 µM, respectively. Cells were incubated (1 h, 37°C) and plates were imaged by phase contrast microscopy (20x magnification, EVOS FL Auto 2). Cells positive for formazan deposits were counted (3 different fields per replicate).

### 18. TempO-Seq transcriptome profiling

HL-60 cells were seeded at 20,000 cells per well in a 96-well plate and differentiated towards macrophages as aforementioned with three independent biological replicates. After 48 h, medium was removed, cells were washed with 100 µL PBS and subsequently lysed in 50 µL 1x BioSpyder lysis buffer (15 min, rt). Lysates were flash-frozen in liquid nitrogen, stored at -80 °C and shipped to BioSpyder technologies on dry ice, where the TempO-Seq® profiling was conducted as previously described.[15] Raw gene transcription counts were subjected to internal normalization using DESeq2[16] and statistical significance determined by *t*-test with Benjamini-Hochberg multiple comparison correction (FDR = 1%). Differentially expressed genes were identified using following cut-offs: fold change of corrected counts of $FES^{S700C}$ over WT HL-60 cells < 0.5 or > 2 with q-value < 0.05.

### 19. Co-immunoprecipitation

U2OS cells were co-transfected with FLAG-tagged FES and V5-tagged SYK and grown for 48 h as described above, followed by incubation with vehicle or WEL028 (200 nM) for 1 h. Cells were then washed with PBS and collected by scraping in IP-buffer (20 mM Tris-HCl pH 7.5, 150 mM NaCl, 1% Triton X-100 supplemented with 1x Halt™ phosphatase and protease inhibitor cocktail (Thermo Fisher)). Cells were lysed by sonication on ice (3 cycles of 10'' on, 10'' off at 25% maximum amplitude), centrifuged (14,000 *g*, 10 min, 4°C). The clear lysate was diluted to 1 mg/mL in IP-buffer and subjected to immunoprecipitation using Dynabeads™ Protein G Immunoprecipitation kit (Thermo Fisher) following manufacturer's protocol. Briefly, anti-FLAG M2 antibody (1:100, Sigma Aldrich, F3156) was incubated with beads with gentle rotation (10 min, rt), after which lysate (500 µL of 1 mg/mL) was added and incubated (1 h, 4°C). Beads were washed three times, transferred to clean tubes and eluted by suspension in 2x Laemmli buffer (50 µL, 10 min, 70°C). Samples (10 µL per lane) were resolved by SDS-PAGE and immunoblotted using anti-V5 or anti-FLAG antibodies. Immunoprecipitations were performed in three independent replicates.

### 20. Phagocytosis assays

*Flow cytometry*

HL-60 cells were differentiated to neutrophils as described. Cells were counted and centrifuged (200 *g*, 5 min), followed by resuspension in growth medium without antibiotics. Cells were incubated with vehicle or inhibitor (from 1000x concentrated stocks in DMSO) for 1 h at 37°C prior to infection ($1x10^6$ cells in 900 µL in 12-well plate). Meanwhile, *E. coli* B834(DE3) constitutively expressing $GFP^{A206K}$ were grown in LB medium to an OD600 of 0.4-1.0, after which bacteria were centrifuged (2,000 *g*, 5 min), washed and resuspended in PBS to appropriate density.[17] Neutrophils were infected by addition of bacteria at multiplicity of infection (MOI) of 30 unless stated otherwise. Cells were then incubated for 1 h at 37°C unless stated otherwise, after which cells were resuspended and transferred to Eppendorf tubes, washed in FACS buffer (1 mL, 500 *g*, 3 min) and fixed in 1% PFA in PBS (15 min, 4°C, in the dark). Samples were further processed as described above.

Events (generally 20,000 per condition) were gated by forward and side scatter (cells), side scatter area (singlets) and the percentage of GFP-positive cells and GFP mean fluorescence intensity (MFI) were determined based on background fluorescence for non-infected cells. The GREEN-B channel (525/30 filter) was used to detect GFP. Phagocytic index was calculated as fraction of GFP-positive cells (number of phagocytic cells) multiplied by GFP MFI (number of phagocytized bacteria).

*Detection of phosphoproteins by immunoblot*

HL-60 neutrophils were infected as described above, but in 15 mL tubes ($5x10^6$ cells in 4 mL per sample). After indicated infection times, ice-cold PBS was added (10 mL) and suspensions were immediately centrifuged (3,500 *g*, 2 min). Supernatant was completely aspirated and cells were thoroughly resuspended in 2x Laemmli sample buffer (100 µL), followed by incubation at 95°C for 10 min and brief sonication to reduce sample viscosity. Samples were stored at -20°C or immediately resolved by SDS-PAGE (20 µL per lane).

**21. Statistical analysis**

All statistical measures and methods are included in the respective Figure or Table captions. In brief: all replicates represent biological replicates and all data represent means ± SEM, unless indicated otherwise. Statistical significance was determined using Student's *t*-tests (two-tailed, unpaired) or ANOVA with Holm-Sidak's multiple comparisons correction. *** $P < 0.001$; ** $P < 0.01$; * $P < 0.05$; NS if $P > 0.05$. All statistical analyses were conducted using GraphPad Prism® 7 or Microsoft Excel.

**Supplementary Materials & Methods - Chemistry**

*General information*

All reactions were performed using oven- or flame-dried glassware and dry solvents. Reagents were purchased from Sigma-Aldrich, Acros, and Merck and used without further purification unless noted otherwise. All moisture sensitive reactions were performed under an argon atmosphere. $^{1}$H and $^{13}$C NMR spectra were recorded on a Bruker AV 400 MHz spectrometer at 400.2 ($^{1}$H) and 100.6 ($^{13}$C) MHz or on a Bruker DMX-600 spectrometer at 600 ($^{1}$H) and 151 ($^{13}$C) MHz using $CDCl_3$, DMSO-*d6* or MeOD as solvent. Chemical shift values are reported in ppm with tetramethylsilane or solvent resonance as the internal standard ($CDCl_3$: δ 7.26 for $^{1}$H, δ 77.16 for $^{13}$C; DMSO-*d6*, δ 2.50 for $^{1}$H, δ 39.52 for $^{13}$C; MeOD: δ 3.31 for $^{1}$H, δ 49.00 for $^{13}$C). Data are reported as follows: chemical shifts (δ), multiplicity (s = singlet, d = doublet, dd = double doublet, td = triple doublet, t = triplet, q = quartet, br = broad, m = multiplet), coupling constants J (Hz), and integration. HPLC purification was performed on a preparative LC-MS system (Agilent 1200 serie) with an Agilent 6130 Quadruple MS detector. High-resolution mass spectra were recorded on a Thermo Scientific LTQ Orbitrap XL. Compound purity (>95% unless stated otherwise) was determined by liquid chromatography on a Finnigan Surveyor LC-MS system, equipped with a C18 column. Flash chromatography was performed using SiliCycle silica gel type SiliaFlash P60 (230−400 mesh). TLC analysis was performed on Merck silica gel 60/Kieselguhr F254, 0.25 mm. Compounds were visualized using $KMnO_4$ stain ($K_2CO_3$ (40 g), $KMnO_4$ (6 g) in water (600 mL)) or ninhydrin stain (ninhydrin (20 g) in ethanol (600 mL)).

*Scheme 1 - Synthesis of compound **2***

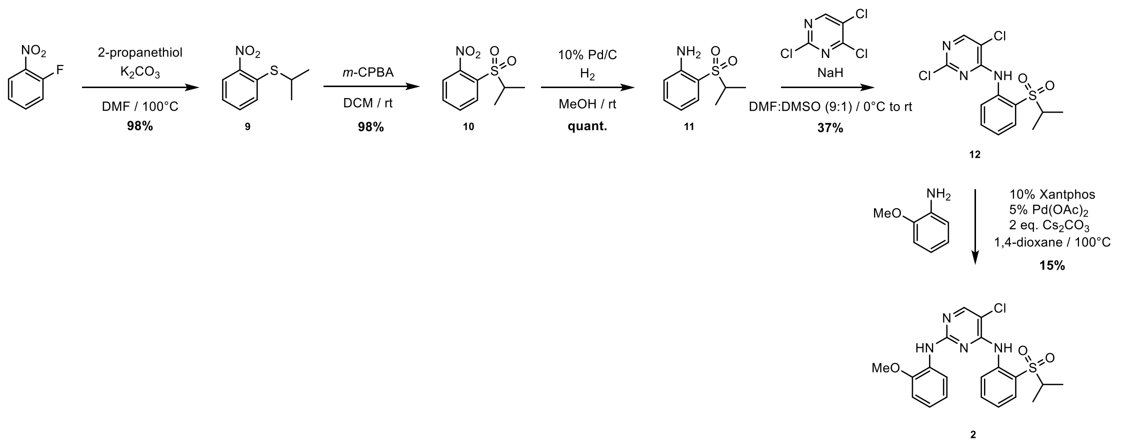


*Synthesis of isopropyl(2-nitrophenyl)-sulfane (**9**)*

1-Fluoro-2-nitrobenzene (9.88 g, 70 mmol) was dissolved in dry DMF (100 mL), followed by the addition of 2-propanethiol (5.47 g, 70 mmol, 1 eq.) and potassium carbonate (24.19 g, 175 mmol, 2.5 eq.). The reaction mixture was heated at 100°C for 16 h and subsequently cooled to rt. The mixture was diluted with $H_2O$ (200 mL) and extracted with EtOAc (3 x 200 mL). The organic layers were combined, dried over $Na_2SO_4$ and concentrated to obtain title compound (13.5 g, 68 mmol, 98%). $^1$H NMR (400 MHz, $CDCl_3$): δ 8.12 (dd, *J* = 8.3, 1.5 Hz, 1H), 7.54 (ddd, *J* = 8.4, 7.1, 1.4 Hz, 1H), 7.48 (dd, *J* = 8.1, 1.4 Hz, 1H), 7.25 (ddd, *J* = 8.4, 7.1, 1.5 Hz, 1H), 3.58 (p, *J* = 6.6 Hz, 1H), 1.40 (d, *J* = 6.6 Hz, 6H). $^{13}$C NMR (101 MHz, $CDCl_3$): δ 147.2, 136.5, 133.3, 128.3, 125.9, 124.9, 35.8, 22.4.

*Synthesis of 1-(isopropylsulfonyl)-2-nitrobenzene (**10**)*

Compound **9** (12.2 g, 61.3 mmol) was dissolved in DCM (120 mL), followed by portion wise addition of 3-chloroperbenzoic acid (31.6 g, 143 mmol, 2.3 eq.) The reaction mixture was stirred at rt for 16 h. The mixture was diluted with 10% $Na_2SO_3$ (120 mL) and stirred for 10 min, after which layers were separated and the aqueous layer was extracted with DCM (3 x 120 mL). The organic layers were combined, washed with sat. $NaHCO_3$ (2 x 200 mL), brine (1 x 250 mL), dried over $MgSO_4$ and concentrated to obtain title compound (13.9 g, 60.7 mmol, 98%). $^1$H NMR (400 MHz, $CDCl_3$): δ 8.15 – 8.07 (m, 1H), 7.86 – 7.72 (m, 3H), 4.04-3.95 (m, 1H), 1.41 (d, *J* = 6.8 Hz, 6H). $^{13}$C NMR (101 MHz, $CDCl_3$): δ 134.8, 133.1, 132.2, 125.1, 56.0, 15.5.

*Synthesis of 2-(isopropylsulfonyl)aniline (**11**)*

Compound **10** (2.03 g, 8.85 mmol) was dissolved in MeOH (25 mL), followed by addition of Pd/C (93 mg, 0.87 mmol, 0.1 eq.). The mixture was stirred under $H_2$ atmosphere for 16 h at rt, after which it was filtered over celite and concentrated to obtain title compound (1.76 g, 8.85 mmol, quant.). $^1$H NMR (400 MHz, $CDCl_3$): δ 7.64 (dd, *J* = 8.0, 1.5 Hz, 1H), 7.40 – 7.30 (m, 1H), 6.80 (m, 1H), 6.74 (d, *J* = 8.2 Hz, 1H), 4.49 (s, 2H), 3.34 (m, 1H), 1.31 (d, *J* = 6.7 Hz, 6H). $^{13}$C NMR (101 MHz, $CDCl_3$): δ 147.2, 135.2, 131.4, 118.3, 117.7, 117.7, 54.3, 15.4.

*Synthesis of 2,5-dichloro-N-(2-(isopropylsulfonyl)phenyl)pyrimidin-4-amine (**12**)*

A mixture of DMF/DMSO (10:1 ratio, 10 mL) was cooled to 0°C, after which NaH (354 mg, 14.8 mmol, 2.5 eq.) was added. Compound **11** (1.15 g, 5.77 mmol) was dissolved in DMF/DMSO (10:1 ratio, 5 mL) and added to the reaction mixture. The suspension was stirred at 0°C for 30 min, followed by the addition of 2,4,5-trichloropyrimidine (1.94 g, 10.6 mmol) diluted in DMF/DMSO (5 mL). The mixture was allowed to warm to rt and was then stirred for 16 h. The reaction mixture was diluted with $H_2O$ (150 mL) and extracted with EtOAc (50 mL). The organic layer was washed with $H_2O$ (5 x 50 mL), 5% LiCl (50 mL) and brine (50 mL), dried over $MgSO_4$ and subsequently concentrated. The crude residue was purified by flash column chromatography (pentane → 20% EtOAc in pentane), yielding title compound (777 mg, 2.24 mmol, 37%). $^1$H NMR (400 MHz, $CDCl_3$): δ 10.06 (s, 1H), 8.62 (dd, *J* = 8.5, 1.2 Hz, 1H), 8.30 (s, 1H), 7.92 (dd, *J* = 8.0, 1.7 Hz, 1H), 7.75-7.71 (m, 1H), 7.37-7.27 (m, 1H), 3.26-3.16 (m, 1H), 1.31 (d, *J* = 6.8 Hz, 6H). $^{13}$C NMR (101 MHz, $CDCl_3$): δ 157.9, 156.4, 155.7, 137.5, 135.3, 131.6, 124.7, 124.3, 122.8, 115.4, 56.2, 15.5.

*Synthesis of 5-chloro-$N^4$-(2-(isopropylsulfonyl)phenyl)-$N^2$-(2-methoxyphenyl)pyrimidine-2,4-diamine (**2**)*

Compound **12** (399 mg, 1.15 mmol), *o*-anisidine (142 mg, 1.15 mmol), XantPhos (67 mg, 0.12 mmol), $Pd(OAc)_2$ (14 mg, 0.06 mmol) and $Cs_2CO_3$ (756 mg, 2.30 mmol) were dissolved in dry 1,4-dioxane (15 mL) under argon and the resulting mixture was heated at 100°C for 16 h. The reaction mixture was diluted with EtOAc (50 mL), filtered over celite and subsequently concentrated. The crude residue was purified by flash column chromatography (pentane → 30% EtOAc in pentane), yielding title compound (77 mg, 0.18 mmol, 15%). HRMS (ESI+) m/z: calculated for $C_{20}H_{21}ClN_4O_3S$ ([M+H]): 433.10957; found: 433.10837. $^1$H NMR (400 MHz, DMSO): δ 9.53 (s, 1H), 8.53 (d, *J* = 8.4 Hz, 1H), 8.37 (s, 1H), 8.25 (s, 1H), 7.82 (dd, *J* = 8.0, 1.6 Hz, 1H), 7.72 (dd, *J* = 7.9, 1.5 Hz, 1H), 7.66-7.62 (m, 1H), 7.38-7.29 (m, 1H), 7.14-7.02 (m, 2H), 6.92-6.88 (m, 1H), 3.79 (s, 3H), 3.47-3.40 (m, 1H), 1.15 (d, *J* = 6.8 Hz, 6H). $^{13}$C NMR (101 MHz, DMSO): δ 158.6, 155.8, 155.3, 151.3, 138.5, 135.3, 131.4, 128.5, 124.6, 124.5, 124.0, 123.9, 123.5, 120.6, 111.6, 105.1, 56.0, 55.3, 15.3.

*Scheme 2 – Synthesis of compound **3***

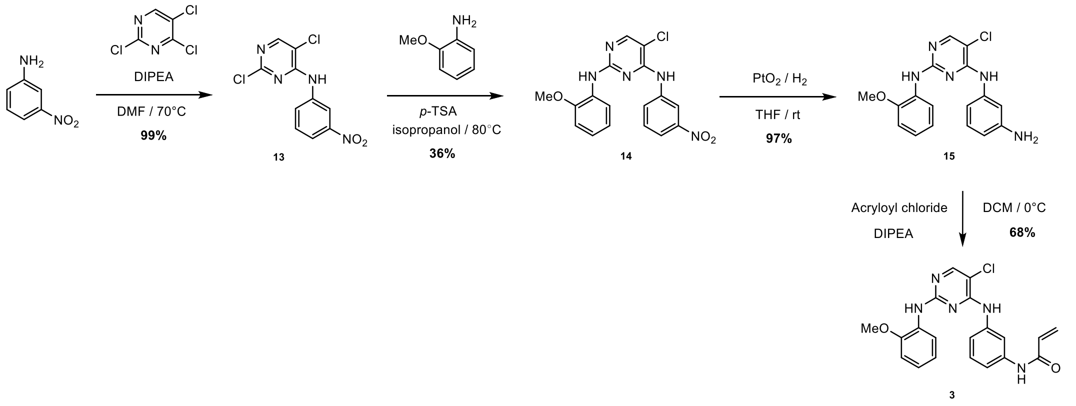


*Synthesis of 2,5-dichloro-N-(3-nitrophenyl)pyrimidin-4-amine (**13**)*

2,4-5-Trichloropyrimidine (1.42 g, 7.74 mmol) and DIPEA (2.00 g, 15.5 mmol) were dissolved in DMF (4 mL). To the stirring solution was added 3-nitroaniline (1.07 g, 7.74 mmol), after which the reaction mixture was heated at reflux for 16 h. The mixture was allowed to cool down to rt and subsequently diluted with EtOAc (75 mL) and washed with $H_2O$ (3 x 50 mL). The organic layer was dried over $MgSO_4$ and concentrated to dryness, yielding title compound (2.18 g, 7.66 mmol, 99%). $^1$H NMR (400 MHz, DMSO): δ 9.96 (s, 1H), 8.63 (t, $J$ = 2.2 Hz, 1H), 8.49 (s, 1H), 8.17 – 8.09 (m, 1H), 8.01 (ddd, $J$ = 8.3, 2.3, 0.9 Hz, 1H), 7.68 (t, $J$ = 8.2 Hz, 1H). $^{13}$C NMR (101 MHz, DMSO): δ 157.0, 156.2, 147. 8, 139.0, 129.9, 129.0, 122.5, 119.1, 117.2, 114.2.

*Synthesis of 5-chloro-N$^2$-(2-methoxyphenyl)-N$^4$-(3-nitrophenyl)pyrimidine-2,4-diamine (**14**)*

Compound **13** (600 mg, 2.10 mmol) and *o*-anisidine (259 mg, 2.10 mmol) were taken up in isopropanol (20 mL), followed by the addition of *p*-TSA (400 mg, 2.10 mmol). The reaction mixture was heated under reflux for 16 h, after which it was concentrated under reduced pressure. The residue was taken up in saturated aqueous $NaHCO_3$ (20 mL) and the product was extracted with EtOAc (20 mL). The organic layer was concentrated under reduced pressure and the crude residue was purified by flash column chromatography (10% → 30% EtOAc in pentane), yielding title compound (284 mg, 0.38 mmol, 36%). $^1$H NMR (400 MHz, DMSO): δ 9.31 (s, 1H), 8.51 (t, $J$ = 2.2 Hz, 1H), 8.23 – 8.15 (m, 2H), 8.09 (s, 1H), 7.92 (ddd, $J$ = 8.2, 2.3, 0.9 Hz, 1H), 7.81 – 7.74 (m, 1H), 7.55 (t, $J$ = 8.2 Hz, 1H), 7.04 – 6.98 (m, 2H), 6.80 – 6.74 (m, 1H), 3.80 (s, 3H). $^{13}$C NMR (101 MHz, DMSO): δ 157.8, 155.6, 155.5, 150.0, 147.7, 140.0, 129.5, 128.5, 128.1, 123.5, 121.8, 120.0, 117.8, 116.8, 110.9, 104.3, 55.6.

*Synthesis of N$^4$-(3-aminophenyl)-5-chloro-N$^2$-(2-methoxyphenyl)pyrimidine-2,4-diamine (**15**)*

Compound **14** (200 mg, 0.54 mmol) was dissolved in anhydrous THF (10 mL), followed by addition of $PtO_2$ (21 mg, 0.09 mmol, 0.1 eq.). The mixture was stirred under $H_2$ atmosphere for 72 h at rt, after which it was diluted in MeOH, filtered over celite and concentrated to obtain title compound (179 mg, 0.54 mmol, 97%), which was directly used in the next step.

*Synthesis of N-(3-((5-chloro-2-((2-methoxyphenyl)amino)pyrimidin-4-yl)amino)phenyl)acrylamide (**3**)*

Compound **15** (77 mg, 0.23 mmol) was taken up in DCM (2 mL) and cooled to 0°C. Subsequently, DIPEA (29 mg, 0.23 mmol) was added and the reaction mixture was stirred for 10 min. Acryloyl chloride (20 mg, 0.23 mmol) dissolved in DCM (1 mL) was dropwisely added to the mixture. After stirring for 30 min, the reaction was quenched by addition of water (5 mL). The product was extracted from the reaction mixture with DCM (3 x 25 mL), the organic extract was dried over $MgSO_4$ and concentrated to dryness. The crude residue was purified by flash column chromatography (10% → 40% EtOAc in pentane), yielding title compound (61 mg, 0.15 mmol, 68%). HRMS (ESI+) m/z: calculated for $C_{20}H_{18}ClN_5O_2$ ([M+H]): 396.12218; found: 396.12156. $^{1}$H NMR (400 MHz, $CDCl_3$): δ 8.21 (dd, *J* = 7.9, 1.6 Hz, 1H), 8.04 (d, *J* = 12.2 Hz, 2H), 7.70 (s, 1H), 7.58 (d, *J* = 7.9 Hz, 1H), 7.41 (s, 1H), 7.33 (t, *J* = 8.0 Hz, 1H), 7.26 (d, *J* = 8.5 Hz, 1H), 7.16 (s, 1H), 7.04 – 6.95 (m, 1H), 6.94 – 6.83 (m, 2H), 6.45 (dd, *J* = 16.9, 1.3 Hz, 1H), 6.23 (dd, *J* = 16.8, 10.2 Hz, 1H), 5.79 (dd, *J* = 10.2, 1.3 Hz, 1H), 3.88 (s, 3H). $^{13}$C NMR (101 MHz, $CDCl_3$): δ 155.7, 148.8, 138.6, 138.5, 131.2, 129.6, 128.8, 128.2, 122.6, 120.5, 120.1, 117.3, 116.5, 115.7, 112.8, 110.3, 105.1, 55.8.

Scheme 3 – Synthesis of compound **4**

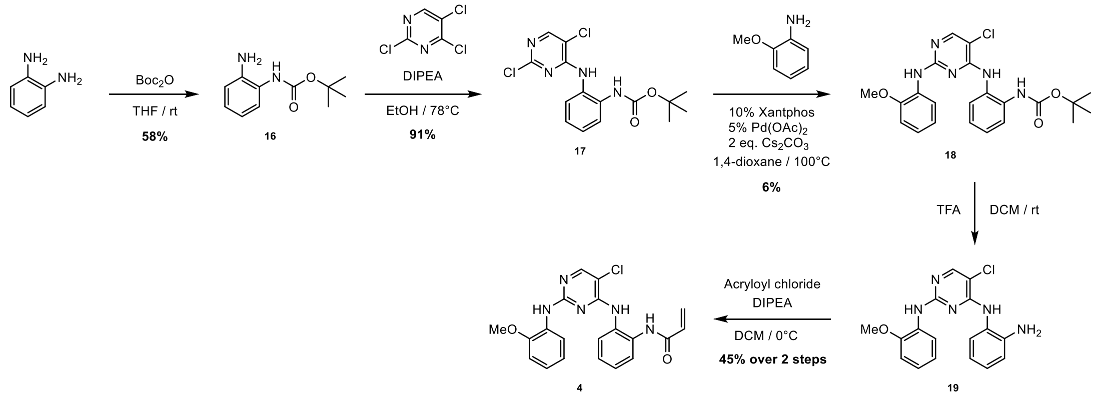

*Synthesis of tert-butyl (2-aminophenyl)carbamate (**16**)*

To a solution of *o*-phenylenediamine (2.19 g, 20.2 mmol) in THF (20 mL) was dropwisely added a solution of $(Boc)_2O$ (4.45 g, 20.4 mmol) in THF (5 mL). The reaction mixture was stirred at rt for 16 h, after which the mixture was concentrated under reduced pressure and the residue was taken up in a cold mixture of EtOAc/Petroleum ether (1:4 ratio, 15 mL), causing the product to precipitate. The precipitate was collected by filtration and dried to yield the title compound (2.43 g, 11.7 mmol, 58%). $^1H$ NMR (400 MHz, $CDCl_3$): δ 7.28 (s, 1H), 7.00 (td, *J* = 7.6, 1.5 Hz, 1H), 6.82 – 6.74 (m, 2H), 6.28 (s, 1H), 3.55 (s, 2H), 1.51 (s, 9H). $^{13}C$ NMR (101 MHz, $CDCl_3$): δ 154.0, 140.0, 126.3, 124.9, 120.5, 119.7, 117.7, 117.0, 80.6, 28.5, 28.4. Spectroscopic data are in accordance with those reported in literature[18].

*Synthesis of tert-butyl (2-((2,5-dichloropyrimidin-4-yl)amino)phenyl)carbamate (**17**)*

2,4-5-Trichloropyrimidine (1.18 g, 6.43 mmol) and DIPEA (1.68 g, 13.0 mmol) were dissolved in EtOH (25 mL). To the stirring solution was added compound **16** (1.35 g, 6.48 mmol), after which the reaction mixture was heated at reflux for 16 h. After TLC indicated depletion of starting material, the mixture was allowed to cool down to rt and subsequently triturated with cold $H_2O$ (20 mL), causing precipitation of the product. The precipitate was collected by vacuum filtration and dried to yield the title compound (2.11 g, 5.94 mmol, 92%). $^1H$ NMR (400 MHz, $CDCl_3$): δ 8.64 (s, 1H), 8.16 (s, 1H), 7.78 (dd, *J* = 8.1, 1.2 Hz, 1H), 7.32-7.27 (m, 1H), 7.24 – 7.16 (m, 2H), 6.59 (s, 1H), 1.53 (s, 9H). $^{13}C$ NMR (101 MHz, $CDCl_3$): δ 158.3, 157.4, 154.8, 154.7, 130.9, 130.2, 126.6, 126.5, 126.4, 124.9, 114.4, 82.0, 28.4.

*Synthesis of tert-butyl(2-((5-chloro-2-((2-methoxyphenyl)amino)pyrimidin-4-yl)amino)phenyl)carbamate (**18**)*

Compound **17** (600 mg, 1.69 mmol), *o*-anisidine (208 mg, 1.69 mmol), Xantphos (98 mg, 0.17 mmol), $Pd(OAc)_2$ (19 mg, 0.084 mmol) and cesium carbonate (1.10 g, 3.38 mmol) were dissolved in 20 mL dry 1,4-dioxane. The reaction mixture was purged with argon and subsequently heated at 100°C for 16 h. Subsequently, the mixture was diluted with EtOAc and filtered over Celite®, after which the filtrate was concentrated. The crude residue was purified by flash column chromatography (pentane → 20% EtOAc in pentane), yielding title compound (48 mg, 0.11 mmol, 6%). $^1H$ NMR (400 MHz, $CDCl_3$): δ 8.12 – 8.03 (m, 2H), 7.79 (s, 1H), 7.72 – 7.66 (m, 1H), 7.63 (s, 1H), 7.49 – 7.41 (m, 1H), 7.29 – 7.22 (m, 2H),

6.89 (m, 1H), 6.82 (dd, $J$ = 8.1, 1.5 Hz, 1H), 6.78 – 6.70 (m, 1H), 6.66 (s, 1H), 3.85 (s, 3H), 1.50 (s, 9H). $^{13}C$ NMR (101 MHz, $CDCl_3$): δ 157.0, 154.1, 147.8, 131.9, 130.9, 129.2, 128.9, 127.6, 126.5, 125.4, 123.9, 121.6, 120.8, 118.7, 114.2, 109.8, 105.2, 81.5, 55.8, 28.4.

*Synthesis of $N^4$-(2-aminophenyl)-5-chloro-$N^2$-(2-methoxyphenyl)pyrimidine-2,4-diamine (**19**)*

Compound **18** (50 mg, 0.11 mmol) was dissolved in DCM (3 mL), after which an equal volume of TFA was slowly added. The reaction mixture was stirred for 16 h at rt and subsequently evaporated to dryness to afford the title compound, which was directly used in the next reaction.

*Synthesis of N-(2-((5-chloro-2-((2-methoxyphenyl)amino)pyrimidin-4-yl)amino)phenyl)acrylamide (**4**)*

Compound **19** (84 mg, 0.18 mmol) was taken up in DCM (2 mL) and cooled to 0°C. Subsequently, DIPEA (24 mg, 0.18 mmol) was added and the reaction mixture was stirred for 10 min. Acryloyl chloride (17 mg, 0.18 mmol) dissolved in DCM (1 mL) was dropwisely added to the mixture. After stirring for 15 min, the reaction was quenched by addition of water (5 mL). The product was extracted from the reaction mixture with DCM (50 mL), the organic extract was dried over $MgSO_4$ and concentrated to dryness. The crude residue was purified by flash column chromatography (10% → 40% EtOAc in pentane), yielding title compound (20 mg, 0.05 mmol, 45% over two steps). HRMS (ESI+) m/z: calculated for $C_{20}H_{18}ClN_5O_2$ ([M+H]): 396.12218; found: 396.12156. $^1H$ NMR (400 MHz, MeOD): δ 8.04 (s, 1H), 7.81 (dd, $J$ = 8.1, 1.6 Hz, 1H), 7.76 – 7.69 (m, 1H), 7.51 – 7.44 (m, 1H), 7.41 – 7.33 (m, 2H), 7.03 – 6.95 (m, 2H), 6.73 (ddd, $J$ = 8.6, 7.1, 1.8 Hz, 1H), 6.48 (dd, $J$ = 17.0, 9.5 Hz, 1H), 6.40 (dd, $J$ = 16.9, 2.4 Hz, 1H), 5.81 (dd, $J$ = 9.4, 2.4 Hz, 1H), 3.87 (s, 3H). $^{13}C$ NMR (101 MHz, MeOD): δ 166.9, 158.8, 149.9, 132.9, 132.5, 131.4, 129.9, 129.2, 128.9, 128.2, 127.9, 127.4, 126.0, 125.5, 125.0, 122.0, 121.6, 111.5, 106.5, 56.3.

*Scheme 4 – Synthesis of compound **5***

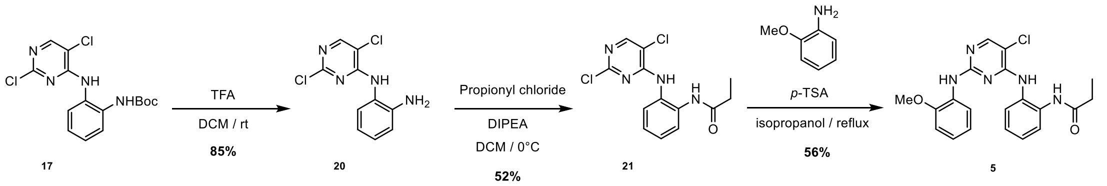

*Synthesis of N[1]-(2,5-dichloropyrimidin-4-yl)benzene-1,2-diamine (**20**)*

Compound **17** (323 mg, 0.91 mmol) was dissolved in DCM (10 mL), after which TFA (3 mL) was slowly added. The rection mixture was stirred at rt for 16 h and subsequently evaporated to dryness. The residue was dissolved in water, neutralized with aqueous $K_2CO_3$ to pH 7-8 and the resulting precipitate was collected by filtration and dried to afford title compound (197 mg, 0.77 mmol, 85%). $^1$H NMR (400 MHz, DMSO): δ 9.01 (s, 1H), 8.14 (s, 1H), 7.03 (d, *J* = 7.8 Hz, 1H), 6.95 (s, 1H), 6.72 (d, *J* = 8.0 Hz, 1H), 6.54 (t, *J* = 7.5 Hz, 1H), 4.92 (s, 2H). $^{13}$C NMR (101 MHz, DMSO): δ 158.7, 158.2, 157.3, 154.4, 144.9, 128.4, 127.8, 121.7, 115.8, 115.6.

*Synthesis of N-(2-((2,5-dichloropyrimidin-4-yl)amino)phenyl)propionamide (**21**)*

Compound **20** (171 mg, 0.67 mmol) was dissolved in DCM (5 mL) and cooled to 0°C. Subsequently, DIPEA (85 mg, 0.67 mmol) was added and the reaction mixture was stirred for 10 min. Propionyl chloride (64 mg, 0.67 mmol) dissolved in DCM (3 mL) was dropwisely added. After stirring for 2 h, the reaction mixture was diluted with DCM (50 mL) and washed with $H_2O$ (2 x 50 mL). The organic extracted extract was dried over $MgSO_4$ and concentrated to dryness. The crude residue was purified by flash column chromatography (20% → 30% EtOAc in pentane), yielding title compound (109 mg, 0.35 mmol, 52%). $^1$H NMR (400 MHz, $CDCl_3$): δ 8.67 (s, 1H), 8.16 (s, 1H), 8.07 (s, 1H), 7.73 (dd, *J* = 8.2, 1.3 Hz, 1H), 7.32 – 7.23 (m, 1H), 7.14 – 7.05 (m, 1H), 6.87 (dd, *J* = 7.9, 1.5 Hz, 1H), 2.38 (q, *J* = 7.6 Hz, 2H), 1.20 (t, *J* = 7.6 Hz, 3H). $^{13}$C NMR (101 MHz, $CDCl_3$): δ 174.4, 158.1, 157.4, 155.0, 131.4, 130.2, 127.0, 126.5, 126.1, 125.0, 114.7, 30.0, 10.2.

*Synthesis of N-(2-((5-chloro-2-((2-methoxyphenyl)amino)pyrimidin-4-yl)amino)phenyl)propionamide (**5**)*

Compound **21** (59 mg, 0.19 mmol) and *o*-anisidine (23 mg, 0.19 mmol) were taken up in isopropanol (10 mL), followed by the addition of *p*-TSA (37 mg, 0.19 mmol). The reaction mixture was heated under reflux for 16 h, after which it was concentrated under reduced pressure. The residue was taken up in saturated aqueous $NaHCO_3$ (20 mL) and the product was extracted with EtOAc (20 mL). The organic layer was dried over $MgSO_4$ and concentrated. The crude residue was purified by flash column chromatography (10% → 40% EtOAc in pentane), yielding a mixture of compound **21** and **5**, which was further purified by HPLC, affording title compound (42 mg, 0.10 mmol, 56%). HRMS (ESI+) m/z: calculated for $C_{20}H_{20}ClN_5O_2$ ([M+H]): 398.13783; found: 398.13722. $^1$H NMR (400 MHz, $CDCl_3$): δ 10.04 (s, 1H), 9.24 (s, 1H), 7.85 (s, 1H), 7.68 – 7.60 (m, 2H), 7.58 (dd, *J* = 8.0, 1.6 Hz, 1H), 7.26 (s, 3H), 7.16 (dd, *J* = 7.5, 2.0 Hz, 1H), 7.06 (ddd, *J* = 8.3, 7.6, 1.6 Hz, 1H), 6.84 (dd, *J* = 8.3, 1.4 Hz, 1H), 6.70 (td, *J* = 7.8, 1.3 Hz, 1H), 3.80 (s, 3H), 2.46 (q, *J* = 7.5 Hz, 2H), 1.26 (t, *J* = 7.6 Hz, 3H). $^{13}$C NMR (101 MHz,

$CDCl_3$): δ 174.3, 157.7, 153.3, 152.8, 151.2, 130.6, 127.8, 127.6, 126.9, 126.1, 125.8, 124.9, 123.1, 120.3, 111.0, 105.4, 56.0, 30.3, 10.2.

*Scheme 5 – Synthesis of **6** (**WEL028**)*

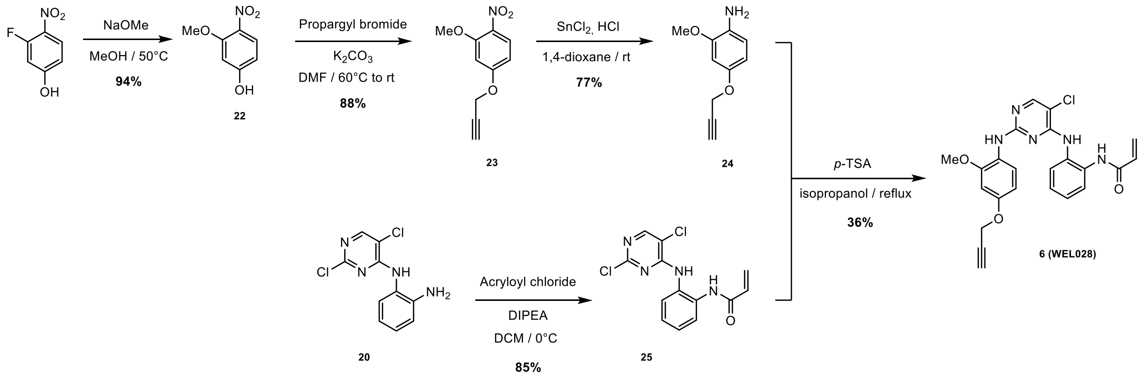


*Synthesis of 3-methoxy-4-nitrophenol (**22**)*

3-Fluoro-4-nitrophenol (1.00 g, 6.4 mmol) was added to a solution of NaOMe in MeOH (0.5M, 14 mL), which was then heated at 50°C for 12 h. Additional NaOMe in MeOH (0.5M, 14 mL) was added and the mixture was stirred at 50°C until TLC indicated complete depletion of starting material. The mixture was diluted with $H_2O$ (100 mL), neutralized with 3M HCl and then extracted with EtOAc (3 x 100 mL). The combined organic layers were washed with brine (100 mL), dried over $MgSO_4$ and evaporated to yield title compound (1.01 g, 6.0 mmol, 94%). $^1$H NMR (400 MHz, DMSO): δ 10.90 (s, 1H), 7.89 (d, *J* = 9.0 Hz, 1H), 6.60 (d, *J* = 2.4 Hz, 1H), 6.47 (dd, *J* = 9.0, 2.4 Hz, 1H), 3.86 (s, 3H). $^{13}$C NMR (101 MHz, DMSO): δ 164.4, 156.1, 131.2, 128.8, 108.0, 100.8, 56.8. Spectroscopic data are in accordance with those reported in literature[19].

*Synthesis of 2-methoxy-1-nitro-4-(prop-2-yn-1-yloxy)benzene (**23**)*

Compound **22** (600 mg, 3.6 mmol) and $K_2CO_3$ (1.47 g, 10.6 mmol) were taken up in anhydrous DMF (10 mL) and heated at 60°C for 30 min under argon. The reaction mixture was cooled to rt, after which propargyl bromide (1.34 g, 9.0 mmol as 80% (w/w) solution in toluene) was added. The mixture was stirred at rt for 16 h, after which it was poured into ice water (200 mL) with stirring for 10 min. The formed precipitate was collected by filtration and dried under vacuum to yield title compound (648 mg, 3.1 mmol, 88%), which was directly used in the next reaction.

*Synthesis of 2-methoxy-4-(prop-2-yn-1-yloxy)aniline (**24**)*

Compound **23** (600 mg, 2.9 mmol) was dissolved in 1,4-dioxane (10 mL) and cooled to 0°C. Cooled (0 °C) stannous chloride dihydrate (3.28 g, 14.5 mmol) in concentrated HCl (10 mL) was dropwisely added to the reaction mixture. After stirring for 16 h at rt, the mixture was basified to pH > 9 by addition of NaOH pellets and extracted with DCM (4 x 10 mL). The organic layer was washed with brine (1 x 10 mL), dried over $MgSO_4$ and concentrated to dryness. The crude product was then purified by flash column chromatography (pentane → 25% EtOAc in pentane) to afford title compound (395 mg, 2.23 mmol, 77%). $^1$H NMR (400 MHz, $CDCl_3$): δ 6.62 (d, *J* = 8.4 Hz, 1H), 6.52 (d, *J* = 2.6 Hz, 1H), 6.42 (dd, *J* = 8.5, 2.6 Hz, 1H), 4.61 (d, *J* = 2.4 Hz, 2H), 3.81 (s, 3H), 3.49 (s, 2H), 2.51 (t, *J* = 2.4 Hz, 1H). $^{13}$C NMR (101 MHz, $CDCl_3$): δ 151.0, 148.3, 130.8, 115.0, 105.9, 100.7, 79.3, 75.3, 56.8, 55.6.

*Synthesis of N-(2-((2,5-dichloropyrimidin-4-yl)amino)phenyl)acrylamide (**25**)*

Compound **20** (250 mg, 0.98 mmol) was dissolved in DCM (5 mL) and cooled to 0°C. Subsequently, DIPEA (127 mg, 0.98 mmol) was added and the reaction mixture was stirred for 10 min. Acryloyl chloride (93 mg, 1.03 mmol) dissolved in DCM (1 mL) was dropwisely added to the mixture. After stirring for 1 h, the reaction was quenched by addition of water (50 mL). The mixture was extracted with DCM (50 mL), the organic extract was dried over $MgSO_4$ and concentrated. The crude residue was purified by flash column chromatography (20% → 40% EtOAc in pentane), yielding title compound (259 mg, 0.84 mmol, 85%). $^1$H NMR (400 MHz, $CDCl_3$): δ 8.72 (d, *J* = 18.0 Hz, 2H), 8.13 (s, 1H), 7.72 (dd, *J* = 8.2, 1.3 Hz, 1H), 7.26 (td, *J* = 7.8, 1.5 Hz, 1H), 7.08 (td, *J* = 7.7, 1.4 Hz, 1H), 6.95 (dd, *J* = 8.0, 1.5 Hz, 1H), 6.44 (dd, *J* = 16.9, 1.4 Hz, 1H), 6.30 (dd, *J* = 16.9, 10.1 Hz, 1H), 5.77 (dd, *J* = 10.1, 1.6 Hz, 1H). $^{13}$C NMR (101 MHz, $CDCl_3$): δ 165.2, 157.8, 157.2, 154.7, 131.1, 129.8, 129.8, 129.0, 127.0, 126.3, 125.8, 125.0, 114.6.

*Synthesis of N-(2-((5-chloro-2-((2-methoxy-4-(prop-2-yn-1-yloxy)phenyl)amino)pyrimidin-4-yl)amino)-phenyl)acrylamide (**6, WEL028**)*

Compound **24** (26 mg, 0.15 mmol) and compound **25** (46 mg, 0.15 mmol) were taken up in isopropanol (5 mL), followed by the addition of *p*-TSA (28 mg, 0.15 mmol). The reaction mixture was heated under reflux for 16 h, after which it was concentrated under reduced pressure. The residue was taken up in saturated aqueous $NaHCO_3$ (20 mL) and the product was extracted with EtOAc (20 mL). The organic layer was dried over $MgSO_4$ and concentrated to dryness. The crude residue was purified by flash column chromatography (10% → 35% EtOAc in pentane), yielding title compound (24 mg, 0.05 mmol, 36%). HRMS (ESI+) m/z: calculated for $C_{23}H_{20}ClN_5O_3$ ([M+H]): 450.13274; found: 450.13231. $^1$H NMR (400 MHz, $CDCl_3$): δ 8.04 (d, *J* = 17.6 Hz, 2H), 7.87 (d, *J* = 8.9 Hz, 1H), 7.67 (s, 1H), 7.59 (t, *J* = 4.8 Hz, 2H), 7.38 (s, 1H), 7.29 (dd, *J* = 6.0, 3.5 Hz, 2H), 6.52 (d, *J* = 2.7 Hz, 1H), 6.40 – 6.31 (m, 2H), 6.16 (dd, *J* = 16.9, 10.3 Hz, 1H), 5.72 (d, *J* = 10.2 Hz, 1H), 4.65 (d, *J* = 2.4 Hz, 2H), 3.81 (s, 3H), 2.54 (t, *J* = 2.4 Hz, 1H). $^{13}$C NMR (101 MHz, $CDCl_3$): δ 164.6, 157.8, 156.9, 153.5, 149.6, 131.6, 131.1, 130.8, 128.4, 127.2, 126.8, 126.4, 125.1, 123.2, 120.1, 105.1, 99.8, 78.9, 75.7, 56.5, 55.9.

Scheme 6 – Synthesis of compound **7** (**WEL033**)

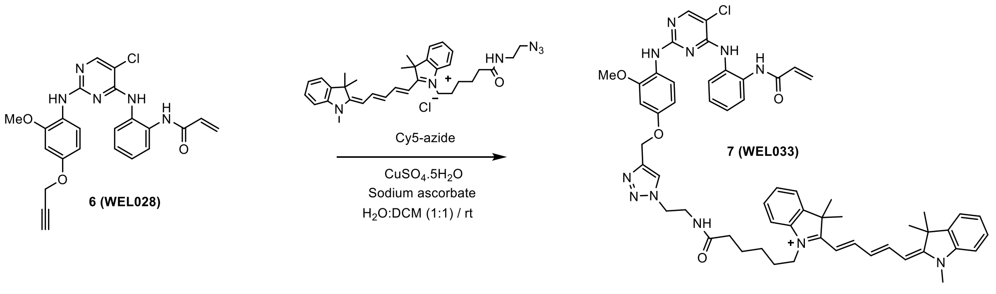


*Synthesis of 1-(6-((2-(4-((4-((4-((2-acrylamidophenyl)amino)-5-chloropyrimidin-2-yl)amino)-3-methoxy-phenoxy)methyl)-1H-1,2,3-triazol-1-yl)ethyl)amino)-6-oxohexyl)-3,3-dimethyl-2-((1E,3E)-5-((E)-1,3,3-trimethylindolin-2-ylidene)penta-1,3-dien-1-yl)-3H-indol-1-ium (**7, WEL033**)*

Compound **6** (30 mg, 0.07 mmol) and Cy5-azide (77 mg, 0.14 mmol) were dissolved in degassed DCM (3 mL). $CuSO_4$ (8 mg, 0.01 mmol) and sodium ascorbate (16 mg, 0.03 mmol) were separately dissolved in degassed $H_2O$ (3 mL) and added to the reaction mixture. The reaction mixture was vigorously stirred for 16 h at rt and subsequently evaporated to dryness. The crude residue was purified by flash column chromatography (DCM → 2% MeOH in DCM), followed by further HPLC purification to afford title compound (3.2 mg, 3 µmol, 5%). HRMS (ESI+) m/z: calculated for $[C_{57}H_{63}ClN_{11}O_4]^+$: 1000.47475; found: 1000.47462. $^1$H NMR (600 MHz, MeOD): δ 8.20 (td, *J* = 13.1, 3.2 Hz, 2H), 8.07 (s, 1H), 7.97 (s, 1H), 7.66 (dd, *J* = 7.8, 1.7 Hz, 1H), 7.52 – 7.45 (m, 4H), 7.39 (m, 2H), 7.35 – 7.30 (m, 2H), 7.29 (d, *J* = 7.1 Hz, 1H), 7.27 – 7.22 (m, 4H), 6.66 (d, *J* = 2.6 Hz, 1H), 6.57 (t, *J* = 12.4 Hz, 1H), 6.51 – 6.42 (m, 2H), 6.40 (dd, *J* = 16.9, 1.9 Hz, 1H), 6.27 (d, *J* = 13.7 Hz, 1H), 6.19 (d, *J* = 13.7 Hz, 1H), 5.82 (dd, *J* = 9.9, 1.9 Hz, 1H), 5.17 (s, 2H), 4.57 – 4.53 (m, 2H), 4.04 (t, *J* = 7.7 Hz, 2H), 3.77 (s, 3H), 3.69 – 3.65 (m, 2H), 3.56 (s, 3H), 2.66 (s, 3H), 2.15 (t, *J* = 7.2 Hz, 2H), 1.79 – 1.58 (m, 18H), 1.40 – 1.31 (m, 2H). $^{13}$C NMR (151 MHz, MeOD): δ 176.2, 175.3, 174.6, 166.9, 155.4, 145.1, 144.2, 143.5, 142.6, 142.5, 133.0, 131.4, 129.8, 129.7, 129.0, 128.7, 128.1, 127.4, 126.6, 126.2, 126.2, 126.0, 125.6, 123.4, 123.2, 112.0, 111.8, 106.8, 104.3, 104.2, 100.7, 62.8, 56.4, 49.8, 44.8, 40.3, 36.5, 28.1, 27.9, 27.9, 27.8, 27.1, 26.4.
